# Inferring the causes of animal social network structure from time-series data

**DOI:** 10.1101/2025.11.08.687336

**Authors:** Ben Kawam, Richard McElreath, Julia Ostner, Daniel Redhead, Oliver Schülke

## Abstract

Behavioural ecologists aim to understand the causes of animal social structure. Connecting theoretical models of social structure with empirical observations remains, however, a formidable challenge. While most of the current statistical methods for animal social network analysis rely on data that are aggregated over time and summarised as one behavioural dimension (*e.g*., an adjacency-matrix), common behavioural sampling techniques (*e.g*., focal-animal sampling) produce data in continuous time, and involve different behaviours. Furthermore, empiricists in the field are generally interested in causal inference, but lack a framework to rigorously analyse focal-animal sampling data in light of transparent causal assumptions. As a consequence, common methods are often inappropriate, and can lead to wrong biological conclusions. Here, we introduce a causal Bayesian modelling framework to empirically study the causes of social network structure from focal-animal sampling data. We start by outlining a *generative model* that encodes how biological and measurement processes jointly produce social network data in continuous time; namely, as a temporal sequence of dyadic behavioural states (*e.g*., no body contact, social resting, grooming). Building upon the generative model, we develop a *statistical model*: a multilevel, multiplex Bayesian model that takes raw focal observations as input, and produces a posterior probability distribution for the generative parameters as output. After validating the statistical model’s performance with sparse data— common in real-world settings—we illustrate its application with an empirical data set collected in wild Assamese macaques. We notably showcase how researchers can compute probabilistic estimates for well-defined causal hypotheses about the drivers of social structure. With this work, we not only contribute novel theoretical and statistical tools to the field, but also illustrate a *workflow* that allows researchers to iteratively translate their domain expertise into a formal analytical strategy—bridging theoretical and empirical research in behavioural ecology.

## 1. Introduction

Behavioural ecologists seek to understand the ecological and evolutionary drivers of animal behaviour. When studying sociality, they may ask: what drives two primates, whether human or non-human, to develop a long-lasting social bond (Redhead et al., 2023a; Seyfarth & Cheney, 2012)? Why do certain species of lizard engage in costly fights (Whiting & Miles, 2019)? What makes vampire bats regurgitate hard-earned blood meals to their conspecifics (Davies et al., 2012)? Such questions, where one focuses on a behavioural outcome and assesses the importance of its causes, can be thought of as a form of *reverse* causal reasoning (Gelman, 2011; Gelman & Imbens, 2013): one starts from the consequence—here, a behavioural phenomenon—and traces it back to its multiple causes.

Behavioural ecologists approach such questions by developing theoretical models—whether verbal (*e*.*g*., Hinde, 1976; Silk, 2002) or formal (*e*.*g*., Seyfarth, 1977; Smaldino, 2023)—that posit how several factors could jointly produce the behaviour of interest. From these models, they can focus on particular mechanisms to be investigated in detail. One might, for instance, hypothesise that the kinship of two individual primates influences the strength of their social relationship. One may further suppose that resource availability shapes competition among lizards (Whiting & Miles, 2019), or that dyadic reciprocity drives food sharing in vampire bats (Davies et al., 2012). When narrowing down their inquiry to one plausible causal mechanism, researchers shift to another form of causal reasoning: so-called *forward* causal reasoning (Gelman, 2011; Gelman & Imbens, 2013). Such reasoning underpins most empirical studies in behavioural ecology (Davies et al., 2012; Shipley, 2016; Tinbergen, 1963), whether experimental or observational—although in the latter case, the causal nature of the results is sometimes left ambiguous (Grosz et al., 2020; Kawam et al., 2025; Pearl & Mackenzie, 2018).

To assess the empirical support for their causal hypotheses, empiricists collect behavioural data using standardised methods like *focal-animal sampling* (Altmann, 1974). This common procedure involves selecting an individual animal at random, and recording their behavioural actions for a predetermined amount of time. That is, a field worker focuses on an animal and follows what they are doing for, say, 40 minutes. Focal-animal sampling generates data sets in *continuous time*, where each row corresponds to a behavioural action that is, at the minimum, characterised by an actor and— for social behaviours—a receiver. We illustrate what these data typically look like in Figure 1. This procedure is sometimes abbreviated as a “focal-animal follow”; accordingly, in the remainder of this manuscript, we also refer to the focal-animal sampling procedure as a *focal-follow*, to the protocols of fixed duration as *focal-follow protocols*, and to the resulting data as *focal-follow data*.

**Figure 1.**
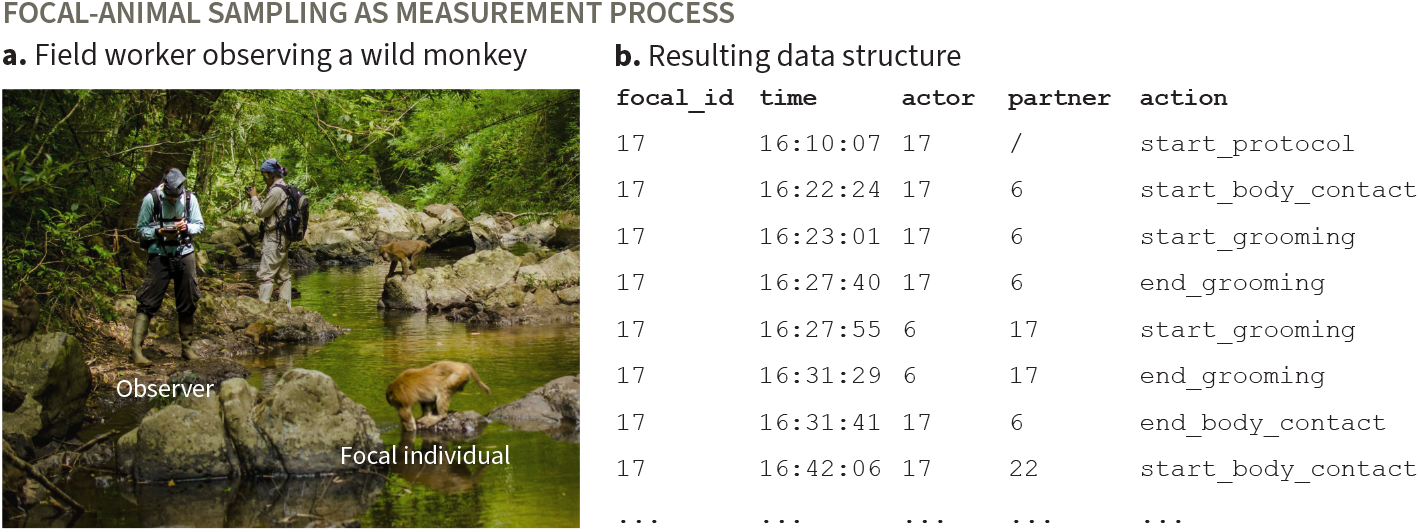
Focal-animal sampling constitutes a standard behavioural data collection method. (**a**.) A field worker conducts a focal-follow protocol on a wild Assamese macaque (*Macaca assamensis*) by recording its behaviour continuously for a pre-determined amount of time. (**b**.) Reduced example of data frame resulting from a focal-follow protocol, focusing exclusively on social behavioural events. The numbers in the first, third, and fourth column refer to individual animals.

The core inferential challenge that we address in this manuscript can be stated simply: how can one assess causal hypotheses about social behaviour from focal-follow data (Figure 1)? Behavioural ecologists are generally taught that to study causality, one needs to experimentally *manipulate* the variable, or experimental “treatment” of interest (Shipley, 2016). However, randomised controlled experiments do not always generalise to the natural populations of interest (Cronin et al., 2017; Dal Monte et al., 2022; Hagen & Hammerstein, 2006; Redhead & Power, 2022). Moreover, experimental manipulation of wild animals is often impossible for ethical and/or practical reasons. Consequently, the subfield of behavioural ecology that focuses on animal societies—*i*.*e*., animal social network analysis—is a largely observational field, where causal evidence must be built by studying the compatibility of causal models of the world with observed data like those generated by focal-animal sampling (Kawam et al., 2025). In this context, researchers are left with only one option: the examination of their causal assumptions, and the careful connection between these assumptions and their data.

Evaluating the empirical support for a causal hypothesis of interest poses a serious challenge when relying primarily on observational behavioural data. One important reason is that the measurement process through which researchers acquire behavioural data is markedly different from those described in the conventional statistics textbooks that are typically used to train behavioural ecologists. To explain the observed variation in a dataset, one typically learns to use a Gaussian linear model if the response variable is continuous; a Bernoulli generalised linear model if the outcome is binary; and so on—and such models are generally fitted using popular R packages like *brms* (Bürkner, 2017) or *lme4* (Bates et al., 2015). However, where does one even start when confronted with time-series data like that shown in Figure 1**b**?

The most common analytical methods in the field involve aggregating behavioural data in such a way that they can be directly fitted to off-the-shelf statistical models. Researchers usually start by splitting their data sets into arbitrarily defined time windows (*e*.*g*., a year), over which the variables of interest are aggregated. For instance, one may record whether two individuals have interacted in a certain way (*e*.*g*., body contact, grooming, fighting) over this period, or count the number of times that they have done so, to fill the entries of an adjacency-matrix. In the case of behavioural counts, one may further divide this number by the amount of time that the dyad was observed for, computing a *Simple Ratio Index* (SRI) in an attempt to control for variation in observation effort (Farine & Whitehead, 2015). Sometimes researchers do so, not for one behaviour only, but for several of them, and average over these behaviours to construct composite indices (*e*.*g*., DSI and related indices; see Silk et al., 2013). Either way, the resulting adjacency-matrix typically operationalises the social network under study (Krause et al., 2015; Whitehead, 1997). Once the edges of the social network have been defined, researchers may further aggregate their already-aggregated data, by computing social network metrics at the node-level (*e*.*g*., degree, strength) or at the network-level (*e*.*g*., density), and explain the variation in these metrics using multiple regression and/or network permutation (Farine, 2017; Farine & Carter, 2022; Farine & Whitehead, 2015). In short, instead of tailoring a statistical model to the data and the process of interest, these practices consist in reshaping the data so as to fit off-the-shelf models.

However, these popular procedures carry fundamental flaws, many of which have been well documented in applied statistics and network science, as well as, more recently, in animal social network analysis (Redhead et al., 2025). First of all, aggregating data over pre-defined time windows discards the fine-grained information characterising specific behavioural observations (*e*.*g*., observer identity, age, temporary dominance rank). As a result, researchers are either unable to incorporate this information into their analyses or are forced to make arbitrary—and potentially deleterious— simplifications; for example, by assuming that an individual’s rank on day one remains constant throughout the year. Subsequently aggregating social network into SRI and related indices further amounts to ignoring the inferential *uncertainty* about the underlying rates of dyadic interactions. The true rates of behavioural events (*e*.*g*., grooming, aggression) are typically unknown, and should be integrated into a statistical model in such a way that more behavioural data results in more certainty in the estimated rates (Bonnell et al., 2024; Duboscq et al., 2023; Hart et al., 2023; Redhead et al., 2023b; Ross et al., 2023; Sosa et al., 2025; Young et al., 2020). Social network data are usually sparse, in that typically, not every dyad is observed interacting many times over the course of a study period, making the problem particularly potent. Composite indices (*e*.*g*., DSI) have similar precision issues (Mielke & Samuni, 2021), and can additionally suffer from construct invalidity (McKenna & Heaney, 2020; Ravallion, 2012). Regardless of the technique used to construct social network edges, the variation in the resulting network metrics (*e*.*g*., node strength) is then typically explained by statistical models that are not derived from formal causal assumptions. Without such assumptions, there is no principled way to address *confounding* (Byrnes & Dee, 2024; Cinelli et al., 2022; Franks et al., 2025; Pearl, 2009; Pearl et al., 2016; Siegel & Dee, 2025) or *missing data* (Mohan, 2022)—both of which plague observational ecology at large, with animal social networks being no exception. Generally, the field lacks a transparent workflow that logically connects causal models of social networks to empirical data (Deffner et al., 2024; Gelman et al., 2020; Kawam et al., 2025). In fact, from a theoretical standpoint, many analyses of animal social network data cannot do what they are intended to do, even in principle—as exemplified by the common use of permutation tests to automatically handle confounding (Hart et al., 2022; Weiss et al., 2021), or the use of predictive criteria (*e*.*g*., AIC) to select models for causal inference (Arif & MacNeil, 2022). As a result, many studies in the field stand on loose methodological ground, which leads to unjustified, and often wrong conclusions. We do not have the space to expand on these problems here, but we redirect interested readers to Kawam et al. (2025).

Although the challenges of animal social network analysis are specific to the field, they share important commonalities with other areas of ecology. Community ecologists aim to uncover interaction networks between species from noisy, and potentially biased observations (Banville et al., 2024; Young et al., 2021); movement ecologists infer latent behavioural processes and individual differences from spatio-temporal recordings (Auger-Méthé et al., 2021; Leos-Barajas et al., 2017b, 2017a); and population ecologists work with incomplete observations to estimate true population sizes (Hilborn & Mangel, 1997). Like social network data, these observations are often noisy, biased, subject to confounding, and—*e*.*g*., in the case of movement data—often collected as temporally resolved observations. Because of their data structure, they are unsuited to be directly fitted to general-purpose linear models. In all these fields, researchers have addressed their respective inferential challenges by developing specifically-tailored models that link assumed data-generating mechanisms and empirical observations—a tradition with decades-long roots in some cases (*e*.*g*., Chapman & Robson, 1960; Jackson, 1939). Somewhat surprisingly, animal social network analysis still lacks a principled modelling framework for its core data types, including focal-follow data.

Here, we introduce a causal Bayesian modelling framework to empirically study the causes of social network structure from focal-animal sampling data. We start by outlining a generative model that describes how biological and measurement processes jointly produce such data. At its core, it is a Markov process that governs how pairs of individuals sojourn (*i*.*e*., stay) in, and transition between behavioural states; *e*.*g*., no body contact, social resting, grooming. We present a quantitative version of this model—a simulation that can be used to generate data structured like those obtained from focal-follows—, along with a qualitative representation using causal diagrams (Pearl, 2009; Pearl et al., 2016). Subsequently, we develop a statistical model that mirrors the generative model, and is capable of recovering its parameters using Bayesian inference. While distinct, our statistical model draws inspiration from prior work in social network analysis (Kenny & La Voie, 1984; Li & Loken, 2002; Redhead et al., 2023b; Ross et al., 2025; Shalizi & Thomas, 2011; Snijders & Kenny, 1999; Van Duijn et al., 2004; Wasserman & Faust, 1994), statistics (Gelman & Shalizi, 2013; Gelman et al., 2021; Kass, 2011; McElreath, 2020), and behavioural ecology (Adriaense et al., 2024; Hart et al., 2023; Ross et al., 2023). We describe the model’s joint posterior distribution, and show how formally defined causal estimands can be computed, yielding a probability distribution over clear causal questions (Chatton & Rohrer, 2024; Lundberg et al., 2021). Generally, we embed these components into a *modelling workflow* (Gelman et al., 2020; McElreath, 2020) that helps behavioural ecologists logically connect their domain expertise, research questions, and data, through incremental steps.

This manuscript extends our earlier framework for causal inference in animal social networks (see Kawam et al., 2025) by making several key contributions. To start, we model social interactions as they unfold over continuous time, rather than aggregating them into pre-defined time windows. This allows us to more closely approximate the empirical processes generating animal social network data—redefining what constitutes an “animal social network” in practice. In this respect, our framework aligns with some of the ideas underlying relational event modelling (Bianchi et al., 2024; Stadtfeld & Block, 2017) and stochastic actor-oriented models (Snijders, 2017). Furthermore, our approach allows researchers to study multiple aspects of social relationships jointly—specifically, the duration of several behavioural states (*e*.*g*., time spent grooming, interaction frequency), and the transitions between them (*e*.*g*., immediate reciprocation of grooming). We do so while retaining fine-grained information for each data point, laying the groundwork to rigorously study how social behaviours unfold, and affect one another over time.

In addition, while all of our models build upon existing methodology, we may nonetheless emphasise some particularly interesting technical aspects that are novel to animal social network analysis and, in several cases, to social network analysis more generally: (i) we use plated causal graphs to represent the multilevel causal structure of social network data-generating processes—providing tools to study group-, individual-, dyad-, and observation-level causal effects (see Koller & Friedman, 2009; Weinstein & Blei, 2024); (ii) we develop a cross-level varying-effects structure to effectively estimate generative parameters even if little or no data are available for certain units; (iii) we define a time-to-event observation model that deals with the censoring that is inherent to focal-follows—helping us, *e*.*g*., to model grooming bouts whose start or end are unknown; (iv) we model hidden behavioural sub-states with a mixture likelihood—enabling us to probabilistically assess, *e*.*g*., whether two individuals are not interacting because they are far away from one another, or because of a short break between two interactions; (v) we illustrate how posterior predictive checks can guide a rigorous and intuitive analytical workflow for causal inference in animal social network analysis.

In the following section of the manuscript, we start by highlighting a set of *core* models that constitute the scaffolding of our framework. We deploy these models in the context of a case study of wild Assamese macaques (*Macaca assamensis*). Then, we showcase how our models can be extended to study the causal effect of a simple individual-level feature, sex, on social behaviour. Our goal is not to study the behaviour of Cercopithecinae monkeys, nor to propose a one-size-fits-all model that could be applied to any species. Rather, we emphasise the causal structure underlying social network data and how it can be mapped onto statistical models. Empiricists can then build upon this foundation by tailoring the models to their study system and inferential goals.

## 2. Core models

### 2.1. Generative model

The first step to meaningfully analyse social network data is to think of the mechanisms that could have generated the networks in the first place, and to translate these mechanisms into formal assumptions. These assumptions might be more or less credible—after all, certain aspects of a researcher’s knowledge will unavoidably be uncertain, or incomplete. Even so, transparent assumptions are necessary, for they provide a licence for an analytical strategy (Pearl & Bareinboim, 2014; Woensdregt et al., 2024). Formal assumptions allow researchers to *validate* that their statistical approach works under known conditions, and helps them *assess* whether these conditions are biologically plausible. Put differently: unless one is confident that they can interpret complex statistical patterns that have been generated from known (assumed) processes, it is unreasonable to make sense of such patterns generated from unknown empirical mechanisms (Smaldino, 2017, 2023).

In our case, we aim to model the generating mechanisms underlying focal-follow data (Figure 1).

The key idea to model these data directly is to consider that each row in Figure 1**b** corresponds to a change in the behavioural *state* of a dyad, where a state is a type of activity that a dyad performs for a certain amount of time (*e*.*g*., grooming, social resting, no body contact, play). Let us for instance consider the second and third rows of panel **b**: the row stating that “individual 17 and individual 6 start body contact”, followed by “17 starts grooming 6”, reflect a transition between two distinct dyadic states: direct contact without grooming, and grooming. The number of possible behavioural states will thus depend on the level of detail at which behavioural actions were recorded.

In the following section, we turn to defining a simple generative model that can generate such states. We tried to explain our mathematical notation, and to give intuitive examples as much as we could. If they remain too opaque, we strongly suggest McElreath (2020) for a general introduction to generative thinking and Bayesian inference. Kawam et al. (2025) covers these topics in the context of animal social networks.

#### 2.1.1. Markov process

We assume that there is a number *K* of possible behavioural states that describe what a dyad is doing over time (Figure 2). These states form what is called the *state-space* of our generative model.

**Figure 2.**
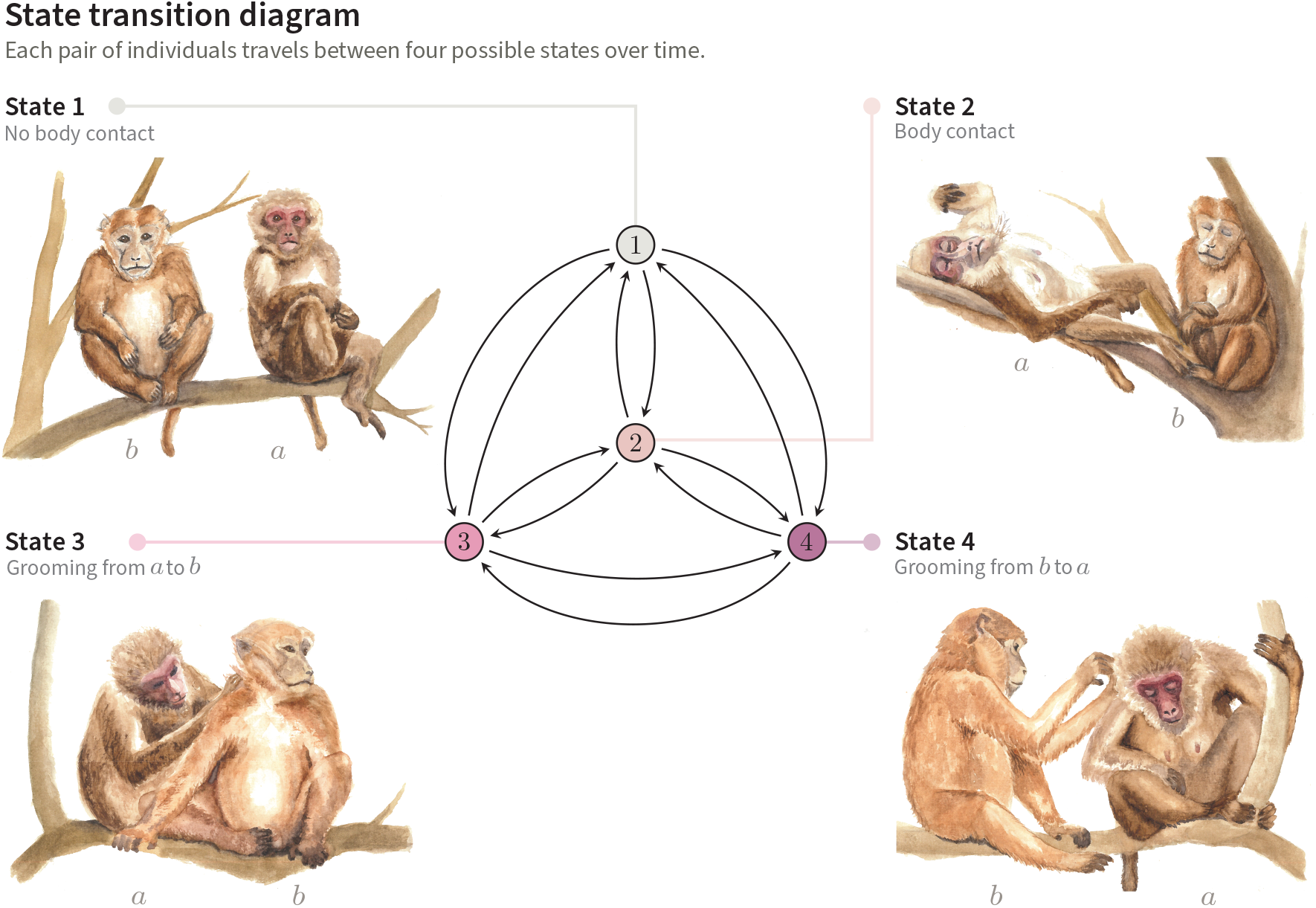
State transition diagram of our generative model. Each dyad (*a, b*) moves through four discrete states over time, represented by coloured circles. The dyad remains in a given state for a certain duration, or *holding time*, before transitioning to a new state according to state-specific *transition probabilities*, indicated by arrows showing all possible (non-zero) transitions. Paintings by Sofia M. Pereira & Judith von Nordheim.

Each dyad is labelled by a unique pair of individuals, (*a, b*), where:

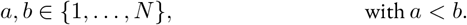

That is, *a* and *b* are individual identifiers ranging from 1 to *N*, where *N* is the total number of individuals under study. To avoid duplicates, we write *a* < *b* so that each pair appears only once— *e.g*., we use (1, 2) but not (2, 1). For example, if *N* = 3, the unique dyads are: (1, 2), (1, 3), and (2, 3).

In the examples used throughout this manuscript, we consider *K* = 4 dyadic states. These are: (1) no body contact between two individuals; (2) body contact without grooming; (3) grooming from *a* to *b*; (4) grooming from *b* to *a*. Note that we follow standard notation when possible, *e.g*. we use lowercase letters like *k* to refer to specific states (*e.g*., state *k* = 3), and uppercase letters like *K* to refer to the total number of states.

Two key ingredients determine how a dyad moves through the state-space introduced above: how long it stays in a given state, and the probability of switching to the possible new states once that time is up. These processes take place sequentially. First, a dyad (*a, b*) enters a state—say, body contact, or *k* = 2—and stays there for some amount of time. A stay in a given state is called a *sojourn*. The duration of the sojourn is called the *holding time*. Once this time is over, the dyad transitions to a new state *l* ∈ { 1, 2, 3, 4}, where *l* ≠ *k*. That is, the dyad transitions from its current state (called *k*) to a new state (called *l*).

These dynamics are controlled by two fixed sets of parameters that are different for each dyad. First, an array *θ* (theta) for each pair (*a, b*), *θ*_[*a,b*]_, determines the average holding time in each of all possible states (Table 1):

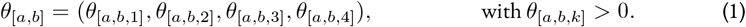

**Table 1.** Key variables and their definition.

| Name | Domain & constraints | Indexed by | Definition | Example |
| --- | --- | --- | --- | --- |
| $\theta$<br>(theta) | $> 0$ | State $k$ of a dyad $(a, b)$<br>$\theta_{[a,b,k]}$ | Average duration of a sojourn (= average *holding time*). | $\theta_{[1,2,3]}$ is the average duration of a grooming bout (state 3) from individual 1 to individual 2. |
| $\Gamma$<br>(gamma) | $\sum_l \Gamma_{[a,b,k,l]} = 1$ | Current state $k$ and potential next state $l$, for a dyad $(a, b)$<br>$\Gamma_{[a,b,k,l]}$ | Transition probability from current state to future state. | $\Gamma_{[6,12,3,4]}$ is the probability that individual 12 directly reciprocates grooming to 6 following a grooming from 6 to 12 (state 3 to state 4). |
| $U$ | $\mathbb{R}$ | Group $g$, ind. $a$ or $b$, or dyad $(a, b)$<br>$U_{[g]}^{\text{gr}}, U_{[a]}^{\text{ind}}, U_{[b]}^{\text{ind}}, U_{[a,b]}^{\text{dyad}}$ | Unobserved cause of $\theta$ and $\Gamma$. | $U_{[1]}^{\text{ind}}$ is the "gregariousness" of individual 1, affecting what states it sojourns in with others through $\Gamma_{[1,b,k,l]}$, and how long for, through $\theta_{[1,b,k]}$. |
| $S$ | $\{1, \dots, K\}$ | Observation $j$ of a dyad $(a, b)$<br>$S_{[a,b,j]}$ | State number. | $S_{[1,2,36]}$ is the dyadic state that the (1, 2) dyad was in, for observation 36. |
| $X^{\text{true}}$ | $> 0$ | Observation $j$ of a dyad $(a, b)$<br>$X_{[a,b,j]}^{\text{true}}$ | True sojourn duration. | $X_{[1,2,36]}^{\text{true}}$ is the true duration of the 36th observation for individuals (1, 2). |
| $X$ | $> 0$ | Observation $j$ of a dyad $(a, b)$<br>$X_{[a,b,j]}$ | Observed sojourn duration. | $X_{[1,2,36]}$ is the recorded duration of the 36th observation of individuals 1 and 2. |
| $C$ | $\{0, 1\}$ | Observation $j$ of a dyad $(a, b)$<br>$C_{[a,b,j]}$ | Indicates whether an observation was censored ($C = 1$) or not ($C = 0$). | $C_{[1,2,36]}$ describes whether the 36th observation for the (1, 2) dyad was censored, or if instead, its ending time is known. |
| $Q$ | $\{0, 1\}$ | Observation $j$ of a dyad $(a, b)$<br>$Q_{[a,b,j]}$ | Indicates whether a sojourn in state $k = 1$ was short ($Q = 0$) or long ($Q = 1$). | Given that $S_{[1,2,36]} = 1$, $Q_{[1,2,36]} = 1$ indicates that the sojourn captured by the 36th observation for the (1, 2) dyad is a long sub-state in $k = 1$. |
| $\delta$<br>(delta) | $0 \text{ min} < \delta < 60 \text{ min}$ | Fixed across dyads | Average duration of a short sojourn in state 1. | $\delta = 5.0$ means that all dyads spend, on average, 5 minutes in the short sub-state 1. |

- i. $Q$ and $\delta$ only apply to the extended generative model (section 3.1). - ii. In this table, as well as in the rest of the manuscript, we leave the nestedness of social groups' subunits implicit. That is, to represent variables defined over units within a social group $g$ (e.g., $\theta_{[g,a,b,k]}$ ), we skip the $g$ in the index for readability (e.g., $\theta_{[a,b,k]}$ ). - iii. Although observations $j$ are nested within dyads $(a, b)$ , we number $j$ from 1 to $J$ , the total number of observations across dyads. For example, if $j = 1$ is a sojourn in state $k = 1$ for dyad (14, 15), and $j = 2$ is in the same state for dyad (14, 21), these will be written as $S_{[14,15,1]} = 1$ and $S_{[14,21,2]} = 1$ . - iv. $\mathbb{R}$ denotes the set of real numbers, indicating that $U$ are continuous, unbounded variables (e.g., z-scores).

Here, each *θ*_[*a,b,k*]_ represents the average duration that (*a, b*) spends in state *k*. It means that, for instance, the average holding times that the dyad (1, 2) spends in the behavioural states is given by a vector of size *K*:

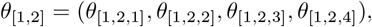

Where each *θ*_[*a,b,k*]_ corresponds to a positive real number (*e.g*., 5.3 minutes). The realised holding times are then random draws from an exponential distribution with the specified mean *θ* (Figure 3**a**). The exponential distribution was chosen because it is a conservative and convenient^1^ time-to-event distribution (McElreath, 2020).

**Figure 3.**
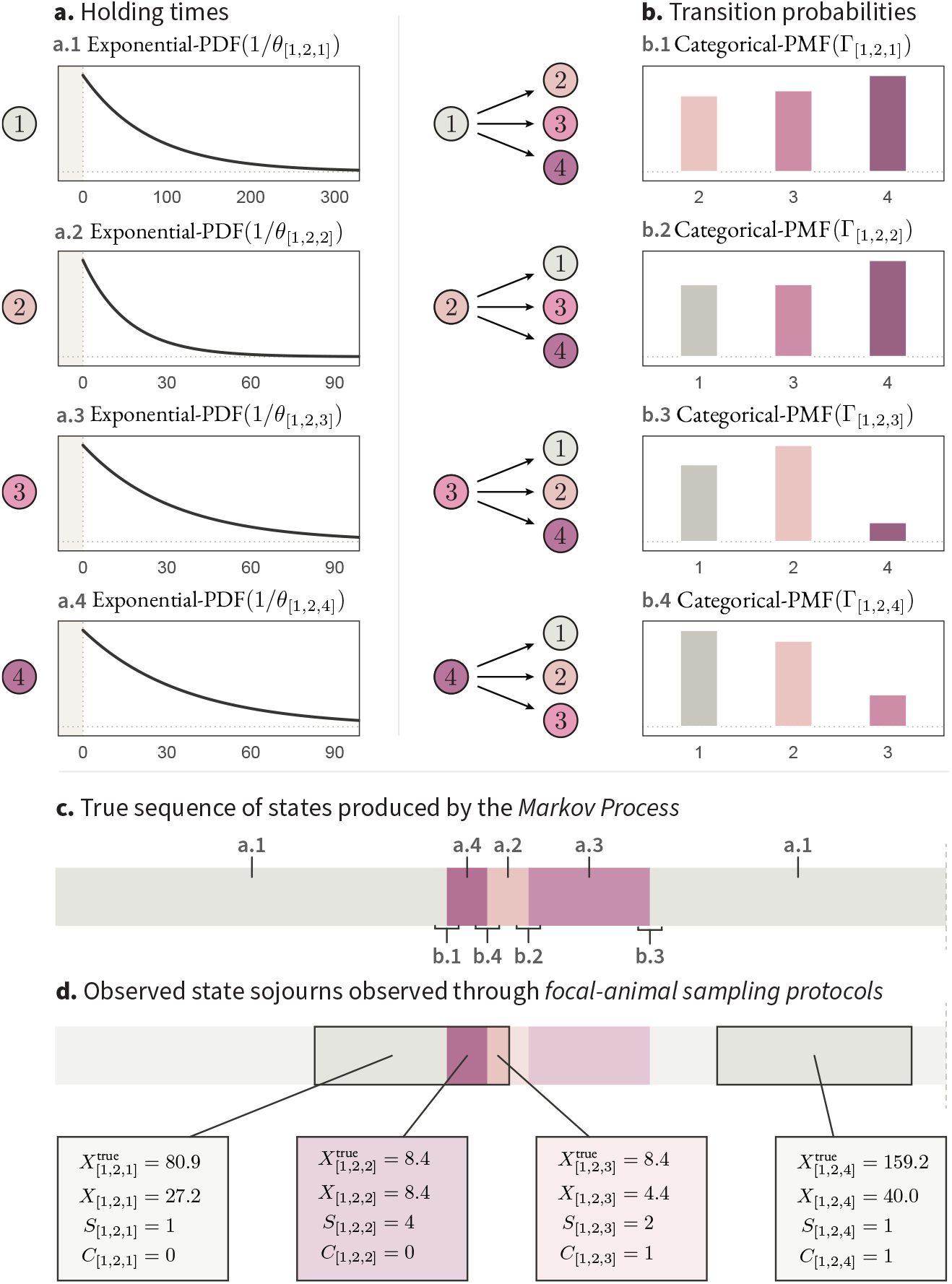
Parameters and distributions of the generative model for an example dyad, (1, 2), along with the resulting sequences of true and observed state sojourns. (**a**.) Probability Density Functions (PDFs) from which holding times are drawn for the dyad under scrutiny. Notice that the x-axis’ scale differs across states *k*. The parameters *θ*_[1,2,*k*]_ are the true average holding times in the corresponding states *k*, and they govern the shape of the PDFs. (**b**.) Probability Mass Functions (PMFs) from which state transitions are drawn. For each current state *k*, Γ_[1,2,*k*]_ is a vector of probabilities controlling how likely the dyad is to transition to each possible next state—a higher probability corresponds to a higher bar. (**c**.) Sequence of true state sojourns generated using the Markov process. The x-axis indicates time, moving left to right, with colours corresponding to the different states. The distribution from which the sojourns and transitions were sampled are indicated by the labels. (**d**.) Sequences of observed sojourns *j* generated by focal-follows. The true holding time *X*^true^, observed holding times *X*, states *S*, and censoring levels *C* are shown for four such observations, across two focal-follow protocols represented by black rectangles overlapping the state sequence.

Second, an array **Γ**—capital gamma—for each dyad (*a, b*), **Γ**_[*a,b*]_, describes the transition probabilities between states (Table 1):

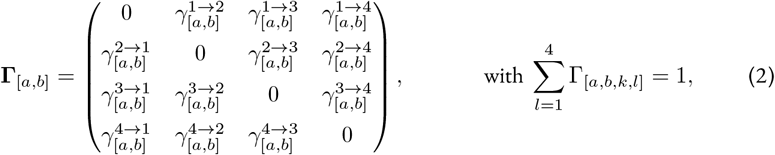

Each parameter 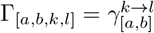 gives the probability of transitioning from state *k* to state *l*, with *γ* denoting lowercase gamma. In other words, each dyad is characterised by a *K* × *K* matrix controlling its transition dynamics. If a dyad (*a, b*) is currently in state *k*, the probabilities of transitioning to any other state are given by the *k*^th^ row of the matrix, denoted Γ_[*a,b,k*]_ (Figure 3**b**). For instance, the probability that the dyad (1, 2) transitions from state 4 to the next states is given by:

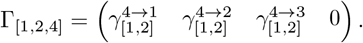

Such a probability mass function is shown as an example on Figure 3**b2**. On somewhat technical matters, note that the diagonal of **Γ**_[*a,b*]_ is filled with 0s, to encode that a dyad cannot transition from a state *k* to a state *l* = *k*; if a dyad remains in the same state, their prolonged sojourn is counted as part of the same holding time. Furthermore, to ensure the model behaves properly, the probabilities in each row must sum to 1. This guarantees that Γ_[*a,b,k*]_ forms a valid categorical—*i.e*., multinomial— distribution over the possible next states (Blitzstein & Hwang, 2019).

Lastly, notice that the assumed dynamics—and stability—of our model is already reflected by the generative parameters **Γ** and *θ*. First, the transitions of a dyad (*a, b*) between behavioural states are governed by its matrix **Γ**_[*a,b*]_ *only*; specifically, by the *k*^th^ of **Γ**_[*a,b*]_, with *k* corresponding to its current state. Because the probability of transitioning to a new state solely depends on the current state *k*, and not on past states, this setup is known as a *Markov Model* (Ross, 2014). The resulting random sequence of behavioural states over time is referred to as a *Markov process* or *Markov chain* (Box 1). Second, by defining a fixed set of parameters *θ* and **Γ** per dyad (*a, b*), we assume that these parameters are constant, and not affected by time-varying causes like age or season. While these two assumptions are likely unrealistic for most species and behaviours, they serve as a pragmatic starting point, to be relaxed in future extensions (see Discussion).

With this nomenclature in place, the process by which dyads interact over time can be summarised as follows (Figure 3**a-c**):

0 The dyad (*a, b*) begins in an arbitrary initial state *k*; *e.g*., *k* = 1.
1 It remains in this state for a holding time drawn from an Exponential distribution with mean *θ*_[*a,b,k*]_.
2 It transitions to a new state *l*, drawn from a categorical distribution with probabilities Γ_[*a,b,k,l*]_.
3 Repeat (1)–(2) until a fixed runtime threshold is reached.

The Markov process at the core of our generative model is described in more detail in section S1.1.2 and implemented in *R* (version 4.4.3; R Core Team, 2022). The full source code is publicly available in our GitHub repository.

We ran this model for four groups of ten synthetic individuals (*N* = 40), simulating a total of approximately 266 hours of elapsed simulation time. Parameter values were tuned to obtain biologically plausible outcome distributions (Figure S2). The resulting sequence represents the “true” series of social behaviours exchanged among individuals, including when no observer is present to observe them. We represent a simplified output for one dyad in Figure 3**c**. We can think of this process as representing the “biological” component of the data-generating mechanism. However, to structurally match real focal-follow data, one crucial ingredient is still missing: the *focal-follows* themselves—that is, the measurement process.

#### 2.1.2. Measurement process

The measurement procedure can be thought of as an additional layer applied on top of the true sequence of behavioural states generated with the Markov process (Figure 3**d**). By conducting focal-follows, field observers practically cut out windows of continuous observation time from the— otherwise unobserved—behavioural sequence. These windows constitute the observed focal-follow protocols represented in Figure 1**b**.

Before describing how to go from a true sequence of behavioural states (Figure 3**c**) to focal-follow data (Figure 3**d**), let us shortly highlight the relationship between focal-animal follows and dyadic observation effort. While observers focus on one individual *a* at a time when they conduct a focal-follow protocol, it is important to realise that, in doing so, they practically follow several dyads. They follow all of the dyads that *a* is a part of, *i.e*., *N* − 1 pairs. Suppose, for instance, that there are *N* = 5 individuals under study. Following individual 1 for 40 minutes corresponds to observing all of the potential interactions that 1 might exchange with 2, 3, 4, and 5 during this time. This results in a total of (*N* − 1) · 40 = 160 minutes of effective dyadic observation time.

In our simulations, we assumed that each social group was observed by a single observer, that focal animals were selected independently of their behaviour, that sampling protocols had a fixed duration, and that these protocols occurred consecutively in time. For example, the synthetic observer would pick an individual at random—regardless of what it is doing—, observe its behaviour for 40 minutes, thereby observing *N* − 1 dyads for 40 minutes each. Following the end of this protocol, the synthetic observer would then immediately select another individual to follow, and so on. These assumptions generate random sampling-windows layered on top of the true sequence of states, like those shown in Figure 3**d**. We provide more details about the measurement procedure in section S1.1.3, and on our GitHub repository. Note that although our generative model assumes uninterrupted 40-minute focal protocols, empirical protocols may be interrupted by focal-animal loss, particularly for certain behaviours; we therefore encourage researchers to relax these assumptions when necessary.

##### Box 1 Clarifying equivocal jargon

*Generative model*. A model capable of generating data. This includes causal models, like the model described in section 2.1, but sometimes, purely statistical devices as well—*e.g*., Bayesian statistical models can be considered “generative” because they allow for prior and posterior predictions. In this manuscript, however, we use the term “generative model” for our causal models only, as they explicitly encode the different layers of the data-generating mechanism.

*Markov process*. A stochastic process in which the next state depends only on the current state, and not on the states that preceded it. Here, we use Markov processes in two contexts. First, to describe the sequence of dyadic behavioural states of our generative model (a discrete state space; see Figure 2). Second, to refer to the sequence of MCMC draws approximating the posterior distribution of our statistical model’s parameters—a continuous state space across the statistical parameters, that themselves capture the generative parameters governing the Markov process for behavioural states.

*Network*. A network is a collection of nodes that are connected by edges, representing the connection between them. In this manuscript, we refer to three types of networks. (i) Social networks, in which nodes represent individual animals and edges denote their social relationships or interactions (Kawam et al., 2025). Although we do use the term “social network” here, we avoid the corresponding—inherently static—graphical representation of social structure. (ii) The state-transition diagram representing the state-space of a Markov model, with nodes corresponding to behavioural states, and directed edges indicating possible transitions between them; see Figure 2. (iii) Causal diagrams, where nodes represent variables and directed edges encode assumed causal relationships between them; *e.g*., Figure 4. In our framework, social structure—typically represented by (i)—is instead operationalised as a dynamic system (ii) with underlying parameters embedded in a multilevel causal structure (iii).

**Figure 4.**
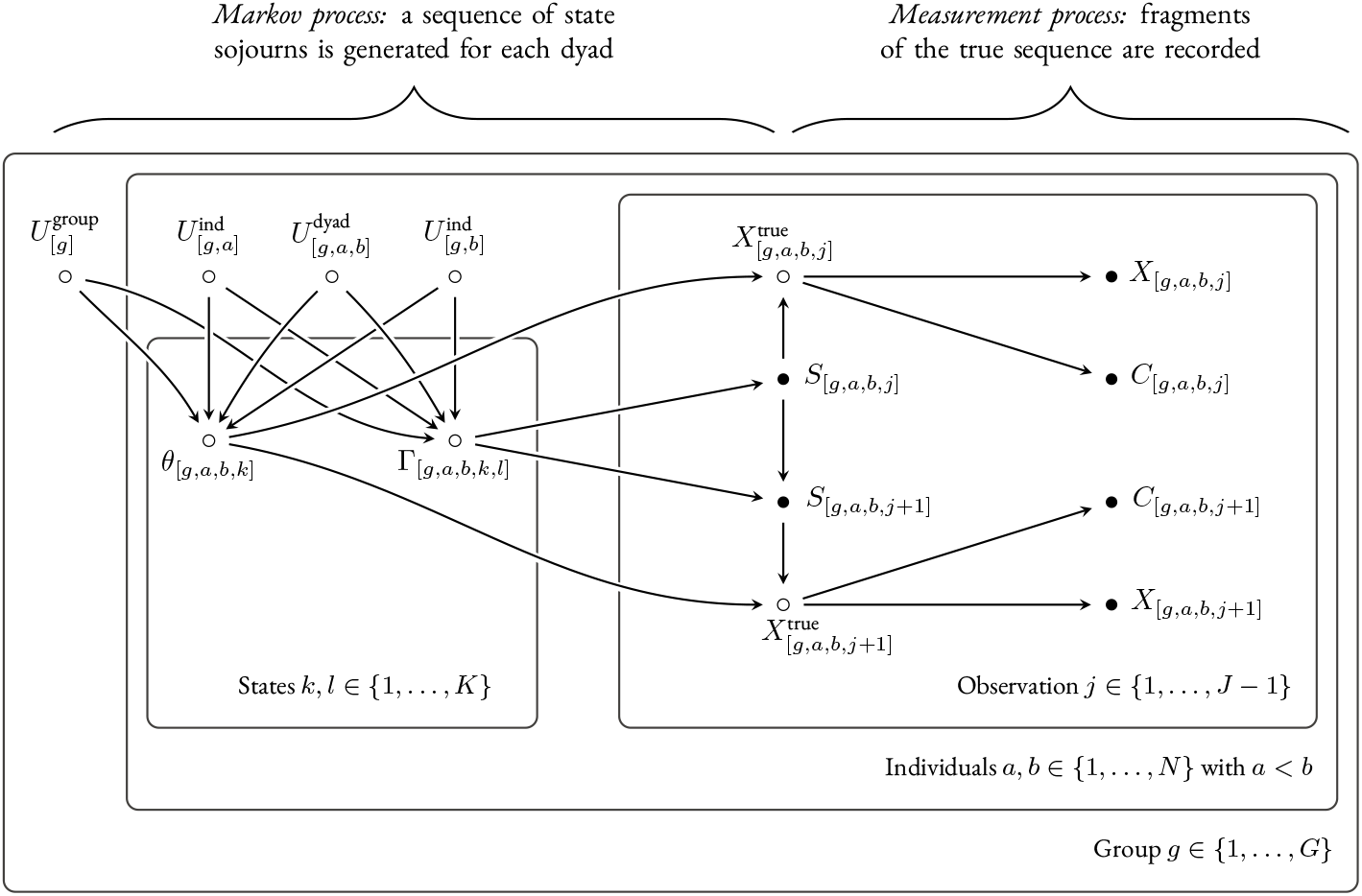
Causal diagram of a social network data-generating process involving several dyadic behavioural states. In this Directed Acyclic Graph, the nodes represent variables, with empty nodes corresponding to unobserved or partially observed variables. The arrows are causal effects, and the *plates*—the rounded-edged rectangles—encode the hierarchical structure of the system. Plates correspond to repeated structures, grouping variables that are replicated over certain indices, shown at their bottom-right. The outermost plate is indexed by group *g*, meaning that variables within that plate are attributes of a group *g*. Variables inside a nested plate are attributes of *sub-units* within *g*. On the second outermost plate, for instance, unordered pairs of individuals (*a, b*) are naturally nested within social groups *g*. Each dyad is further characterised by two sets of parameters: *θ*, describing holding times in each state of the Markov model; and Γ, the probabilities governing how the dyad transitions from one state to the next. These parameters are affected by unobserved group-, individual-, and dyad-level causes *U*, creating important patterns of covariation between them. Moving to the right-hand side of the figure, we show a number of observations *j* for each pair (*a, b*). An observation *j* is a recorded sojourn in a dyadic state. It is characterised by a state number *S*, a generally unobserved true holding time *X*^true^, an observed holding time *X*, and a censoring level *C*. For any pair of sequential observations *j* and *j* + 1 within the same focal-follow protocol, the state of the dyad at *j* + 1 is determined by two factors: the previous state *S* at *j*, and the transition probabilities Γ. We represent this process by drawing arrows from Γ and *S* of *j* to *S* of observation *j* + 1. The arrow from *θ* to *X*^true^ further encodes that *θ* controls *for how long* the dyad stays in the given states. The true holding time, filtered by the sampling procedure, determine the observed holding time *X*, as well as the censoring level *C*—*i.e*., the data—, which we represent by the arrows going from *X*^true^ to *X* and *C*. We refer readers to section S1.1.1 for an introduction to the plate notation, and to section S1.1.2-S1.1.3 for details regarding the Markov process and the measurement process, respectively.

Our assumed joint data-generation process, combining biological and measurement processes, produces a number *J* of observations (Figure 3**d**). An observation *j* ∈ {1, …, *J*} is defined as the recorded stay, or sojourn, of a dyad (*a, b*) in a behavioural state—the equivalent of a coloured block in Figure 3**d**. Each observation is characterised by four variables, which we detail below.

First, 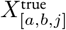 is the true holding time of observation *j*; *i.e*. the true duration of the sojourn of (*a, b*) in a given behavioural state corresponding to observation *j*. 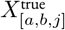 is often unknown in practice: it is especially true for *k* = 1 (no body contact), whose averages *θ*_[*a,b*,1]_ are orders of magnitude higher than the duration of focal-follow protocols. For instance, suppose that a dyad (5, 12) spends on average 2 days in state *k* = 1, but that they are observed for 40 minutes at a time. In most such cases, the 40-minutes protocol will only cover a small fragment of the full holding time. The second variable is *X*_[*a,b,j*]_. It is the *observed* holding time of observation *j*. This duration is equal to 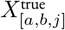 if the sojourn has been fully observed, and is smaller than 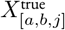 otherwise. Third, *S*_[*a,b,j*]_ ∈ {1, 2, 3, 4} is the behavioural state corresponding to observation *j*. It encodes similar information as *k*, except that we use *k* as an index, and *S* as a variable that characterises observations *j*. Finally, *C*_[*a,b,j*]_ is the censoring state of an observation *j*, where *C*_[*a,b,j*]_ = 0 indicates that the ending time of the sojourn is known (uncensored), and *C*_[*a,b,j*]_ = 1 that it is unknown (censored).

By applying this measurement process to the output of the Markov (biological) process, we obtain a relatively sparse data set: one where many dyads were rarely—if ever—observed in certain behavioural states (Figures S3-S4). For instance, several dyads were not observed grooming at all over the total observation time. Such sparseness is common in empirical social network data sets and, as we will discuss below, represents one of the key inferential challenges that analysts face when designing statistical models.

#### 2.1.3. Causal graph

Before moving on to developing a statistical model, there is one last step we need to take: translating our generative model into a set of qualitative causal assumptions using Directed Acyclic Graphs, or DAGs. DAGs provide a graphical representation of how variables in a system cause one another. Causal graphs are qualitative because they only represent the presence and absence of causal relationships among a set of variables, regardless of their functional forms (*e.g*., whether they are strong, weak, linear, etc.). Despite their apparent simplicity, DAGs are powerful tools. They clarify the causal structure of a system, together with some of the key obstacles to, and corresponding solutions for causal inference (Kawam et al., 2025; Pearl et al., 2016).

In our case, we use a DAG to represent the process that generates social network data (Figure 4). The diagram offers a compact summary of the generative model described above. On a causal graph, an arrow going from a variable *V*_1_ to another variable *V*_2_ indicates that a hypothetical intervention on *V*_1_ would result in a change in the distribution of *V*_2_—a standard definition of “causal effect” in the field of causal inference (Pearl et al., 2016), and the one we use in this manuscript. We also use *plates* (rounded rectangles) to indicate how the variables are repeated across units or levels, making the hierarchical structure of the system explicit. We provide a short introduction to plated graphs in section S1.1.1; see also Koller and Friedman (2009) and Weinstein and Blei (2024).

Below, we briefly unpack our causal graph by discussing, in turn, the Markovian dynamics, the observed holding times, and the causes of *θ* and Γ.

First, the arrows from *S*_[*a,b,j*]_ and Γ_[*a,b,k,l*]_ to *S*_[*a,b,j*+1]_ encode the transition dynamics. Given that *j* and *j* + 1 belong to the same focal-follow protocol (implicit on the graph), the next state *S*_[*a,b,j*+1]_ is sampled from the probability distribution defined by the *k*^th^ row of the transition matrix **Γ**_[*a,b*]_, where *k* is given by the current state *S*_[*a,b,j*]_. In other words, *S*_[*a,b,j*]_ and Γ_[*a,b,k,l*]_ are the two “ingredients” required to determine *S*_[*a,b,j*+1]_, and the arrow from *S*_[*a,b,j*]_ to *S*_[*a,b,j*+1]_ is causal, not in isolation, but through its role in selecting the appropriate row of **Γ**_[*a,b*]_.

Second, following a similar logic, the true holding time 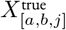 is drawn from an exponential distribution with mean *θ*_[*a,b,k*]_, where *k* is given by the current state *S*_[*a,b,j*]_. That is, *S*_[*a,b,j*]_ filters what element of *θ*_[*a,b,k*]_ is relevant. Then, the sampling procedure takes 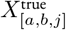 as input, and produces *X*_[*a,b,j*]_ and *C*_[*a,b,j*]_ as output—which, together, specify whether and how 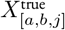 was censored.

Third, our causal graph makes explicit an aspect of the generative model that we have not previously addressed: the causes of *θ* and Γ. In our simulations, we specified that unobserved group-level (*e.g*., habitat quality), individual-level (*e.g*., personality), and dyad-level (*e.g*., kinship) variables influence the values of *θ* and *γ*. We call these variables *U* ^group^, *U* ^ind^, and *U* ^dyad^, respectively. These effects generate substantial heterogeneity between dyads, along with strong patterns of covariation within and between the two sets of parameters, with effect sizes chosen to produce behaviours on a biologically plausible scale (see Figure S2). For example, we generated a negative correlation between *θ*_[*a,b*,1]_ and *θ*_[*a,b*,3]_, indicating that dyads with shorter sojourns in *k* = 1 (*i.e*., those that interact more frequently) also tend to stay for longer in *k* = 3, (*i.e*., to engage in longer grooming bouts). The specific direction or strength of these associations is not important here. What matters is that unobserved variables operating at multiple levels induce a strong dependency structure across dyad-level parameters—the kind of structure we would expect to encounter, in some form, in empirical systems. Lastly, notice that *θ* and Γ are not affected by observation-level causes (*e.g*., time, observer). If they were, they would need to appear within the plate *j*, together with *X, S*, and *C*.

Beyond providing a summary of our generative assumptions, DAGs further support a critical task in causal inference: *causal identification*. The identification of a causal effect consists in determining the conditions under which it can be estimated in principle if researchers had access to an infinitely large data set—and if so, which variables should be included and excluded from a statistical model. For instance, if one is interested in studying the effect of genetic relatedness on the duration spent in one or several behavioural states, and that this effect is *confounded* by dominance rank, graphical criteria (*e.g*., the backdoor criterion; Pearl et al., 2016) can be used to know which variables to include, so that the effect of interest can be estimated without bias. Alternatively, the effect of genetic relatedness on social behaviour might be confounded by unobserved group-level variables (*e.g*., ecological resources), preventing the causal effect to be estimated from the raw association—yet, the system’s hierarchical structure might nonetheless provide further opportunity for causal identification (Byrnes & Dee, 2024; Weinstein & Blei, 2024). Generally, an explicit identification strategy is needed for robust causal inference, especially—though not exclusively—in observational settings. We recommend McElreath (2020) and Pearl et al. (2016) for general introductions to the topic, and Kawam et al. (2025) for animal social networks specifically.

### 2.2. Statistical model

In the previous section, we laid out simple assumptions about the data-generating process leading to focal-follow data—”simple”, if not in a technical sense, at least relative to real-world empirical processes. This task was a sort of engineering task: we assumed generative processes as input, and obtained synthetic data as output. These assumptions will now serve as a basis to build, then validate our statistical model. That is, we turn to a task of reverse-engineering: one where we go from known data as input (the variables *X, C* and *S*) to inferred processes as output—specifically, by estimating the parameters *θ* and Γ.

Our inferential task implies, first, a challenge related to measurement. How can we design a statistical model that connects observed holding times *X* to underlying averages *θ* given that many of the *X*s are censored? For instance, how can we infer that the average holding time of a dyad—say, the dyad (5, 12)—in state *k* = 1 is of two days if we observe it for 40 minutes at a time only? As detailed in section 2.2.1, we will do so by separately modelling censored and uncensored holding times using the Exponential distribution’s PDF and CCDF.

A second challenge relates to a well known issue in animal social network analysis—limited and uneven sampling effort across dyads. Here, we ask: how can we recover *θ* and Γ for each pair of individuals if many dyads were rarely or never observed in certain states? The task of capturing regular, unobserved features like *θ* and *γ* while relying on such irregular, sample-specific features constitutes a common challenge in statistics—one of *regularisation*. As a solution, we design parallel linear models across behavioural dimensions whose parameters are part of a crossed varying-effects structure sharing information across units and behaviours. We define these parameters in sections 2.2.1 to 2.2.3.

#### 2.2.1. Holding times

We start by describing how observed holding times *X* relate to their underlying means *θ*:

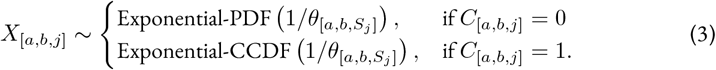

In plain terms: the observed holding times *X* are described as random draws from an Exponential Probability Distribution Function (PDF) if the observation is non-censored; and by an Exponential Complementary Cumulative Density Function (CCDF) if the observation is censored. Both of these density functions are characterised by the same, unknown mean *θ*_[*a,b,k*]_, with *k* being given by *S*_[*a,b,j*]_ (abbreviated *S*_*j*_).

The Exponential PDF assesses how likely it is that, given certain values of *θ*, the non-censored holding time takes on value *X*. This follows straightforwardly from our generative assumptions, and directly mirrors the procedure described in section 2.1.1 (recall that the holding times were literally drawn from Exponential PDFs).

The second part of the likelihood has, however, no direct equivalent in the generative model. Instead, it provides information about *θ* from censored observations: *i.e*. those observations with unknown end times. The Exponential CCDF assesses how plausible it is that, given certain values of *θ*, the end of the holding time *has not been observed yet* after duration *X*. Like in the PDF case, this function quantifies how plausible observed holding times are with different parameter values of *θ*, but it does so by focussing on a different function—and thus, a different aspect—of the Exponential distribution. For readers interested in building a stronger intuition about this modelling technique, we offer more details in section S1.2.2.

Next, the parameters *θ* are described by linear (or additive) models that are part of an overarching multilevel structure, with cross-level varying-effects (Redhead et al., 2024; Snijders & Kenny, 1999). Let us start with *k* = 1, capturing the average holding times without body contact:

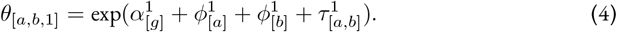

Here, the superscripts denote state *k* = 1, not mathematical powers. Observe, too, that the linear predictors are exponentiated. We do this to ensure that *θ*_[*a,b*,1]_ remains positive even if the predictors are unbounded.

The parameters of equation 4 capture group-, individual-, and dyad-level variation in average holding times—respectively caused by *U* ^group^, *U* ^ind^, and *U* ^dyad^ in our generative model. 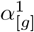 is a group-level intercept. It quantifies the log holding time in state 1 for an average dyad in a social group *g*. The parameters 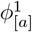 and 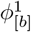 (phi), are individual-level varying (or random) effects for *a* and *b*. These parameters capture individual tendencies to spend more or less time in state *k* = 1 across partners. A low value for 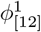 would for instance indicate that an individual, 12, tends to interact more frequently with its conspecifics. Lastly, 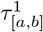 (tau) estimates residual variation in *θ*_[*a,b*,1]_ that is not explained by 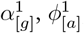, and 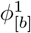 . This parameter is symmetric 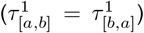, because the state *k* = 1 is non-directed (as opposed, for instance, to states 3 and 4).

We follow the same logic for the time spent in body contact, another undirected state:

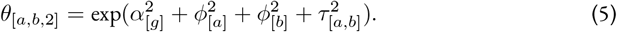

The linear structure is identical to equation 4, but the predictors—distinguished by their superscripts— now capture variation in body contact duration (state *k* = 2).

Things are slightly different for states 3 and 4, which are distinct states in the Markov model’s statespace, but capture the same behaviour—grooming—only in different directions. Thus,

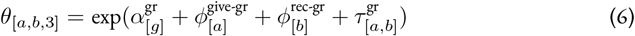

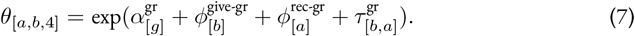

Both linear models share the same group-level intercept 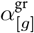, reflecting a common baseline for grooming bout duration, whether exchanged from *a* to *b*, or from *b* to *a* (recall that *a* and *b* are arbitrary labels). The individual-level effects further reflect the directionality of grooming: *ϕ*^give-gr^ captures a tendency to give longer grooming bouts across partners, while *ϕ*^rec-gr^ captures a tendency to receive longer bouts. These roles are reversed across the states 3 and 4. This is the case because *a* is defined as the giver in state 3 and as the received in state 4, and on the flip side, *b* as the giver in *k* = 4 and as the receiver in *k* = 3. Finally, in contrast to *τ*_1_ and *τ*_2_, the dyadic effect *τ*_gr_ is directed 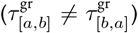, allowing the model to account for asymmetries in dyad-specific grooming tendencies.

#### 2.2.2. Transition probabilities

To describe the transitions dynamics between behavioural states, we build a likelihood that perfectly mirrors the generative model—a simpler case than that of the holding times. Yet, subsequently describing the inferred probabilities *γ* with linear models will require additional modelling tools.

The transition from any current state *S*_[*a,b,j*]_ to the next state *S*_[*a,b,j*+1]_ is given by:

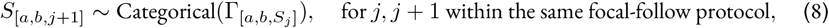

Where Γ_[*a,b,k*]_ is a row in the **Γ**_[*a,b*]_ matrix, as defined in equation 2. The likelihood is only defined for pairs of observations *j* and *j* + 1 that belong to the same focal-follow protocol, because observations that do not satisfy this condition are not directly following one another in the true sequence of behavioural states (Figure 3).

For each non-zero transition probability *γ*—where *γ* are entries of **Γ**—, we define a corresponding latent score *ψ* (“psi”). We organise these scores into a dyad-specific matrix, **Ψ**_[*a,b*]_:

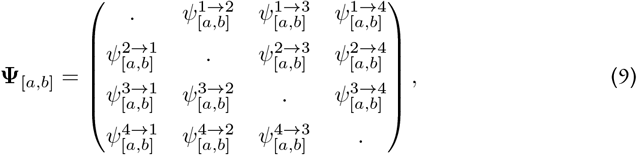

Where the dots on the diagonal represent missing entries. For each state, we define a vector of latent scores Ψ_[*a,b,k*]_, reflecting the probability of a dyad (*a, b*) to transition from its current state *k* to the three possible next states *l* ≠ *k*. These scores are not on the probability scale, and thus, not directly interpretable. But they allow us to map dependent probabilities to linear models using a function called the *softmax*, or *multinomial logit* (McElreath, 2020). This function is analogous to the *logit* function of Binomial GLMs, but works for for more than two dependent probabilities. The probability to transition from a state *k* to another state *l* is given by:

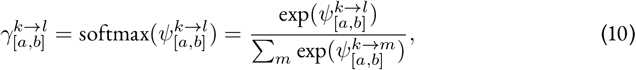

where the sum in the denominator is taken over all possible states *m*≠*k* in {1, …, *K*}. For instance,

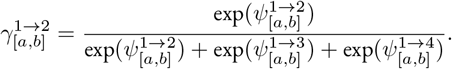

To summarise: the transition from a current state *k* to a future state *l* is described by a vector of probabilities, Γ_[*a,b,k*]_, containing three non-zero probabilities that are deterministically mapped onto a vector of latent scores Ψ_[*a,b,k*]_ using a softmax function.

What shall we do with the scores *ψ*? Like the parameters *θ* above, they will be described by linear models. Recall that from any current state *k*, we have three possible categorical outcomes *l* ≠ *k*. That means that we only need to model two of the outcomes *l* directly, and can use the third one as a pivot. The logic is again similar to the Binomial case. In Binomial GLMs, there are two possible outcomes, 0 and 1, and we only model the probability of “successes”. Once we know the probability of “success”, we automatically know the probability of “failure” by subtracting this probability to one. Similarly, suppose that from the current state (say, *k* = 1), a dyad has: (i) a probability of 0 to transition to state 1; (ii) a probability of 0.3 to transition to state 2; and (iii) a probability of 0.3 to transition to state 3. Then, we automatically know that (iv) it has a probability of 1 − 0 − 0.3 − 0.3 = 0.4 to transition to state 4. The idea is thus to model all the moving parts except one, because the latter can be automatically computed from the former.

In the case of the transitions from *k* = 1, a dyad (*a, b*) can transition to *l* ∈ {2, 3, 4 }. We define the transition to *l* = 2 as pivot, and assign it an arbitrary fixed value of zero.

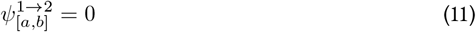

The other two are described by models that are structurally very similar to equations 6-7:

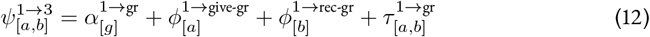

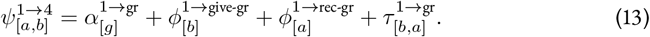

Here, 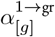 captures the baseline probability of transitioning from *k* = 1 to *l* ∈ {3, 4} for group 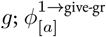 captures the tendency of *a* to transition from *k* = 1 to 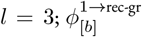 captures the tendency of *b* to transition from *k* = 1 to *l* = 3; and 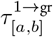 captures the directed dyad-level tendency to transition from *k* = 1 to *l* = 3; all, on the latent linear scale.

We model the transition probabilities from *k* ∈ {2, 3, 4} by applying the same logic, and enforce similar symmetry constraints. However, for brevity, we do not present them in the main text. They are shown in section S1.2.1.

#### 2.2.3. Multilevel structure

Average holding times *θ* and transition probabilities *γ*, for all *k*s and *l*s have now been described— albeit indirectly for *γ*—by linear predictors capturing group-, individual-, and dyad-level variation. Crucially, within each of these levels, different behaviours are likely associated with one another. In fact, we know that such correlations exist in our generative model because unobserved variables *U* ^ind^ and *U* ^dyad^ influence all dimensions of *θ* and *γ* (Figure 4). This means that individuals who give long grooming bouts across partners might for instance receive longer grooming bouts in return. Such gregarious individuals might also be more likely to transition from *k* = 2 to *l* ∈ {3, 4} than to *l* = 1 (*i.e*., to experience longer sequences of affiliative states). Similarly, unobserved dyad-level variables cause associations between dyad-specific tendencies to, for instance, spend less time in *k* = 1 (*i.e*., to engage more frequently in affiliative interactions) and to transition more often from *k* = 3 to *l* = 4 (*i.e*., to perform immediate grooming reciprocation).

We can leverage these correlations to improve estimation of *θ* and *γ*, by treating individual- and dyad-level parameters as draws from adaptive multivariate prior distributions. Specifically, we describe the vector of individual-level effects for individuals *a* as draws from a Multivariate Normal distribution (MVN):

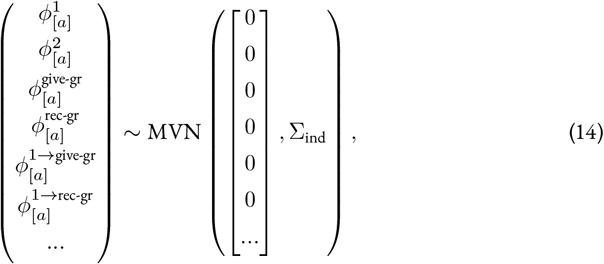

Where each parameter *ϕ* represents the deviation from the intercept of a specific individual—indices *a* ∈ {1, 2, …, *N*} —, and for a given behavioural dimension—*i.e*., *k* = 1, *k* = 2, tendency to give longer grooming bouts (“give-gr”), and so on. We only show the individual parameters introduced in the main text. Yet, the full array includes additional parameters (denoted by the ellipsis) that correspond to transitions from states *k* ∈ {2, 3, 4}, for a total of 12 dimensions.

The Multivariate Normal distribution is centred on zero, and governed by a variance-covariance matrix Σ_ind_:

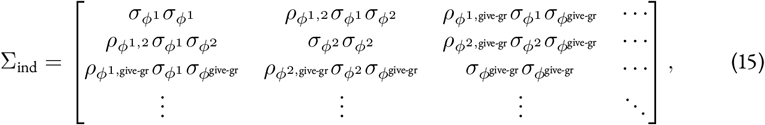

With *σ* (sigma) and *ρ* (rho) describing standard deviations and correlation coefficients, respectively. This matrix may appear intimidating at first glance, but its structure is relatively straightforward. Its diagonal entries capture inter-individual variation for each behavioural dimension: 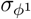 is the standard variation of individual effects for state 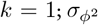, for state *k* = 2, and so on. This is the *variance* part of the “variance-covariance” matrix. On the matrix’s off-diagonal, we find all possible associations between behavioural dimensions—measured as *covariances*. For example, 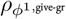 is the correlation coefficient between the individual tendency to spend longer in *k* = 1, and the tendency to groom others for longer.

The dyad-level parameters are similarly defined as crossed-varying effects:

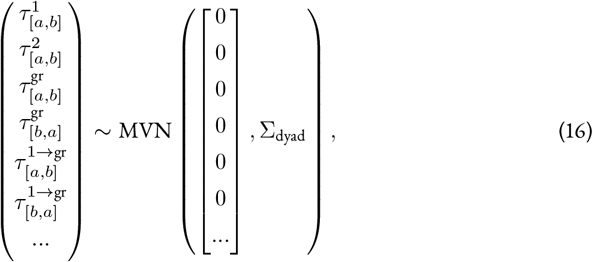

Where Σ_dyad_ is a 12-dimensional variance-covariance matrix. It is structured like Σ_ind_, but is subject to additional symmetry constraints similar to those of Social Relations Models (Redhead et al., 2024; Ross et al., 2023, 2025)—*i.e*., the standard deviation of *τ* within each directed behavioural dimension is shared in both directions (*a* to *b*, and *b* to *a*; for a full model description, see section S1.2.1).

In sum, although observational data will usually be sparse for many dyads and behavioural states, we designed a strategy to estimate *θ* and *γ*, by leveraging the rich patterns of covariation between individual- and dyad-level effects, within as well as between behavioural dimensions.

Before we move on to validating this strategy, there is one more step left to complete our statistical model: to assign a prior distribution to its parameters. Although often overlooked, defining the prior model is not just a technical formality. It is yet another example of scientific intuition guiding formal modelling. Here, behavioural ecologists know that the range of plausible values for *α*^1^, reflecting the time between episodes of body contact or grooming, is on the order of days for macaques—say, from 0 to hundreds, being conservative. By tuning parameter values while plotting the implied prior distribution for *θ*_[*a,b,k*]_ and associated predictions for *X* (Figure S6), we deduce that a biologically reasonable range is covered by:

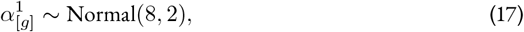

Where the same prior is given to each group *g*. In contrast, the plausible range for body contact or grooming duration is on the order of minutes. Let us say, loosely: between 0 and a few dozen minutes, for an average dyad in uninterrupted body contact (excluding night time), or an uninterrupted grooming bout. Again, by plotting the implied *θ* and *X* (Figure S7), we end up on:

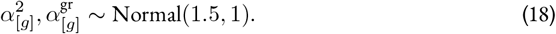

We chose similarly conservative prior distributions for the remaining parameters (see section S1.2.3 for a visualisation):

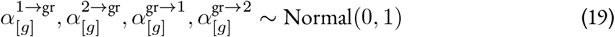

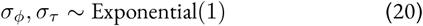

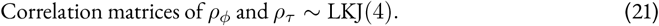

### 2.3. Posterior model

We now have an estimation strategy to recover the parameters of our generative model. Does it actually work? To evaluate its performance, we implemented the statistical model in the probabilistic programming language *Stan* (Carpenter et al., 2017) and ran it via R using the *CmdStanR* interface (Češnovar et al., 2021). Doing so, we drew Markov Chain Monte Carlo (MCMC) samples to approximate the model’s *posterior distribution*. Below, we compare these posterior distributions to the true parameter values of the generative model. We also use the posterior samples to generate *predictions* in R, which we compare to the observed data.

#### 2.3.1. Posterior estimates

We start by plotting the marginal posterior probability distributions for the parameters *θ* and *γ* of each dyad, which we call 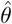 and 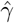 (Figure 5). Posterior probability distributions simply describe the plausibility of parameter values, given the observed data and chosen prior distribution. On the figure, a darker colour indicates a higher plausibility. What we want to see is a good overlap between estimates and red points across parameters—which is precisely what we observe.^2^

**Figure 5.**
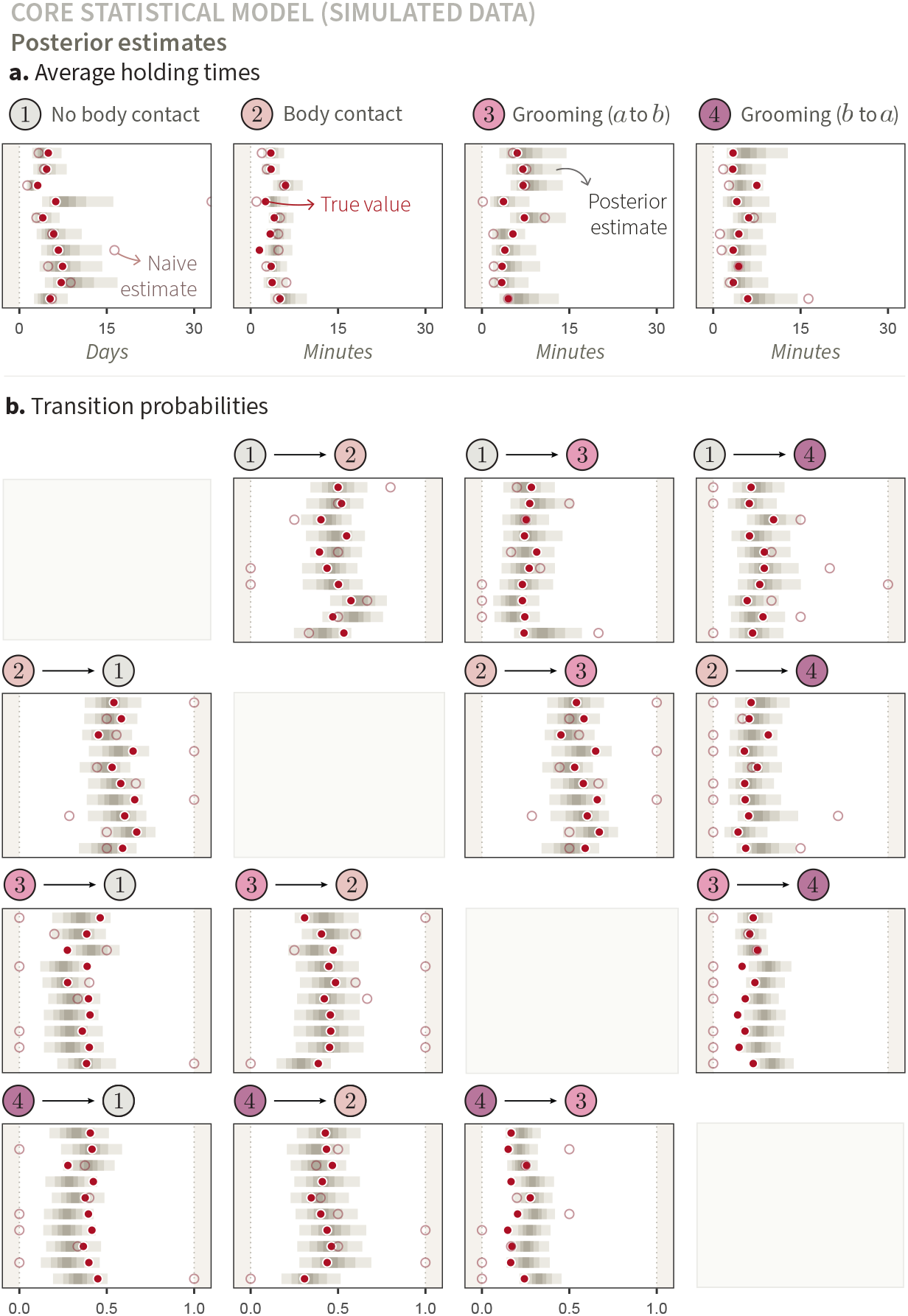
Posterior estimates of average holding times and transition probabilities for ten dyads (synthetic data). We show possible parameter values on the x-axis, and the same ten dyads for all panels on the y-axis. Across panels, we also represent the marginal posterior distributions using percentile intervals of 25%, 50%, 75%, and 95% of posterior samples, from lightest to darkest. These are compared to the true generative parameter values (solid red points) and to naive estimates (empty pink points). **(a.)** We obtained the naive estimates by dividing the total amount of time spent in a given state *k* by the number of times that the dyad was observed leaving the state. Naive estimates that stand on the edge of the plots’ frame indicate an estimate of +∞ (*i.e*., if a dyad has never been observed ending a behavioural state *k*, in which case the observed duration is divided by 0). Missing naive estimates indicate that the dyad was never observed in the state *k* in question. **(b.)** We computed naive estimates by dividing the number of observed transitions from a state *k* to another state *l*, divided by the total number of transitions from *k*. Missing naive estimates indicate that the dyad was never observed transitioning from the state *k* in question, whereas naive estimates of zero indicate that the dyad was observed transitioning from *k* but never to the specific state *l*.

We further compare the posterior estimates to naive estimates: that is, to sample averages analogous to common indices in the field (*e.g*., SRI, reciprocity index, Hinde index; Farine & Whitehead, 2015; Hinde & Atkinson, 1970; Kaburu & Newton-Fisher, 2015; Silk et al., 2013). In Figure 5**a**, we obtained the naive estimates by dividing the total amount of time spent in a given state *k* by the number of times that the dyad was observed leaving the state. For example, if a dyad (4, 8) was observed initiating contact—such as starting to groom or entering in body contact—ten times over the course of 1, 000 minutes of observation, the naive estimate for *θ*_[4,8,1]_ would be 1, 000*/*10 = 100 minutes. (Equivalently, we could think of the *rate* at which a dyad initiates body contact, by dividing the number of events by observation time—the idea underlying the SRI). In Figure 5**b**, we computed naive estimates by dividing the number of observed transitions from a state *k* to another state *l*, divided by the total number of transitions from *k*. For instance, consider the following transitions of a dyad from state 1: once to state 2, twice to state 3, and never to state 4, then the naive estimate for the transition from state 1 to state 2 is equal to 1*/*(1 + 2 + 0) = 1*/*3.

Because data is relatively sparse for most combinations of dyads and states, the naive estimates are bad—they have, by definition, no measure of uncertainty, and spectacularly fail to approach the true values in several cases. On the other hand, the multilevel structure produces regularised estimates, which are pulled away from the population mean only when sufficient data are available to support that deviation. Additionally, we show estimates of population-level parameters for all states *k* and *l*, in Figure S9.

#### 2.3.2. Posterior predictions

Comparing posterior estimates to true parameter value is a powerful way to validate that a statistical model does what it should, but it can only be done using synthetic data, where the true values are known. Another, complementary, approach to assess a Bayesian model’s performance is to compare the predictions implied by its posterior distribution, to its input data—whether synthetic or empirical. That is, instead of comparing true parameter values to inferred parameter values as we did earlier, we here compare (i) the distribution of outcome variables that was used to update the statistical model and (ii) the distribution of outcome variables that is implied by the inferred parameter values, as a result of fitting (i) to the statistical model. This procedure is called *posterior predictive checks* (Figure 6). Specifically, we take (i) the observed holding times and state transitions generated by the generative model in section 2.1, and compare them to (ii) random draws from Exponential and Categorical distributions with, as parameters, posterior draws of the estimates 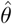and 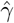, respectively.

**Figure 6.**
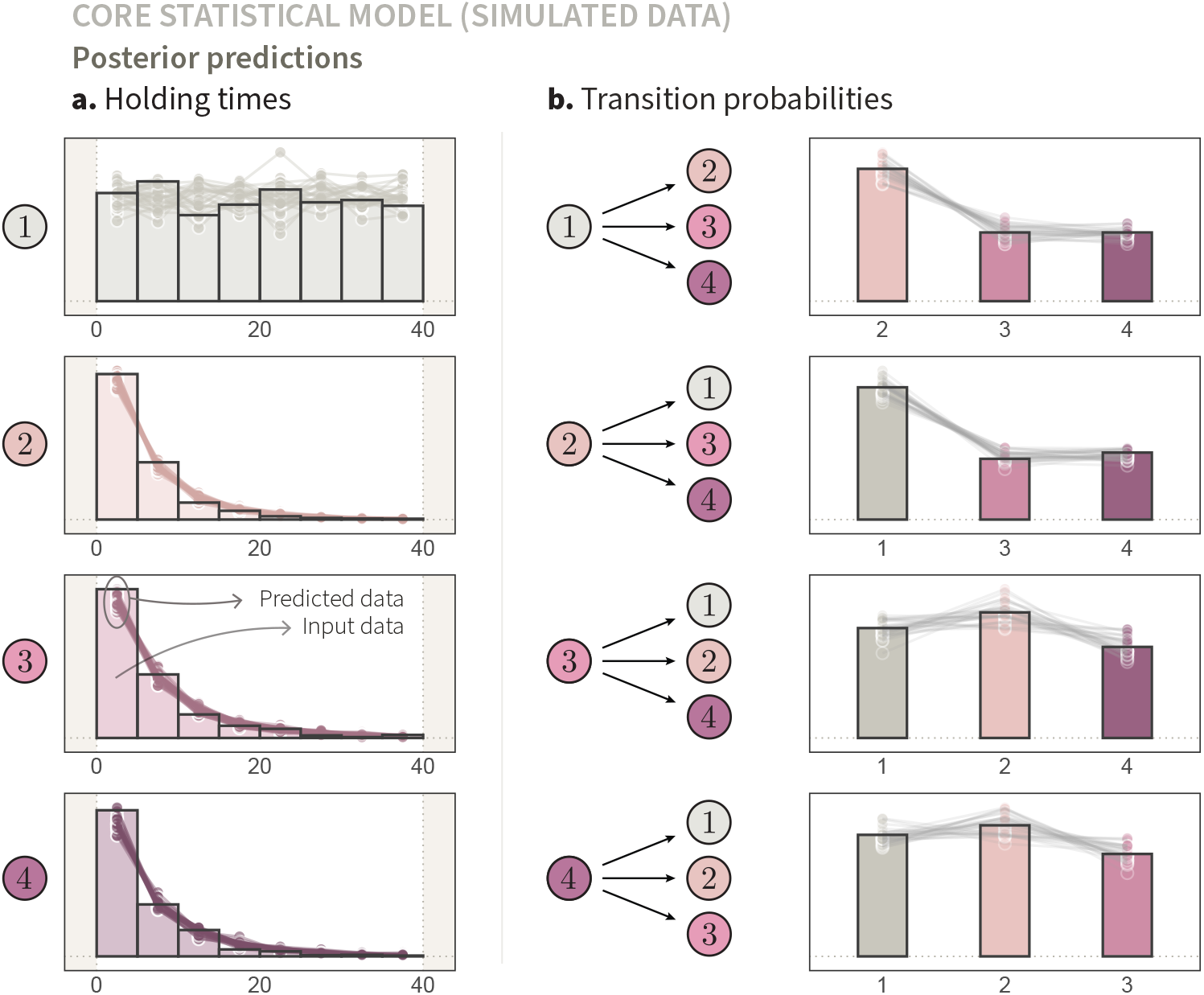
Posterior predictive distribution of the core statistical model compared to synthetic input data. This figure compares two distributions: (i) the input data generated by the generative model described in section 2.1, shown as solid histograms in panel **a** and bar plots in panel **b**; and (ii) the posterior predictive distribution implied by fitting the input data to the statistical model (see sections 2.2 and 2.3.2), shown as sets of points connected by lines, with each set corresponding to one posterior draw *n*. (**a**.) The four sub-panels show the distribution of observed durations (in minutes) spent in behavioural states *k* ∈ {1, 2, 3, 4 }, from top to bottom. We display only non-censored holding times from the input data (*C*_[*a,b,j*]_ = 0) and posterior predictive holding times shorter than 40 minutes. Note that the distribution of holding times for *k* = 1 appears flat because its average is orders of magnitude greater than the values on the x-axis (*i.e*., situated far to the right). (**b**.) The four sub-panels show the distribution of observed transitions from a current state *k* ∈ {1, 2, 3, 4} to future states, from top to bottom. The x-axis corresponds to possible next states, and the y-axis corresponds to observed counts—as opposed to Figure 3, where the y-axis represented probability density and mass.

Let us start with the holding times:

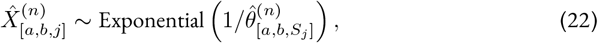

with *n* ∈ {1, …, *N*_draws_}. In plain terms: for each posterior draw *n*, we sample one holding time 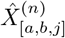 per input observation *j*, from an exponential distribution characterised by a mean 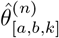, with *k* being given by the state of the input observation *j, S*_[*a,b,j*]_. This procedure produces a “twin” synthetic observation per empirical observation *j*, and therefore, a full distribution 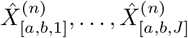 for each draw *n*.

Our generative model assumes that all focal-animal protocols are complete (*i.e*., that they last 40 minutes). In many empirical systems, however—including the one we will focus on in the next section— censoring might also occur when focal animals are lost during observation, interrupting the protocol and producing an excess of shorter holding times. Because this censoring process is not included in the generative model, we focus our assessment on how well the model predicts uncensored data. Accordingly, we subset the predictions 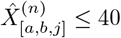 from the distribution 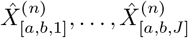, and compare the resulting set to empirical uncensored holding times (*X*_[*a,b,j*]_ | *C*_[*a,b,j*]_ = 0). Each set of connected points and lines in Figure 6**a** corresponds to one posterior draw *n*, describing the distribution of the synthetic holding times 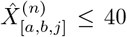. Taken together, these posterior predictive distributions capture variation in holding times that emerges from three key sources: the randomness of the exponential process, variability between dyads, and inferential uncertainty of 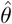 and 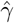.

Similarly, we obtain a posterior predictive distribution of state transitions by:

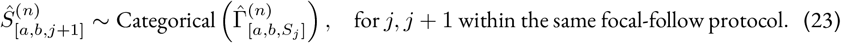

That is, for each posterior sample *n*, and for each pair of observations *j* and *j* + 1 belonging to the same focal-follow protocol, we draw an alternative next state 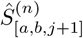, based on the estimated probability distribution 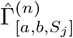 . As in equation 22, we obtain a full distribution of predicted transitions per posterior draw *n*, which we represent by a set of connected points and lines (Figure 6**b**).

Overall, the overlap between the input data and posterior predictions is satisfactory (Figure 6). Together with the previous section, this confirms that our statistical model can theoretically recover the processes underlying the observed holding times and state transitions—at least, given that the assumptions encoded in the generative model are satisfied. This is good news, but not surprising: the statistical model largely mirrors the generative model. As we shall see below, the alignment between posterior predictions and input data will not always be so close when dealing with empirical data.

### 2.4. Empirical system

We finally turn to applying our statistical model to an empirical data set. In doing so, we assume that the data set in question arose from the processes described by our generative model (section 2.1). We provide a brief overview of the data below.

Our data set was collected between January 2015 and June 2017, as part of a long-term research project on wild Assamese macaques at Phu Khieo Wildlife Sanctuary in Northeastern Thailand. We focused on fully habituated adult macaques belonging to four stable social groups (*N* = 82 individuals and 849 dyads). Observers conducted continuous 40-minute focal-follow protocols, during which they recorded all instances of dyadic interactions, including the behavioural actions that marked the beginning and end of the states described in Figure 2.

The processing of the raw behavioural data was minimal. It primarily involved deducing dyadic behavioural states from focal-follow data. For each observed sequence of actions recorded during a focal-follow protocol, we deduced the corresponding dyadic states for all dyads that included the focal individual. For example, if individual 12 was observed engaging in no social interactions for the first 30 minutes, briefly grooming individual 19, and then remaining inactive for the rest of the 40-minute protocol, we inferred that all dyads involving individual 12 were in state *k* = 1 (no contact) throughout, with the exception of (12, 19), which transitioned from state *k* = 1, to state *k* = 3 (grooming), and back to state *k* = 1. Dyads were defined based on daily presence-absence data. For simplicity, if males immigrated during the study period and thus belonged to multiple groups, we retained only their observations from a single group, thereby maintaining strict nesting of individuals within groups; see Figure 4. (While this assumption can be easily relaxed, we judged that introducing additional complexity in the multilevel structure of the causal system would not benefit our primarily conceptual and technical goals.)

Following the procedure above, we arrived at a set of *J* = 318, 952 observations. Notice, once again, that the focal data were never aggregated. In fact, not only did we keep every single relevant interaction that was observed over more than two years, we also counted the “non-interactions” with potential interaction partners as dyadic sojourns in state *k* = 1. The resulting observations *j* are all characterised by *X, C*, and *S*; *i.e*., they have the same structure as the synthetic data generated by our generative model (Figure 3)—except, of course, that the true underlying durations *X*^true^ are unknown.

Unsurprisingly for an empirical data set, sampling effort strongly varied across individuals, dyads, and states. Individual animals were followed for anywhere between 0.05 and 192 hours of cumulative observation time (Figure S10). Because of this variation, combined with biological differences between dyads, many dyads were never observed in certain states *k* (Figure S12) or transitioning from one state *k* to another state *l* (Figure S13). Such sparsity is common in social network data, making the use of aggregated indices (*e.g*., SRI) inappropriate (Redhead et al., 2025). Fortunately, it is precisely the kind of setting that our regularisation strategy (see section 2.2) is designed for.

### 2.5. Empirical results

#### 2.5.1. Posterior estimates

By confronting our statistical model with empirical data, we are able to draw descriptive inferences about how Assamese macaques interact socially in the population under study.

After verifying that the MCMC diagnostics indicate good convergence and sampling efficiency (see Sections S1.4.1), we inspect the dyadic estimates, 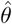and 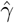. These parameters represent our state of knowledge about how long each pair of macaques sojourns in each behavioural state (*i.e*., no body contact, body contact, grooming) and about the transitions dynamics between them—thereby capturing how interactions unfold over time between the two individuals. Ten randomly chosen dyads are displayed in Figure 7. This figure is almost identical to Figure 5, except that, because we are here dealing with real-world monkeys, as opposed to synthetic ones, the true values are, naturally, unknown. We notice that the dyadic estimates exhibit varying levels of uncertainty (posterior distributions of different widths), partly reflecting differences in sampling effort. We also observe that the ratio indices—the naive estimates—often deviate from the posterior distributions, and, in many cases, are missing entirely because the dyads were never observed in the relevant states *k*.

**Figure 7.**
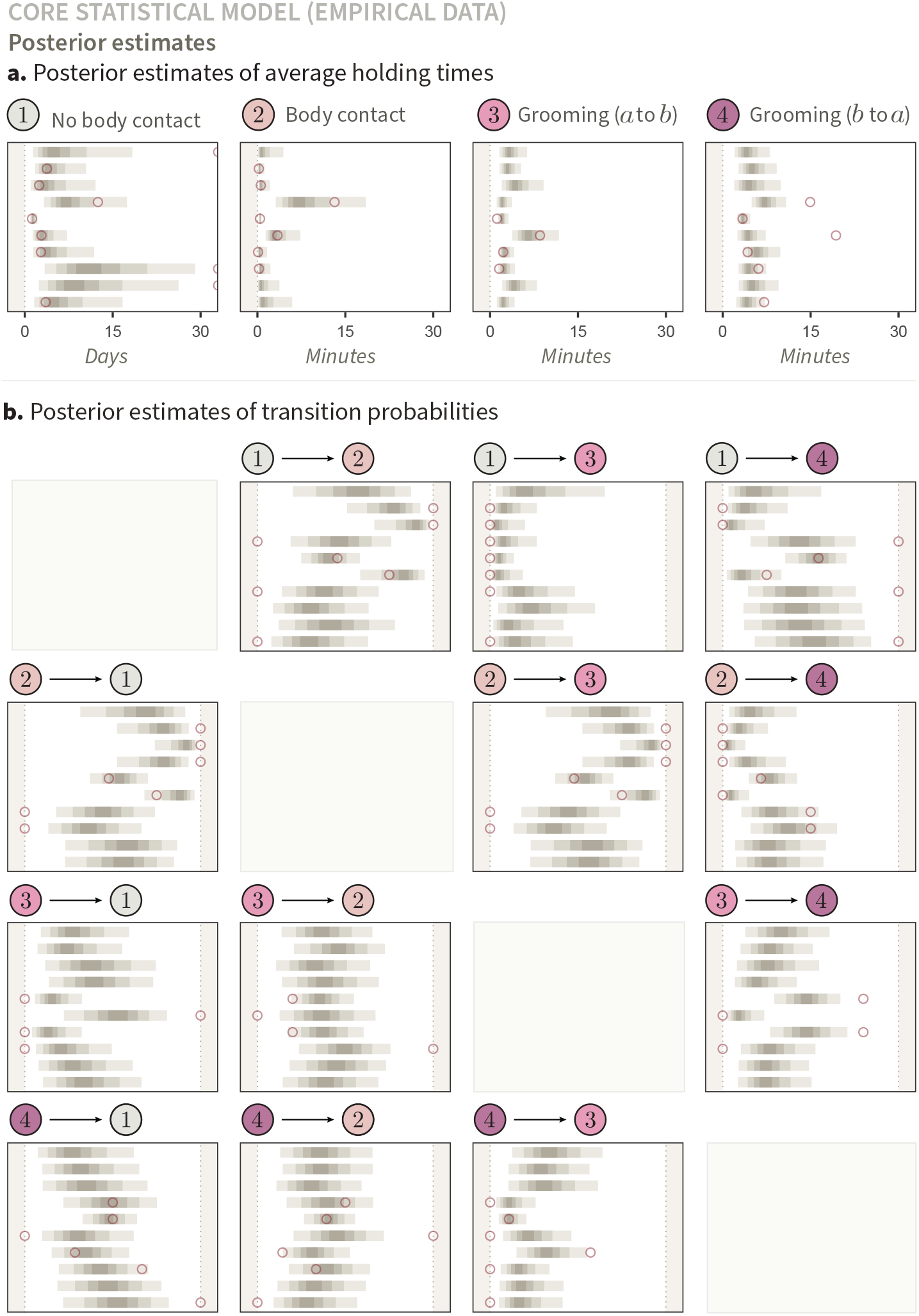
Posterior estimates of average holding times and transition probabilities for ten dyads (empirical data). This figure is similar to Figure 5, except that the input data were here collected in a population of wild Assamese macaques. Consequently, the true values—unknown for empirical monkeys—are absent.

To draw population-level inferences and characterise average behavioural patterns—about both the durations spent in each behavioural state and the expected transitions between states—, we can average the parameters 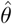and 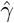 across all dyads and for each behavioural dimension (Figure 8). For instance, the average duration in state 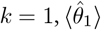, is given by:

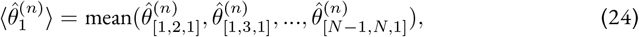

**Figure 8.**
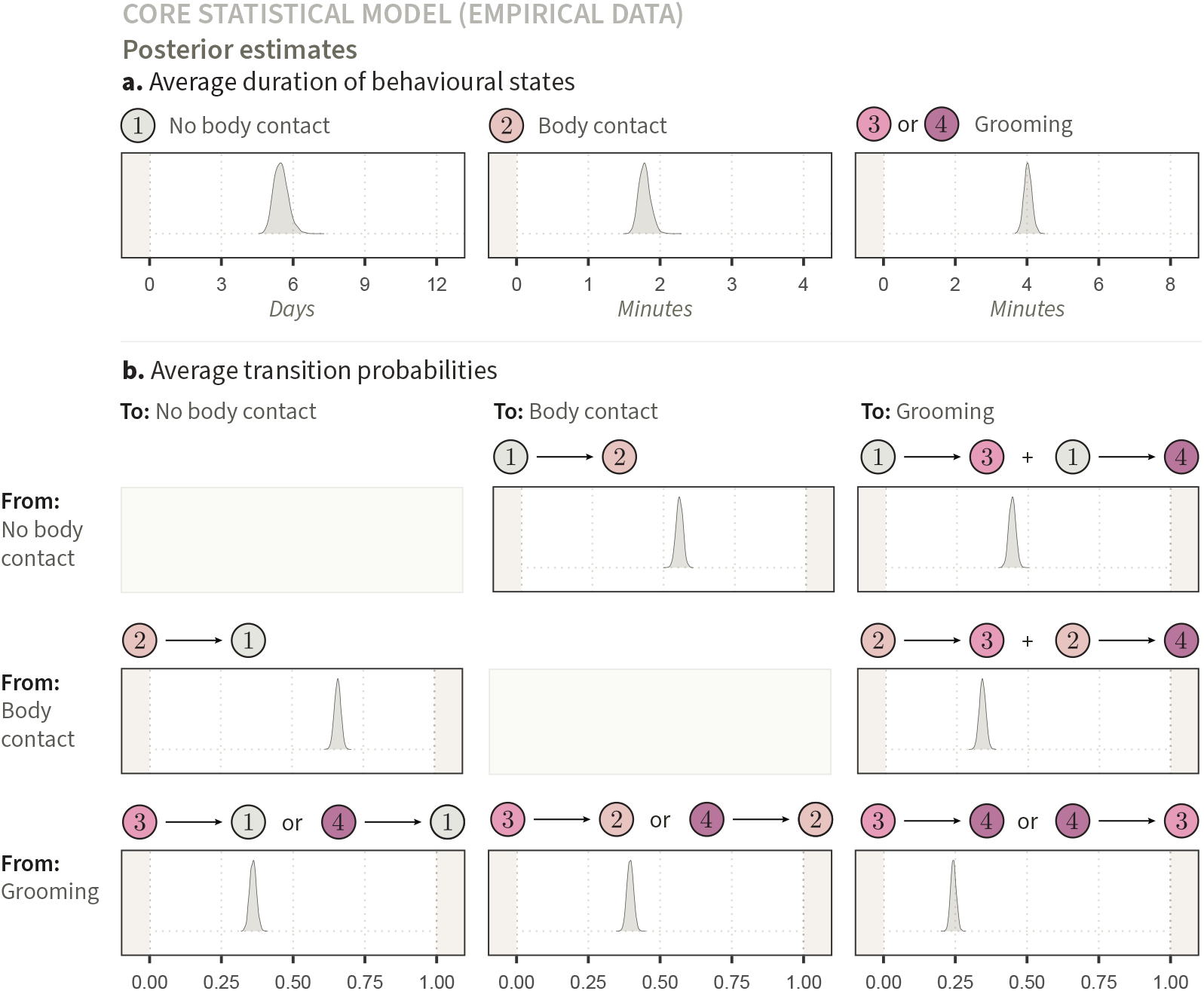
Posterior estimates of expected holding times and transition probabilities, averaged across all dyads (empirical data). (**a**.) Expected holding times in three types of behavioural states, where we combine states *k* = 3 and *k* = 4 (grooming). The posterior distributions correspond to, from left to right: 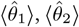, and 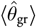 . (**b**.) Estimated transition probabilities between states. The density correspond to, from left to right and top to bottom: 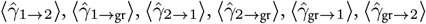, and 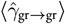. Titles using “or” indicate that the relevant vectors (*e.g*., grooming from *a* to *b* and from *b* to *a*) were appended before computing the average, as illustrated in the main text. In contrast, “+” indicates that vectors were summed. For instance, the top-right panel of **b** represents the average probability that a dyad transitions from state *k* = 1 (no body contact) to grooming, in either direction—which we obtain by summing the expected probability to transition from *k* = 1 to *l* = 3, and from *k* = 1 to *l* = 4.

Where *n* represent posterior draws. We follow the same logic for the other behaviours, combining two behavioural dimensions when necessary. For example, the expected grooming bout duration, 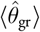, is given by taking an average across dyads of both 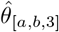and 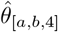:

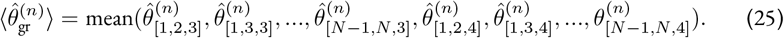

The estimates in Figure 8 can be directly interpreted as probabilistic estimates of population averages. For example, we learn that pairs of individuals interact, on average, every 5.1 to 5.9 days—with this range corresponding to the 80% percentile interval (from the 10th to the 90th percentile) of the posterior distribution for the population mean, and thus, to the most plausible values in light of the available evidence. In this context, and throughout the manuscript, a “day” corresponds to a 12-hour period. We also observe that, when dyads do interact, grooming bouts last on average 3.9-4.2 minutes, and are followed by direct grooming reciprocation in 23-26% of cases (still using an 80% percentile interval; hereafter, our default interval).

#### 2.5.2. Posterior predictions

In addition to providing descriptive inferences, fitting empirical data to the core statistical model can also help us assess its in-sample predictive performance. Following the same procedure described in section 2.3.2, we conducted posterior predictive checks to compare the empirical distribution of holding times and transitions, across dyads, to predicted outcomes drawn from Exponential and Categorical distributions with parameters 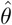and 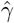 (Figure 9). The correspondence between empirical and posterior predictive distributions appears almost perfect—except in one clear case, *k* = 1, where the predicted holding times diverge from the empirical distribution. Our model underestimates the frequency of short holding times (*i.e*., less than 5 minutes) and overestimate the frequency of longer ones in state *k* = 1. This may be caused by a number of factors. One of them is that our generative assumptions—and associated statistical model—ignore space entirely. In nature, a pair of individuals might briefly interrupt physical contact while remaining in close proximity—for instance, to move to a different branch or catch a passing insect—before resuming their interaction. We return to this issue below.

**Figure 9.**
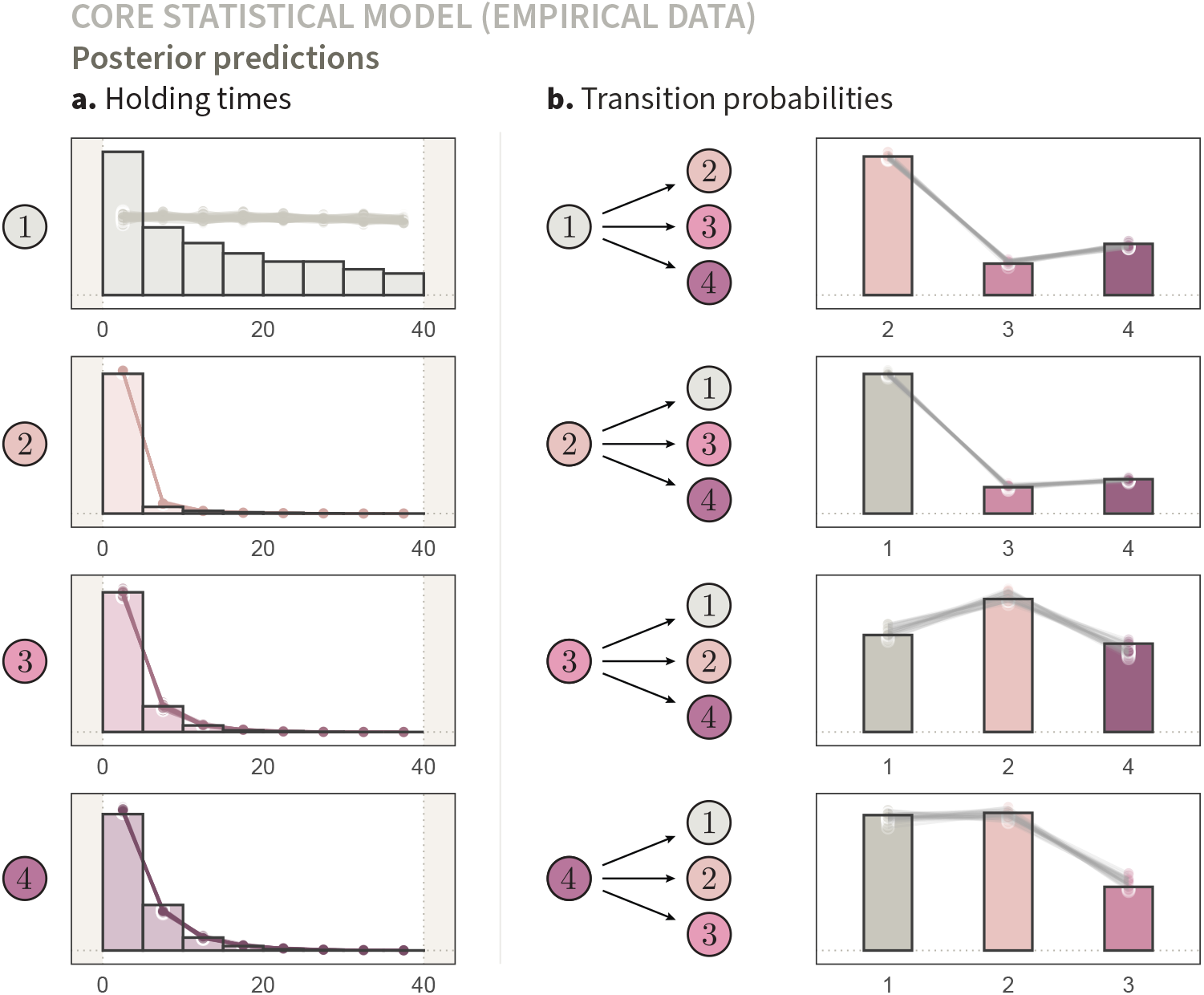
Posterior predictive distribution of the core statistical model compared to empirical input data. This figure is identical to Figure 6—except that here, the input data does not come from our generative model, but was instead collected in a population of wild Assamese macaques. We observe that the posterior predictive distributions align well with the empirical distributions, with the exception of the holding times in *k* = 1, indicating that our generative model omits certain aspects of the empirical data-generating process.

### 2.6. From scaffolds to inferences

In the sections above, we introduced a generative model encoding relatively simple biological and measurement processes leading to behavioural observations structured like focal-follows data. We then designed a statistical model to recover the generative model’s parameters, and validated that it could do so with sparse social network data. We fitted the statistical model with an empirical data set collected in wild Assamese macaques, and saw that it could recover key empirical patterns, with the exception of holding times in state *k* = 1. These models constituted scaffolds, and as such, were not used to address particular theoretical questions.

Building upon this foundation, we turn to causal inference. In the next sections, we showcase how behavioural ecologists can study the causal effect of phenotypic features on dynamic patterns of social interactions. Specifically, we focus on how individual sex affects the time spent in different dyadic behavioural states. We start by adding sex into the generative and corresponding statistical models. Given the observed mismatch between posterior predictions and empirical data, we also introduce two sub-states for *k* = 1, corresponding to short and long sojourns, and design a corresponding likelihood function. With this extended statistical model at hand, we finally compute probabilistic estimates for well-defined causal questions, while correctly accounting for inferential uncertainty and for heterogeneity between dyads.

## 3. Extended models

### 3.1. Generative model

Building upon our core generative model, we introduce two modifications to its causal structure (Figure 10). First, alongside the multiple exogenous variables *U* —which represent unobserved group-, individual-, and dyad-level causes (omitted from the causal graph for visual clarity)—we now include individual sex as a cause of *θ*. Individual sex is denoted by *Z*_[*a*]_ ∈ {1, 2}, where 1 indicates male and 2 female. Both *Z*_[*a*]_ and *Z*_[*b*]_ affect *θ*_[*a,b,k*]_. Thus, for each level *k*, and for each dyad (*a, b*), a state-specific function *f*_*k*_ describes the mechanism assigning the value of *θ*_[*a,b,k*]_:

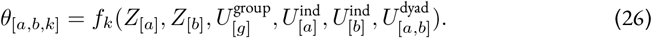

**Figure 10.**
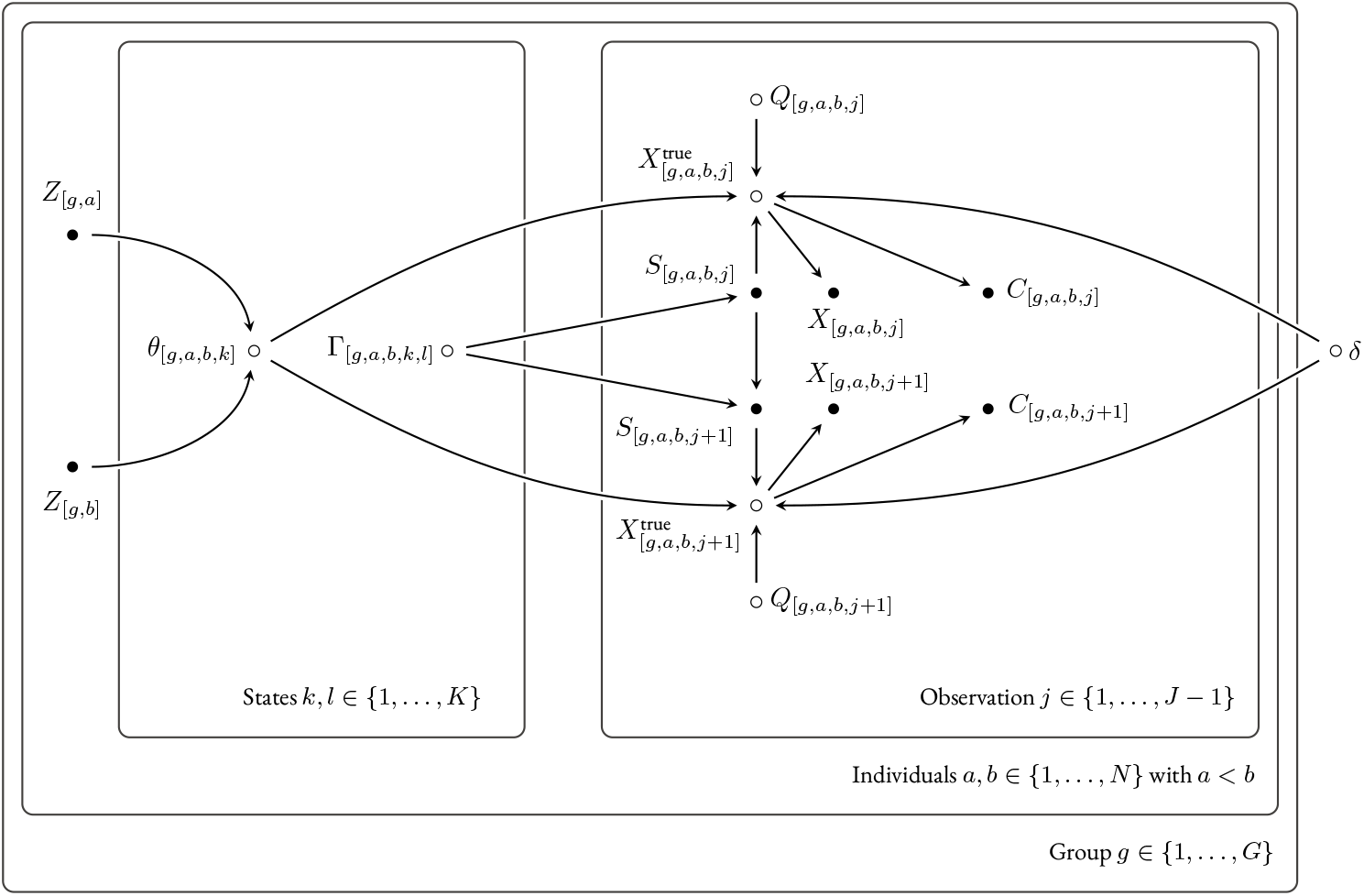
Causal graph for our extended generative model. The graph is similar to Figure 4, with three differences. Fist, we do not explicitly represent *U* ^group^, *U* ^ind^, and *U* ^dyad^ for readability, but they continue to act as unobserved causes of *θ* and *γ*, consistently with Figure 4. Second, individual sexes, *Z*_[*a*]_ and *Z*_[*b*]_, jointly influence the time spent in different behavioural states, as indicated by arrows pointing into *θ*. Third, the true holding times *X*^true^ are now affected not only by *θ*, but also by *δ* and *Q. Q* indicates whether the holding time belongs to the long or short sub-type; in the latter case, the holding time is drawn from a distribution with mean *δ*.

Our inferential task will thus be to design a statistical model that approximates this function—and, in the case of synthetic data, can recover it—at least to the extent necessary to compute the marginal effect of sex. We will return to this idea below.

Second, individuals who transition from *k* ∈ {2, 3, 4} to *l* = 1 can now transition to either a short or a long sojourn in *k* = 1. That is, we assume two *sub-*states for *k* = 1. In short sub-state *k* = 1, two individuals are assumed to briefly interrupt a sequence of direct social interactions (whether grooming or body contact) while remaining in proximity to one another. For example, they might wish to move from a branch to another before resuming affiliative interactions. The longer sub-state is similar to the core model, where the holding time, on average much longer, reflects the interval between two distinct sequences of direct interactions.

Formally, for each observation *j*, a binary variable *Q* that indicates the sojourn type, is given by (Figure 10):

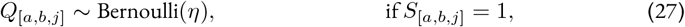

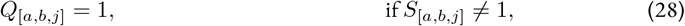

That is, there is no variation in sojourn type *Q* for body contact (*k* = 2) and grooming (*k* ∈ {3, 4}), but each sojourn in state *k* = 1 has a probability *η* ∈ [0, 1] to be of the “long” type (*Q* = 1).

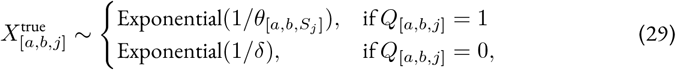

If *Q* = 1, then the true holding time *X*^true^ is drawn from an exponential distribution with mean *θ*_[*a,b,k*]_, with *k* given by *S*_[*a,b,j*]_—so, exactly like in section 2.1. If instead, *Q* = 0 (which only applies to *S*_[*a,b,j*]_ = 1), then *X*^true^ is drawn from a distribution with mean *δ*. For simplicity, we assume that *δ*, the average holding time in the short sub-state *k* = 1, is fixed across dyads.

### 3.2. Statistical model

We then design a statistical model that accounts for the heterogeneity of holding times in states *k* = 1 caused by the two sub-states, and capture the causal effect of sex on *θ*.

The challenge with the two sub-states *k* = 1 is that they are hidden to observers. That is, in an inferential context, the value of *Q* for any observation *j* is unknown. All we can do is assess is how compatible a particular holding time is with either a short or a long sub-state. For instance, if a holding time *X* in state 1 lasts 30 seconds, it is much more plausible that it comes from a short sub-state than a long one. In contrast, a dyad remaining in state 1 during entire focal-follow protocol of 40 minutes is likely sojourning in a long sub-state.

Bayesian modelling offers tools to quantify how plausible these possibilities are, while estimating the averages *θ* and *δ* themselves. To do so, we define a likelihood function for the observed holding times *X*, given *C, S*, and the model parameters: *p*(*X*|*C, S, θ, δ, π*) =

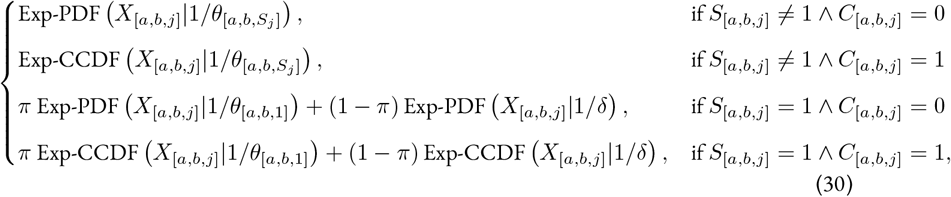

Where “Exp” refers to the exponential distribution; *π* ∈ [0, 1] is a parameter quantifying the probability that any observation *j* in state 1 is a long sub-state (*i.e*., that *Q*_[*a,b,j*]_ = 1); and the ∧ symbol corresponds to the logical conjunction “and”.

The first two lines of equation 30 are the likelihood for states *k* ∈ { 2, 3, 4} . They are identical to equation 3—although, to be consistent with the rest of the likelihood, the notation is slightly different here. The third line gives the likelihood that an *uncensored* holding time *X* is drawn (i) from an exponential PDF with mean *θ*_[*a,b*,1]_, weighted by the probability that it comes from a long sub-state, or (ii) from an exponential PDF with mean *δ*, weighted by the probability that it comes from a short sub-state. In other words, this line assesses how compatible different combinations of values for *θ, δ*, and *π* are with uncensored holding times *X* in state *k* = 1. The fourth line is the likelihood for *censored* holding times *X*, and thus uses an exponential CCDF instead of a PDF (see section S1.2.2). Lastly, note that *π* is distinct from *η. η* is the probability that a sojourn in state *k* = 1 satisfies *Q* = 1, whereas *π* is the probability that a given observation *j*—often, a fragment of the full sojourn—satisfies *Q* = 1. Because the same sojourn might be fragmented into several observations *j*, we expect *π* to be greater than *η*. (Think that even if individuals transitioning to *k* = 1 transition to a short sub-state only 50% of the time, the vast majority of dyadic observation time in *k* = 1 will be fragments of long sojourns).

We then describe variation in the average holding times *θ*, including, the variation caused by individual sex. Beginning with state *k* = 1:

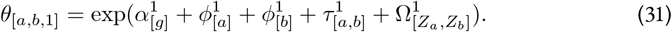

Here, Ω^1^ (uppercase omega) is a 2 × 2 matrix capturing the structuring effects of sex:

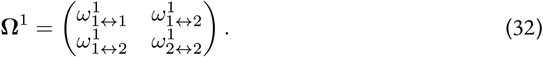

Each entry *ω* (lowercase omega) represents the baseline tendency of a sex combination, relative to its social group *g*. For instance, if individual 5 is male (*Z*_[5]_ = 1) and individual 8 is female (*Z*_[8]_ = 2), then the relevant term in the linear predictor for *θ*_[5,8,1]_ is 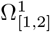, corresponding to the matrix’s first row and second column entry: 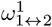 . A positive value for 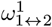 would reflect that male-female dyads tend to interact more than the baseline in group *g*. Because state *k* = 1 does not constitute a directed behaviour (unlike grooming), the male-female and female-male effects are the same. This symmetry is encoded in the off-diagonal entries of **Ω**^1^, and reflected by the double arrows, “↔”.

We follow the same logic for *k* = 2 and *k* ∈ {3, 4}, except that in the latter case, the parameters are allowed to vary depending on the direction of interaction:

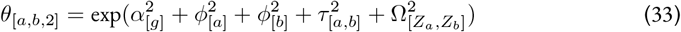

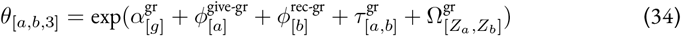

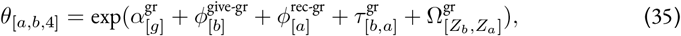

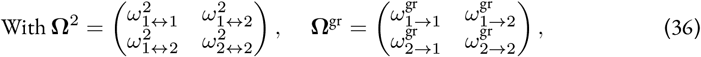

Like state *k* = 1, the state *k* = 2 is undirected and thus, uses a symmetric sex matrix. States *k* = 3 and *k* = 4 are directed, allowing sex effects to vary by direction: *ω*^gr^ (male-to-female) is different from 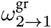 (female-to-male).

To complete this model, we assign prior distributions to the parameters; below, we show only those priors that differ from the core statistical model described in equations 17-21. We need to ensure that the model distinguishes which parameters correspond to the baseline for the short and long sub-states *k* = 1, a problem sometimes referred to as label switching. To do so, we chose priors reflecting plausible time scales and constraints for them. Starting with the short sub-state, we assign:

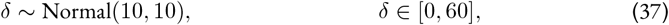

Where *δ* represents time on the natural scale in minutes. This prior constraints the short sub-state to an average of 0 to 60 minutes, with most of the probability density between 0 and 30 minutes. Second, for the longer sub-state, we assign:

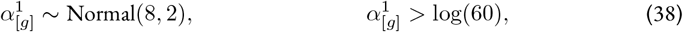

Where *α*^1^ is defined on the log scale. The chosen constraint implies that long sub-state sojourns range from 60 minutes to several days for all groups *g*.

We then assign a prior distribution to *π* that is consistent with our knowledge that any observation *j* in state *k* = 1 is more likely to come from a long compared to a short sub-state. (Recall that for any individual focal-follow protocol, most of the corresponding dyadic observation time consists of no direct behavioural interactions; *i.e*. they are fragments of long sojourns in state *k* = 1). To encode this idea, we use a Beta distribution that concentrates most of the probability density between 0.5 and 1:

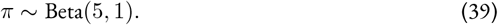

Finally, we assign weakly regularising priors to the parameters capturing the effect of sex:

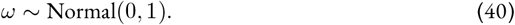

### 3.3. Empirical results

#### 3.3.1 Causal effects

After validating that the extended statistical model described in equations 30-40 can recover the parameters of our generative model (see section S2.1), we updated it with our macaque behavioural data set described in section 2.4. We obtained MCMC draws approximating the posterior distribution of the model’s parameters. We report the corresponding MCMC diagnostics in section S2.2.1. With these posterior samples in hand, we can—at last—discuss the computation of causal effects, our original goal.

An important step in causal inference is to distinguish the causal *estimand* from the corresponding probabilistic causal *estimate*. The estimand is the abstract quantitative goal of an analysis, and ideal target. It can for instance be the “causal effect of sex on the average duration of grooming bouts in a population of Assamese macaques”—though, as we will see below, this statement should preferably be disambiguated with formal notation. The causal estimate, in contrast, is the tangible output of an analysis, an attempt at quantifying the estimand in a given data set. It can for instance consist of MCMC draws, which can be summarised or directly plotted. Thus, the question arises: how should we go from a desired target estimand to a tangible probabilistic estimate?

Let us first discuss how one should *not* go from estimand to estimate. A common misunderstanding in ecology is to conflate the two, by implicitly considering statistical parameters—*e.g*., slopes of multiple regressions—as estimates of causal effects (*e.g*., Gould et al., 2025). Admittedly, the name “effect size” given to regression parameters is not helpful. Yet, this conflation is generally incorrect for at least three reasons: (i) the statistical parameters are often on a different scale than the target estimand; (ii) the estimate often needs to be averaged over study units (*e.g*., individual, dyads); (iii) the estimand might not be identified. Although we have provided causal graphs to conduct causal identification (iii) in our framework—see Figure 10—, we will not discuss this topic in more details here (but see Kawam et al., 2025). Instead, we focus on estimating causal quantities on the right scale (i), factoring in the relevant levels of variations and uncertainty (ii).

We start by defining our causal estimands. We can think of the causal effect of individual sex on social behaviour as a set of *contrasts* between sex categories. These contrasts assess how differently dyads behave across sex combinations, averaging over the other causes of social behaviour. Such effects are called Average Causal Effects (ACE), or marginal causal effects. We compute several ACEs of sex on average holding time, one for each behavioural dimension: no body contact (*k* = 1), body contact (*k* = 2), and grooming (*k* ∈ {3, 4}).

We define the four dyadic sex combinations (*w, z*) as:

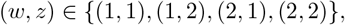

Where 1 denotes male and 2 female. We treat the male-male category (*w, z*) = (1, 1) as the reference group, relative to which causal estimands are defined:

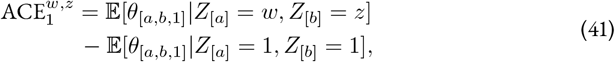

Where 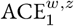 is the ACE of the sex combination (*w, z*) for state *k* = 1. We focus on long sub-states *k* = 1, as we assumed no variation between dyads for short sub-states (equation 30). 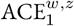 reflects the expected difference in *θ*_[*a,b*,1]_ between dyads of sex combination (*w, z*) and male-male dyads, with the expectation 𝔼 taken over all dyads in the target population that belong to the respective sex categories. Note that we chose not to use causal calculus (*e.g*., Pearl’s *do* operator; Pearl, 2009) to define our estimands, because the estimands are unconfounded and we feared that such novel notation might obscure rather than clarify the concepts for unfamiliar readers.

Let us now return to our extended statistical model (equations 30-40). We observe that no parameter in this model corresponds to 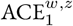. The variation in holding times caused by sex was captured by the parameters Ω that are defined on the log linear scale, not on the scale of *θ*. Furthermore, Ω captures deviations relative to the baseline of social groups, for hypothetical average dyads—which is conceptually and practically different from averaging, or *marginalising* over dyads.

Instead, we can connect our estimand to our statistical model by computing an estimate of 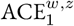 from the model’s joint posterior distribution (Arel-Bundock, 2025; Heiss, 2022; McElreath, 2020; Rohrer, 2024). We start by generating 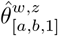, the estimated average time spent in state *k* = 1 by the dyad (*a, b*), keeping the dyad characteristics constant while varying the sex combination to (*w, z*).

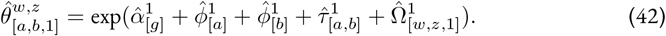

This expression sums the group-, individual-, and dyad-level estimates (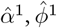, and 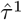, respectively) for the observed dyad, while varying the sex-specific offset 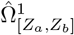. Here, in contrast to equation 31, we manipulate the posterior samples for the parameters 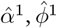, and so on—for instance with R. We obtain 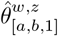 as *output*. In some cases, (*w, z*) matches the observed sexes of individuals *a* and *b*, while in others it does not. Thus, the generated quantities may reflect either factual or counterfactual scenarios.

Using 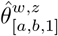, we can estimate the expected holding time in state *k* = 1 for any given sex category, by averaging over all observed dyads, and thus, over the inferred variation in group, individual, and dyad-level characteristics for the population under study. We call these averages 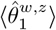:

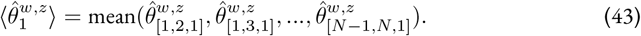

We show the resulting posterior distributions in Figure 11**a**. With these estimates at hand, we can compute the estimated causal effects, AĈE_1_, for different sex combinations (*w, z*) by taking their differences relative to male-male dyads:

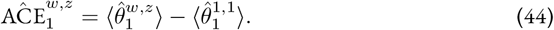

**Figure 11.**
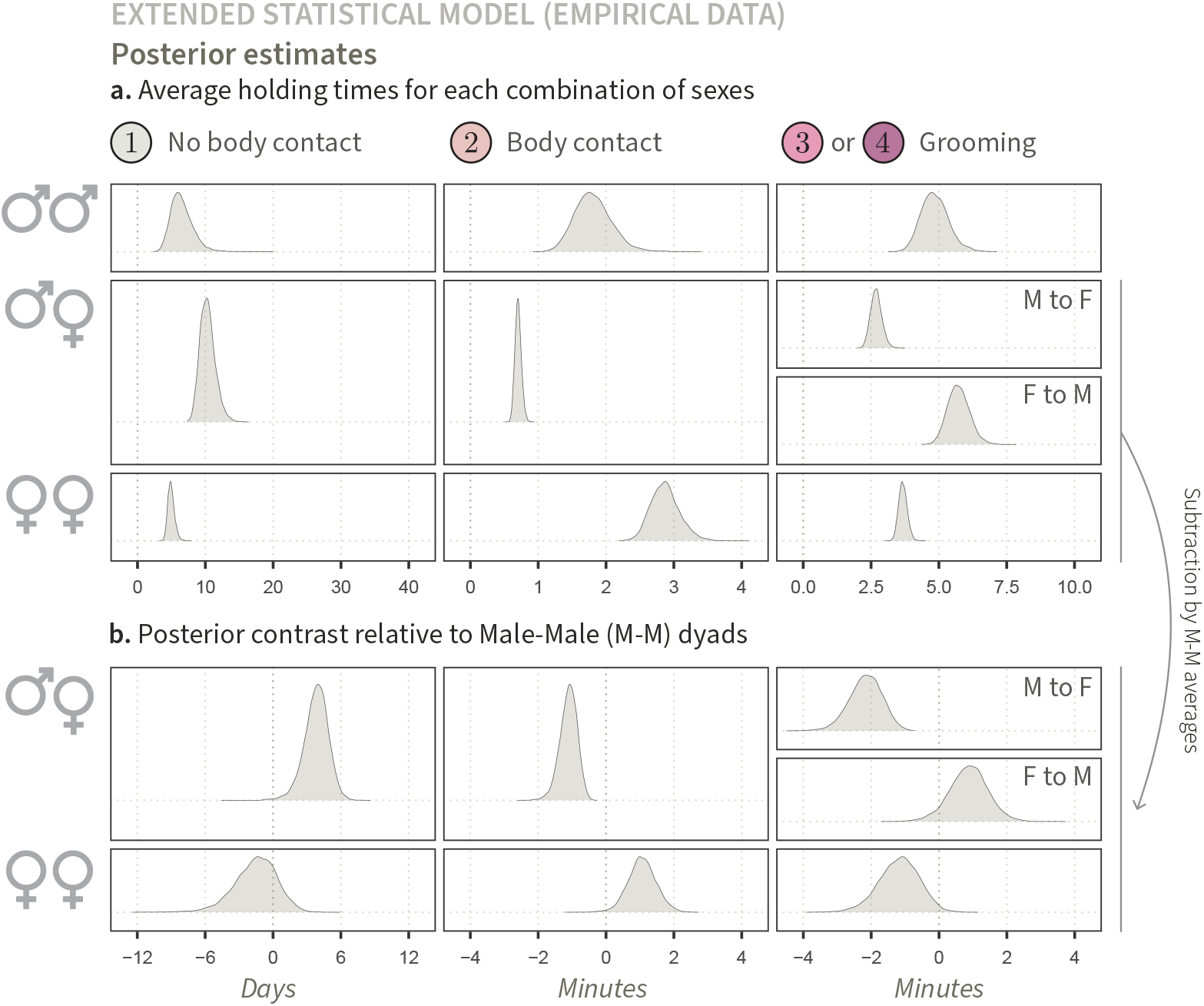
Posterior estimates of average holding times for each combination of sexes andcorrespondingcontrasts (empirical data). (**a**.) Expected holding times across behaviours and sex categories. The horizontal axis indicates the three types of behavioural states, with states *k* = 3 and *k* = 4 (grooming) combined. The vertical axis represents the possible sex combinations, where (*w, z*) = (1, 2) and (*w, z*) = (2, 1) are merged for undirected behaviours. The first column, for instance, shows the estimates 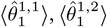, and 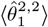. (**b**.) Estimated average treatment effects of sex for each type of behaviour. These causal effects are obtained by subtracting the first-row estimates of panel **a** from the other estimates, column-wise. For instance, to obtain the estimate in the top-left of panel **b**, 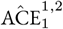, we subtract 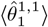 from 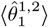. Similarly, the bottom-left estimate, 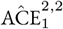, is computed by subtracting 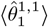 from 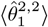.

We followed a similar procedure to compute the average holding times in body contact 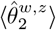, and grooming 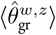, as well as their associated contrasts, 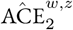, and 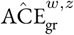. We display the results in Figure 11. Contrary to slope coefficients of multiple regressions and associated *p*-values—which are reported in most empirical studies in the literature—our estimates represent population-level quantitative properties that, first, factor in the heterogeneity between study units, along with their associated inferential uncertainty, and, second, are immediately interpretable in terms of the species’ biology. The panel on the bottom right of Figure 11**a**, for instance, shows the estimated average duration of a grooming bout for a female-female dyad. The width of the distribution reflects the uncertainty associated with this estimate—here, the true value is likely between 3.5 and 3.9 minutes. We compared these rates with those of male-male dyads and found that female-female dyads tend to have shorter grooming bouts on average, with a difference of 0.5 to 2.0 minutes being the most plausible given the available evidence. (Note that, had we modelled an effect of sex on the transition probabilities **Γ**, we could also have compared cumulative grooming duration between sex categories.)

#### 3.3.2. Posterior predictions

To assess whether the modified likelihood of our extended statistical model (equation 30) improves model fit, we perform posterior predictive simulations following the procedure described earlier (section 2.3.2). The only difference is that here, we first draw predicted values for *Q*, indicating whether each predicted sojourn corresponds to a long, or short sub-state:

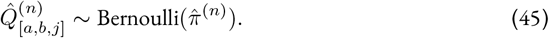

We obtain a predicted value of *Q*, for each predicted observation *j*. Then, we generate predicted holding times *X*:

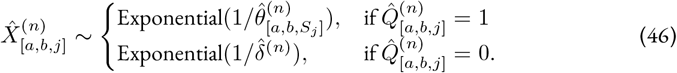

The resulting posterior predictive distributions are shown in Figure 12—one per posterior draw *n*. We observe a clear improvement of the alignment between predicted observations in state *k* = 1 and the empirical data (Figure 12**a**). In particular, we almost fully recover the large number of holding times *X* lasting between 0 and 5 minutes that were previously under-predicted. This supports the idea that even minimal and indirect incorporation of spatial structure in the social network data-generating process is important to predict how dyads engage in direct social interactions over time. Still, the model fails to capture the distribution of holding times between 5 and 40 minutes— highlighting room for further improvement. We return to the issue of spatial structure in the discussion.

**Figure 12.**
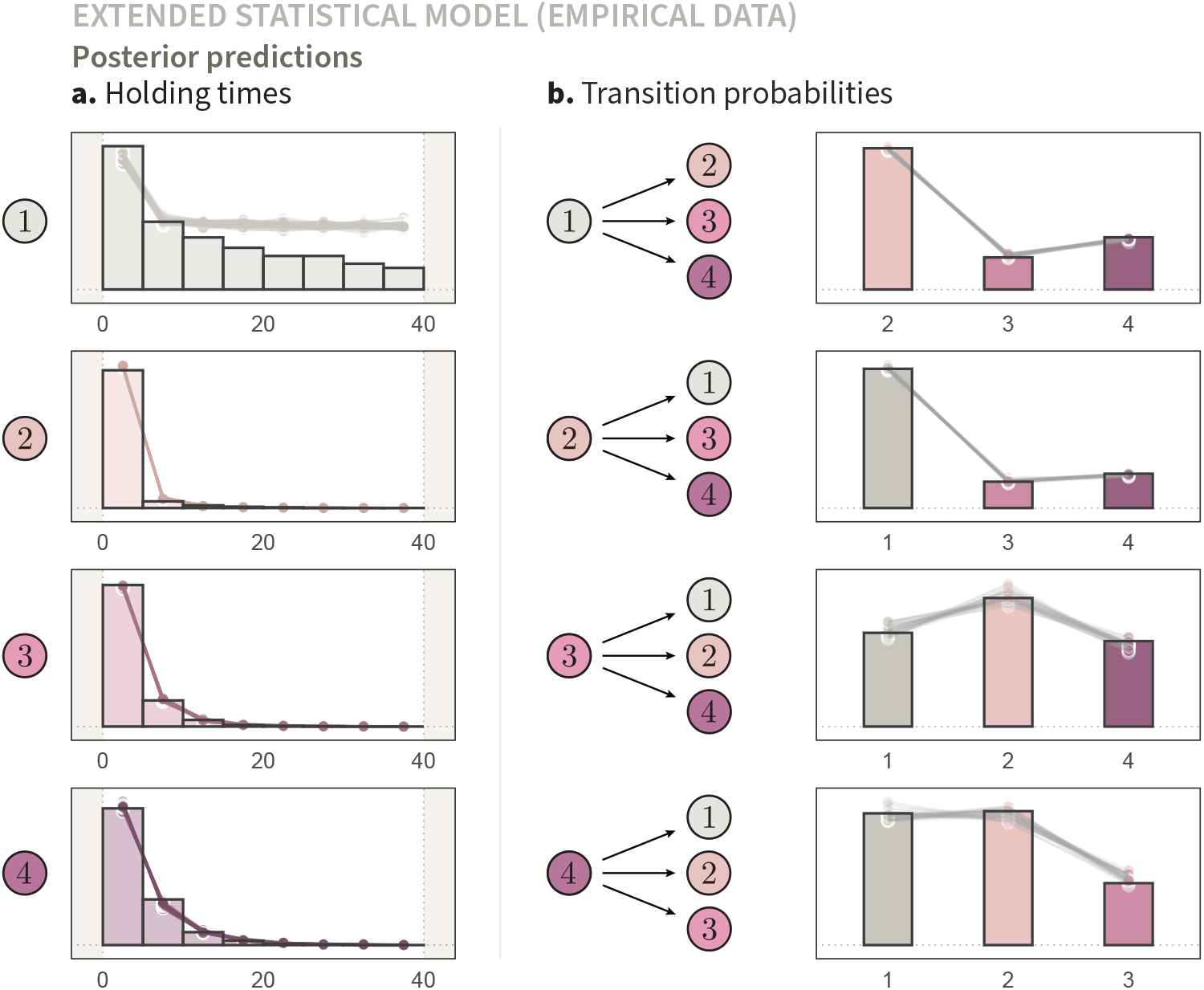
Posterior predictive distribution of the extended statistical model compared to empirical input data. This figure parallels Figures 9, but is based on the extended statistical model described in equations 30-40. Posterior predictions were generated by averaging over short and long holding times in state *k* = 1, as outlined in equations 45-46. We observe an improvement of model fit for holding times in *k* = 1, indicating that a mixture of short and long holding times better approximates this aspect of the empirical data-generating process.

## 4. Discussion

A behavioural ecologist’s work can be thought of as that of a detective (Hilborn & Mangel, 1997). From small fragments of reality, “ecological detectives” attempt to reconstruct a much larger story. In the case of animal societies, behavioural ecologists often work with fragmented behavioural sequences (Figure 1) from which they infer key aspects of their study systems. Their first goal is to generalise, from the noisy and potentially biased observations, to the true patterns of social *interactions* that individual animals exchange with one another—*i.e*., to capture the first level of abstraction of Hinde’s 1976 influential framework. These interactions are not viewed in isolation, but interpreted in the context of social *relationships*, involving different behaviours as well as their patterning in time. The next challenge is to explain the broader *structure* that emerges from these relationships. Moving beyond particular individuals and groups, behavioural ecologists aim to characterise population-level properties. Crucially, making sense of animal social structure comes down to understanding its principles of organisations—*i.e*., to infer its causes, or what Hinde calls “deep structure”. How should ecological detectives assemble their scattered behavioural clues into a coherent causal account of social structure?

Common analytical methods in the field typically involve aggregating the behavioural clues—here, focal-follow data—over time periods (*e.g*., SRI), across behaviours (*e.g*., DSI), and/or across dyads (*e.g*., node strength), to fit standard statistical models. Coefficients and associated *p*-values are then interpreted as evidence for causal hypotheses about the drivers of social structure. However, this approach fails to establish a clear connection between the data and the causal hypothesis of interest. That is, assumptions about the causal process that generated the behavioural observations are usually verbal or implicit, making it unclear whether an analytical strategy works in principle. This disconnect is not only problematic from a theoretical standpoint (we return to this matter below), it leads to practical issues that have received growing attention in recent years (Hart et al., 2023; Kawam et al., 2025; Neumann & Fischer, 2023a; Ross et al., 2023). These issues include poor handling of inferential uncertainty (especially problematic because dyadic social interaction data are often sparse) and confounding (*i.e*. estimates that remain biased regardless of sample size)—both of which can lead to wrong conclusions. Despite growing awareness about problems with current methods in the field (Hart et al., 2022; Redhead et al., 2025), behavioural ecologists have, until now, lacked a framework for causal inference from focal-follow data.

In this manuscript, we have outlined a framework for principled causal analysis of animal social structure from focal-animal sampling data. We began by introducing a generative model that encodes simple biological and measurement processes underlying focal-follow data (section 2.1). This model produces synthetic observations with the structure of empirical data sets. We then designed a Bayesian multilevel statistical model that takes raw (non-aggregated) focal-follow data as input—whether synthetic or empirical—and returns a posterior probability distribution (section 2.2). We validated the statistical model, showing that it could recover the generative parameters and reproduce the distribution of input data when fitted with sparse and censored synthetic observations (section 2.3). Next, we updated the statistical model with an empirical data set collected in wild Assamese macaques (sections 2.4-2.5). As a result, we obtained descriptive population-level inferences about expected durations in several behavioural states, and expected patterns of behavioural sequences over time. We also observed a mismatch between some of our generative assumptions and empirical distributions.

Building on these foundations, we showcased a simple causal inference task by extending the core models (section 3). We first encoded that individual sex shaped social network structure by affecting the durations spent in different behavioural states. In light of the previous mismatch between empirical data and posterior predictions, we also introduced long and short sub-states for the interval between direct social interactions (section 3.1). Next, we designed a statistical model that captures the variation caused by sex and use a likelihood function that marginalises over the unobserved substates (section 3.2). With this statistical model in hand, we turned to computing causal effects using our macaque data set. In particular, we illustrated how empiricists can go from theoretical estimand to tangible estimates, by computing the average treatment effects of sex from the statistical model’s joint posterior distribution (section 3.3). Lastly, we confirmed that the modified statistical model better fitted the empirical data.

More generally, we illustrated a *workflow* for behavioural ecologists to iteratively translate biological hypotheses into formal assumptions; derive, then validate statistical models in light of these assumptions; and finally, compute causal effects of interest from empirical data.

Throughout our manuscript, we emphasised the generative assumptions for the causal processes underlying social interaction data. We did so because these assumptions form the foundations upon which the subsequent statistical models are built and justified (Cartwright, 1995; Deffner et al., 2024; Greenland, 2022; Pearl & Bareinboim, 2014; Woensdregt et al., 2024). That is, the output of an analysis (typically, a causal estimate) is only as plausible as the underlying generative assumptions. If these assumptions are overly unrealistic—*e.g*., the Markov property of our generative model—the resulting inferences will be equally limited. Unrealistic assumptions should thus be critiqued and, eventually, revised. Yet, the first step is to spell them out. It allows researchers to honestly communicate their work to their peers, saying: “If I bring these assumptions in, I get this result out.” An analysis based on overly simple yet explicit assumptions still fares much better than an analysis that is not grounded in any transparent assumptions about the data-generating process at all. In such cases—unfortunately, all too common in the field—, we obtain an output (*e.g*., a statistical estimate) but we do not know under which conditions it can be interpreted, and whether it addresses the hypothesis of interest (Greenland, 2023; McElreath, 2020; Pearl, 2009).

### 4.1. Future extensions

To illustrate our framework, we have focused on a particular system with four behavioural states and specific symmetry constraints for two of these states (Figure 2). This is suited to describing species, like primates, that perform both directed and undirected behaviours. Yet, our framework can be naturally extended to other systems by modifying the number of states and the possible transitions between them. We show three examples in Figure 13. First, our modelling framework could be deployed in a simpler form to model two undirected behavioural states (Figure 13**a**). This includes *k* taking values { “no body contact” }, “body contact”, or { “no body contact”, “play” } . In cases with *K* = 2, dyads would vary in terms of their holding times only, as there would be no variation in transition probabilities. (Recall that with two states in the state-space, there is only one possible transition at the end of a sojourn). Second, extensions of our framework could include *directed transitions* between states in cases where researchers distinguish which individual in a dyad initiates a certain state. When studying mother-infant relationships, behavioural ecologists might, for example, want to know whether body contact is more often initiated by the mother or the infant (Figure 13**b**; *e.g*., Hinde & Atkinson, 1970). This novelty would require modifying the structure of **Γ**. Lastly, our models may provide a foundation to explain more than one type of directed behavioural states unfold over time—for instance, to study the dynamic mechanisms underlying the exchange of grooming bouts and blood-meals in vampire bats (Figure 13**c**; Carter et al., 2024).

**Figure 13.**
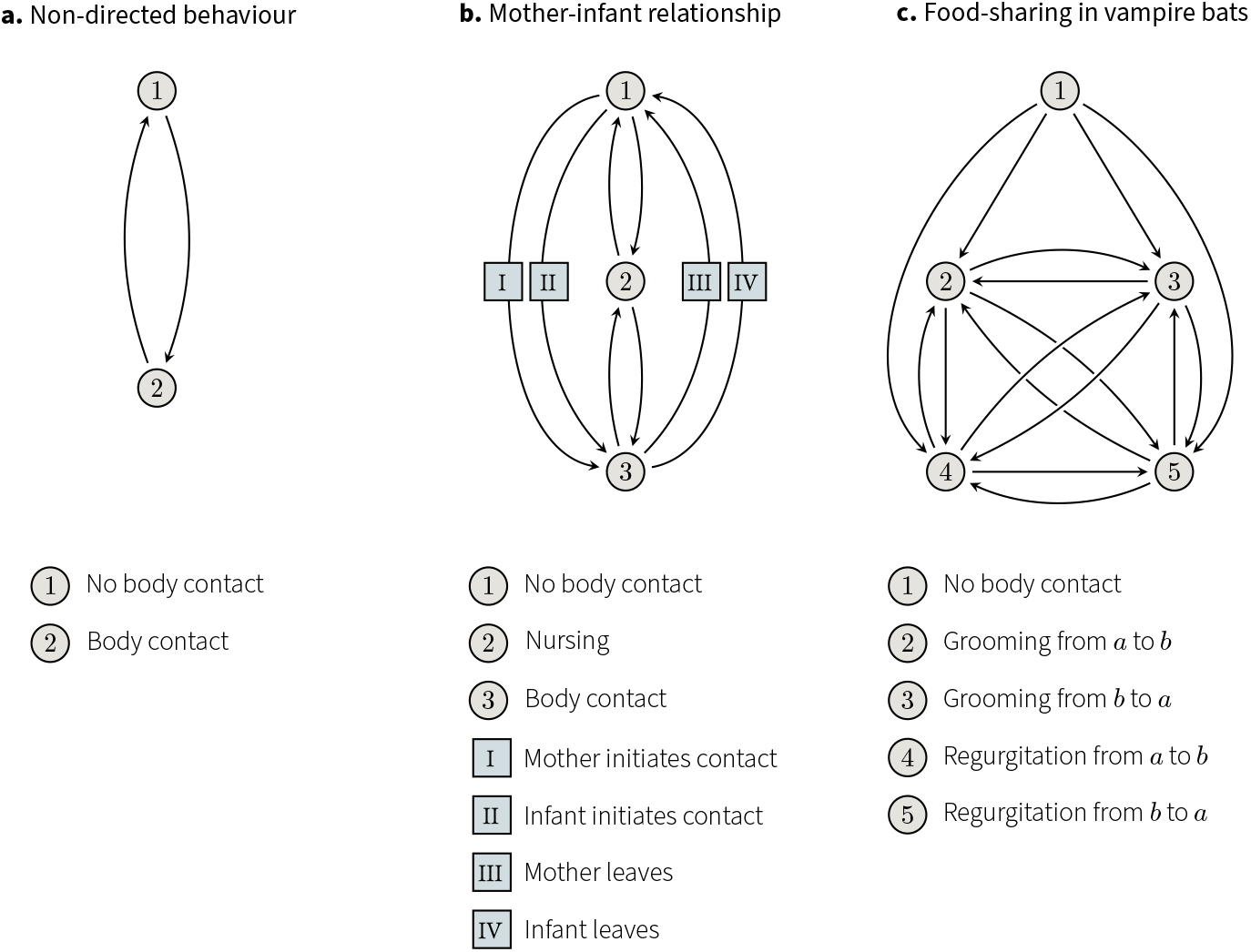
Potential applications of our framework to other social systems. (**a**.) A simpler system with only two non-directed behavioural states. (**b**.) A system where dyads can transition from one state to the next in more than one way. (**c**.) A system with a higher number of states, and thus, a higher number of possible transitions.

Indeed, we believe that one of the most promising directions our framework opens is the study of social relationships over time. Many of the core processes shaping social structure are thought to be intrinsically dynamic. These include processes at the individual level (*e.g*., development, senescence; Arbaiza-Bayona et al., 2025; Sadoughi et al., 2024), dyadic level (*e.g*., reciprocity, dominance relationships; Earl et al., 2025; Leimar & Bshary, 2024; Stranks et al., 2024), and supra-dyadic level (*e.g*., alliance formation, group fission; Lerch et al., 2021; Redhead & von Rueden, 2021; Schülke et al., 2010). Inference in these settings is particularly challenging: one must assess the compatibility between hypothesised dynamic generative processes and empirical data, while accounting for confounding—*e.g*., backdoor paths—not only within but also across time steps (Rohrer & Murayama, 2023; Runge et al., 2023). Additional sources of behavioural variation may be caused by the abiotic environment (*e.g*., weather, seasonality) or the measurement process itself (*e.g*., observer effects)—both of which can vary across observations *j*.

Instead of aggregating their data over arbitrary time windows, researchers are now in a position to model each time-stamped observation directly—enabling more robust inferences about dynamic social processes than was previously possible with common methods in the field. Practically, this involves two steps: first, extending the generative models above by placing parameters *θ* and *γ* on the plate of observations *j* (Figure 4); and second, adapting the statistical models. In simpler cases, *θ* and Γ may be influenced by variables that were *directly measured* such as individual age, time of day, or season. In such cases, researchers may directly add the relevant variables in the statistical model as covariates, like we did with individual sex. Holding times and transition probabilities may alternatively be affected by continuous *latent* variables that must be inferred jointly with the rest of the model’s parameters. Consider, for instance, an individual’s dominance rank. It is an unobserved variable that often changes dynamically as the individual engages in agonistic interactions (Neumann & Fischer, 2023b). This latent variable could be directly estimated alongside *θ* and Γ over time, allowing us to study its effects on affiliative behavioural interactions, even if an individual’s rank is often uncertain at any given point of time.

Modelling temporally resolved observations opens the door to capturing not only the effects of timevarying factors such as age or dominance rank. It also—and, perhaps, most importantly—, allows researchers to study the feedback loops between these variables and social structure by tracking the “(co)evolution” of social network structure and individual phenotype, in a manner similar to stochastic actor-oriented models (Kalish, 2020; Snijders, 2017). (Note that “evolution” here loosely refers to dynamic change over time, distinct from its specific use in evolutionary biology.) For example, suppose that as individuals develop and age, their social relationships change. In turn, individuals’ social relationships may impact their probability of survival, and with it, the age structure of the population (Ostner & Schülke, 2018; Snyder-Mackler et al., 2020). Similarly, an individual’s high dominance rank might increase its attractiveness, shaping the affiliative relationships it forms, which in turn may affect its position in the dominance hierarchy (Schülke et al., 2010). Such feedback loops may also involve reproductive state, hormonal changes, or the affiliative social network itself. Let us return to dyadic reciprocity: a process where what an individual, *a*, did to another individual, *b*, in the past influences how *b* behaves toward *a* in the present. Within our framework, this could be represented by past sojourns (*e.g*., state *S*) affecting the parameters governing current sojourns (*i.e*., Γ or *θ*). Inferences in these cases are complicated by the fact that the relevant past behaviours are often unobserved (*e.g*., “was the last blood-meal regurgitation given from *a* to *b*, or from *b* to *a*?”; Figure 13). Fortunately, the marginalisation technique introduced in equation 30 for the two substates *k* = 1 could be directly applied to this context as well—*i.e*., by averaging, not over possible sojourn types, but over possible past states.

A second key direction for extension involves *space*. The decisions of individual animals to socially interact with conspecifics are generally coupled with spatial decisions about where to go, and thus, generative models that ignore spatial behaviour will sometimes be overly unrealistic. Moreover, spatial co-occurrence is sometimes the best—or only—available evidence from which researchers can infer the social relationships between individuals (*e.g*., in cetaceans, fish, ungulates; Albery et al., 2022; Atton et al., 2014; Gerber et al., 2022), or to study the spread of infectious diseases between them (Krause et al., 2015).

For these reasons, we foresee—or at least, hope—that empirical studies of social network structure will increasingly incorporate spatial behaviour in both causal and statistical models. First, researchers may build generative models in which social interactions unfold in an explicit two-dimensional space (*i.e*., “agent based” models; *e.g*., Puga-Gonzalez et al., 2009, 2015). As opposed to the generative model presented in section 2.1 where all interactions were purely dyadic, such agent-based models could automatically include higher-order interactions. For instance, if an individual *a* approaches another individual *b*, and *b* is in proximity with *c*, then *a* would automatically be in proximity with *c* as well. (This feature could not be directly implemented in the generative model of section 2.1 without substantial alterations. Thus, although it may still be a sensible modelling choice in specific cases, we do not generally advise that researchers naively define “spatial proximity” as a state *k* in the dyadic state-spaces.) Spatially-explicit generative models should further include the measurement process: *e.g*., scan sampling, focal-follows, or gambit of the group (Franks et al., 2010; Ginsberg & Young, 1992; Whitehead, 2008). Regarding statistical modelling, we anticipate that incorporating the higher-order structures that result from spatial associations may be partly achieved with varying-effects (*e.g*., Bonnell et al., 2024) and hypergraphs (Battiston et al., 2021). However, the practical integration of such modelling tools into a coherent framework for causal inference remains largely to be worked out by future research.

The causal study of animal social structure from focal individual data presents a formidable challenge. The interface between causal hypotheses and empirical datasets is thick, and laden with strong assumptions. Yet, in this manuscript, we make a case for optimism. We lay out the practical groundwork for researchers to unpack their assumptions and connect them to an analytical strategy. In doing so, we hope to stimulate a discussion on the assumptions themselves—leading to clarification and improvement of theory; to justify statistical procedures on clear theoretical grounds—leading to an improvement of methods; and, more generally, to provide continuity between theoretical and empirical research in behavioural ecology.

## Data and code

All relevant code and data can be found on this GitHub repository.

## Acknowledgments

We wish to thank Marijtje van Duijn, Alice Hill, and Ana Lucia Arbaiza Bayona for their comments on earlier versions of this work, as well as Bob Carpenter for his help on the Stan forum. Sofia M. Pereira created the watercolor paintings of Figure 2 with the kind help of Judith von Nordheim.

## Figures

The figures in this manuscript were generated using *R* (version 4.2.1), LaTeX (TikZ package), *Adobe Illustrator CC 2015*, or a combination of them. In R, we used the following packages: *ggplot2* (Wickham, 2011), *tidybayes* (Kay, 2020), *patchwork* (Pedersen, 2019), and *ggraph* (Pedersen et al., 2017).

## Ethics and permits

The empirical data used in this manuscript were collected non-invasively, using protocols that adhere to the Association for the Study of Animal Behaviour (ASAB) guidelines for the Use of Animals in Research. The study was further authorised by the Department of National Parks, Wildlife and Plant Conservation of Thailand, and the National Research Council of Thailand under a benefit-sharing agreement, with permit numbers 0002/2424 (April 23^rd^, 2014) and 0002/470 (January 26^th^, 2016).

## Supplementary Materials

### S1. Core models

### S1.1. Generative model

#### S1.1.1. Plate notation

In this section, we provide a short visual introduction to the plate notation for hierarchical graphical structure.

Consider a group of three individuals (*N* = 3), forming 3 dyads (*a, b*): (1, 2), (1, 3), and (2, 3). Suppose further that we study the average holding time *θ* of two different states (*K* = 2), and that the parameters *θ* are affected by a fixed, dyad-specific feature *R* (*e.g*., genetic relatedness).

We represent this hierarchical system in Figure S1**a**. The outer-plate loops over pairs of individuals (*a, b*), while the inner-plate loops over states *k*. To understand what we mean by “looping over” units, let us break it down step by step. In Figure S1**b**, we decompose the outer-plate by explicitly representing each dyad. We further decompose the repetition over *k* in Figure S1**c**, distinguishing the two random variables as separate nodes. The graph at the top of Figure S1**c** may for example illustrate that genetic relatedness between two specific individuals, (1, 2), influences the time they spend in two behavioural states: 1 (*e.g*., no body contact) and 2 (*e.g*., body contact). The compaction of these repeats through plates allow us to scale their representation for any number of individuals and states.

For an introduction to plates and other *template-based* graphical representations, we recommend Chapter 6 of Koller and Friedman (2009).

**Figure S1.**
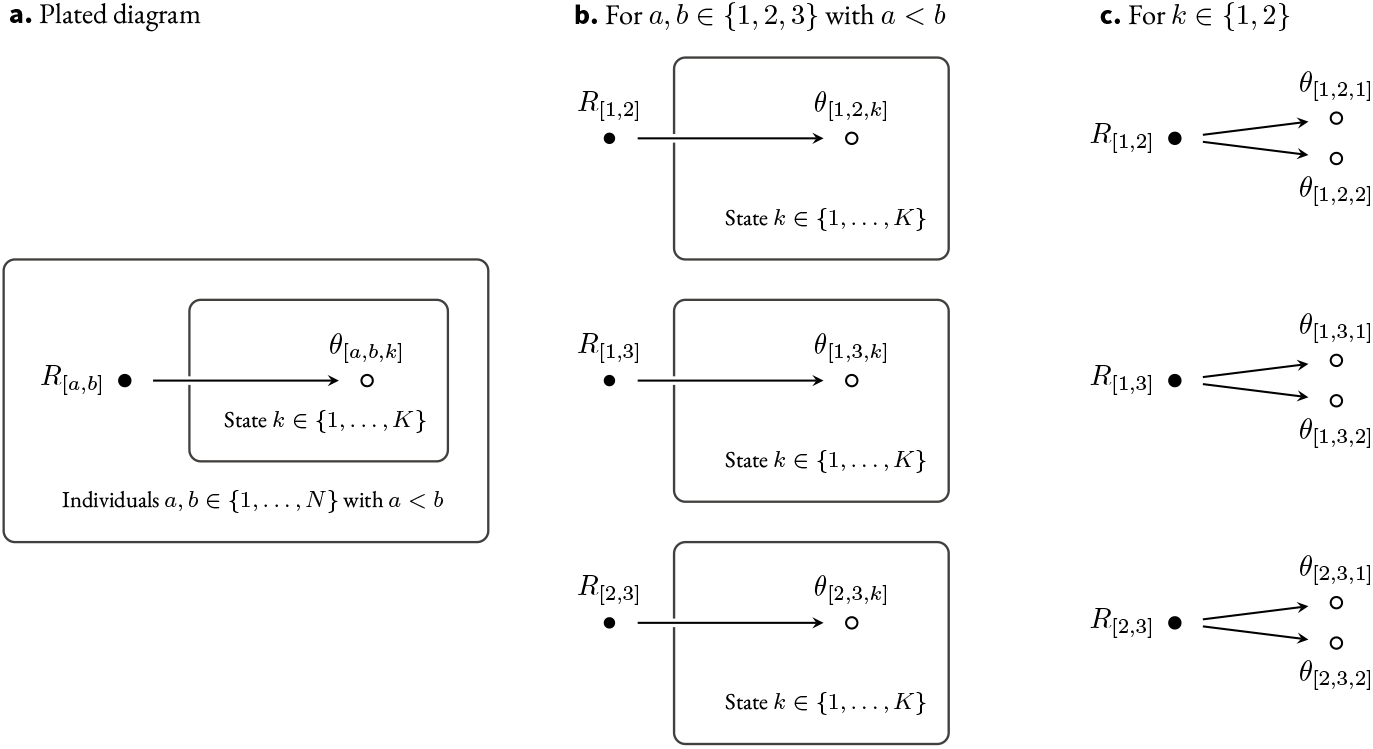
Decomposition of a plated graph.

#### S1.1.2. Pseudocode for the Continuous-Time Markov Chain

##### Algorithm 1

Generate a sequence of state sojourns *i* for each dyad (*a, b*). *t*^max^ represent the maximal time after which the simulation is stopped. Each observation *i* corresponds to a sojourn in a given state. It is characterised by a state number *S*, a true holding time *X*^true^, a starting time *t*^start^, and an ending time *t*^end^. Note that observations *i* are the full sequence of dyadic states, not only those that are observed through focal-follows.

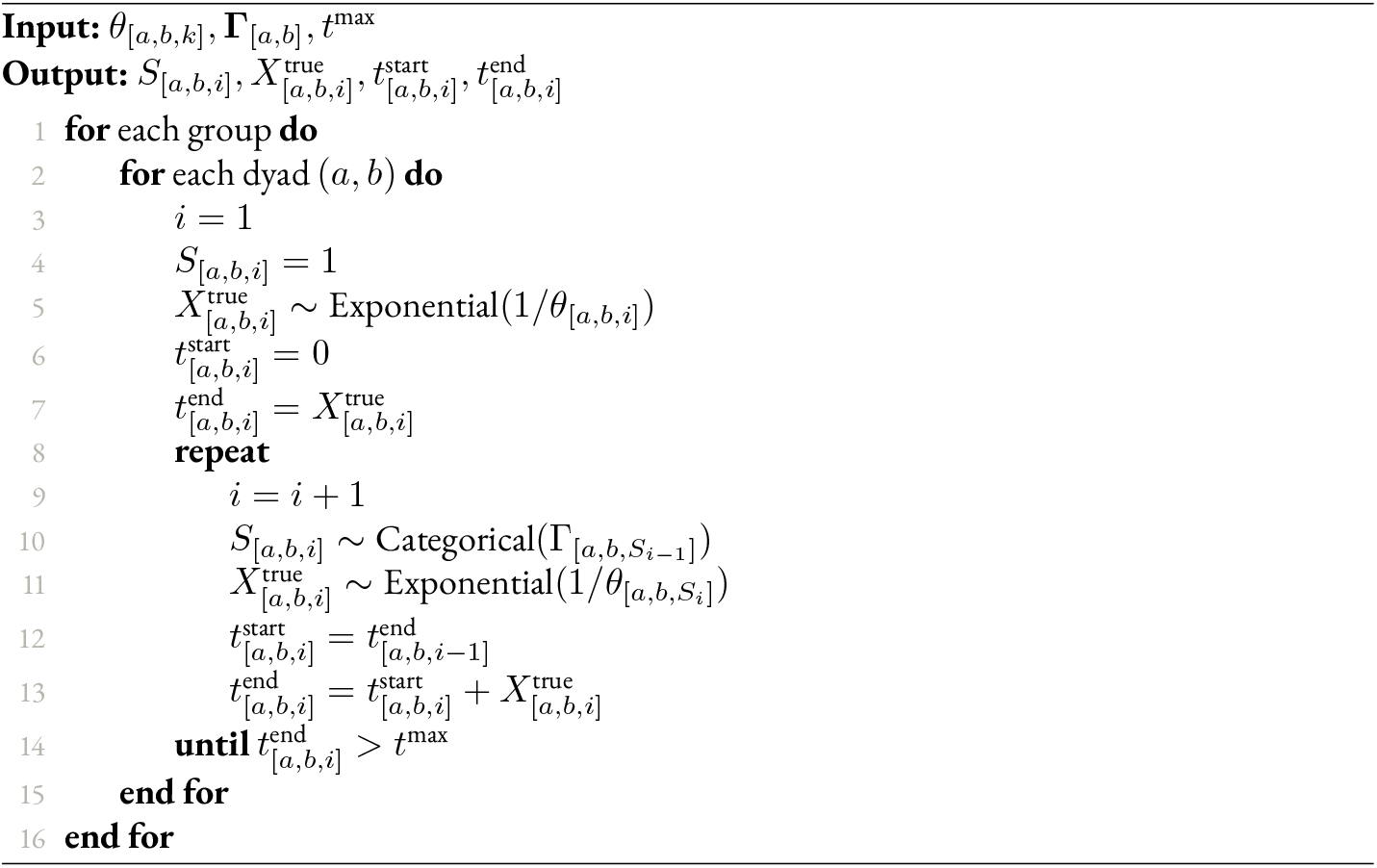

#### S1.1.3. Pseudocode for individual focal-follows

##### Algorithm 2

The state sojourns *i* generated with **Algorithm 1** are stochastically sampled and censored through focal-animal sampling protocols, generating observations *j*. `start_times` is a group-specific vector of focal-follow protocols’ starting times *t*, spaced from one another by 40 minutes. It typically starts at a high first value (*e.g*., `start_times` = {10000, 10040, 10080, … }, to ensure that the state of the first observation *j* is independent of the initial state. The `prot_id` vector takes on a unique value for each dyadic focal-follow protocol, *i.e*. its value is shared by all observations *j* that are part of the same focal-follow protocol.

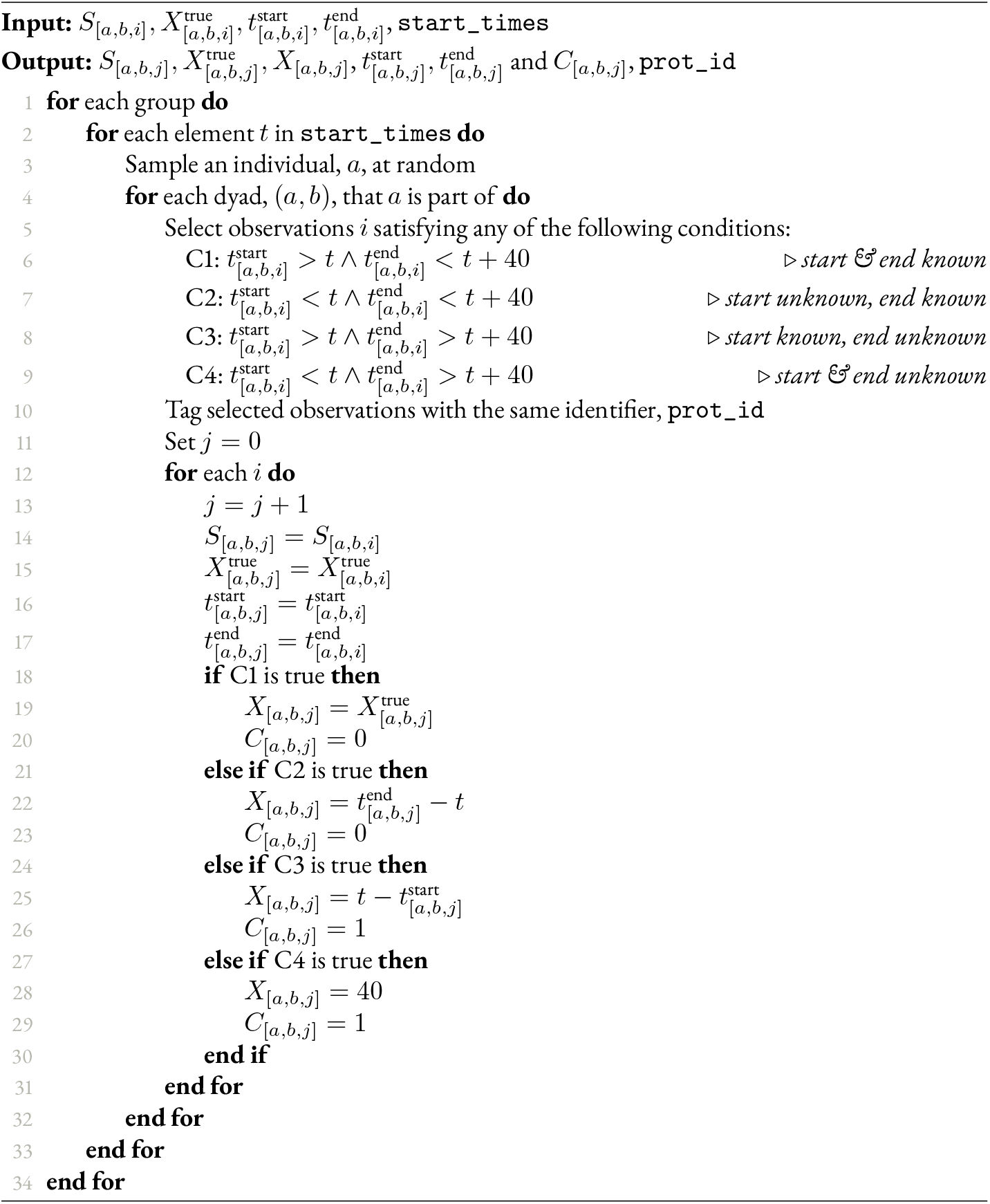

#### S1.1.4. Generative parameters

**Figure S2.**
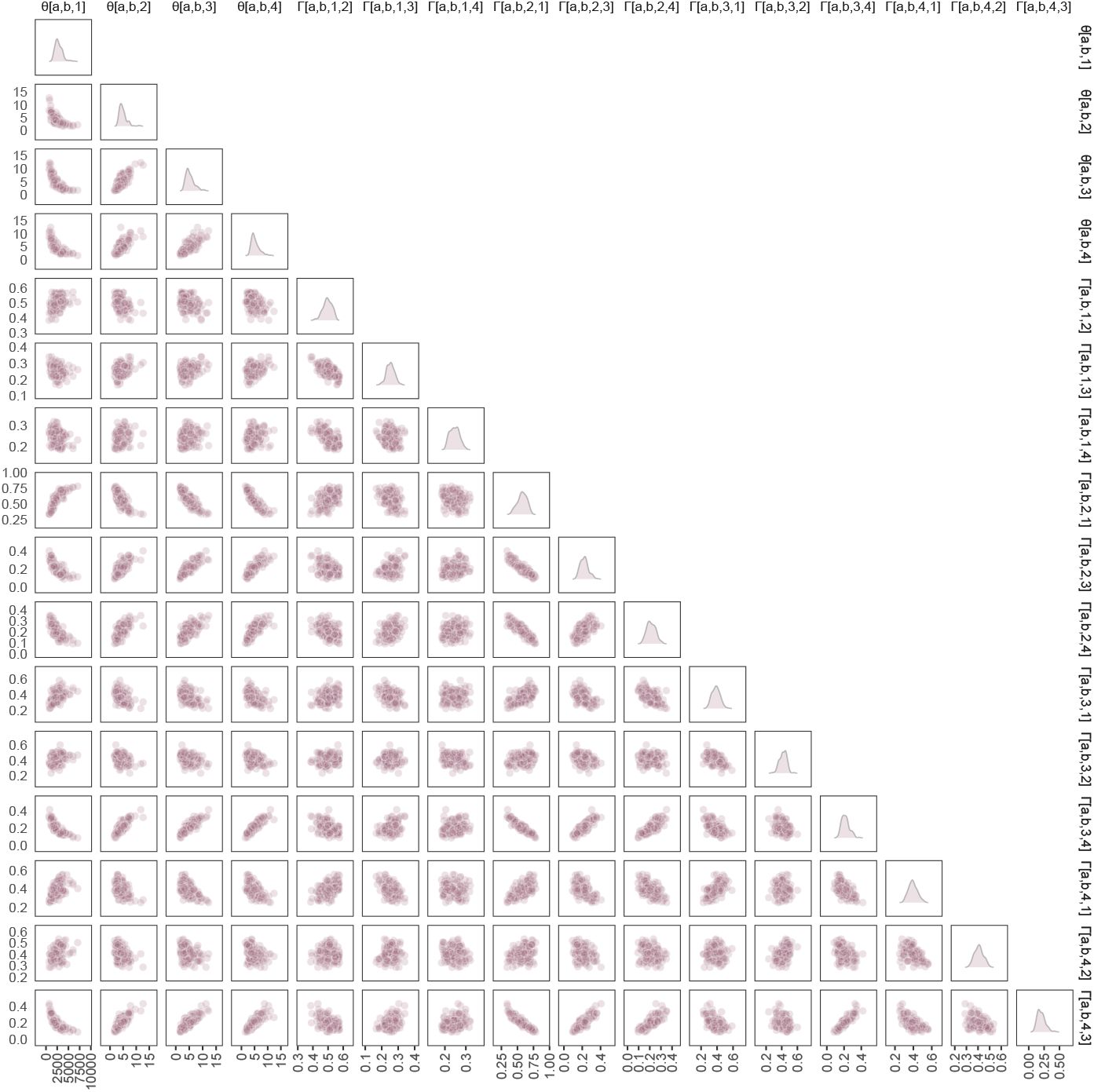
Dyadic parameters θ_[a,b,k]_ and Γ[a,b,k,l] for three groups of 10 individuals (135 dyads in total) and for k, l ∈ {1, 2, 3, 4} . The parameters’ marginal distribution is shown on the diagonal; their pairwise associations, on the scatter plots.

#### S1.1.5. Distributions of synthetic data

**Figure S3.**
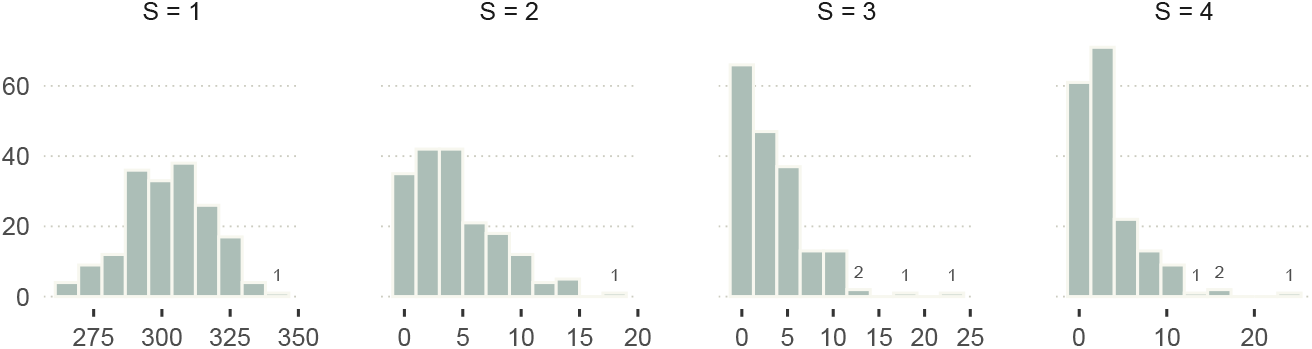
Number of observations j per dyad and per state k, for one iteration of our core generative model.

**Figure S4.**
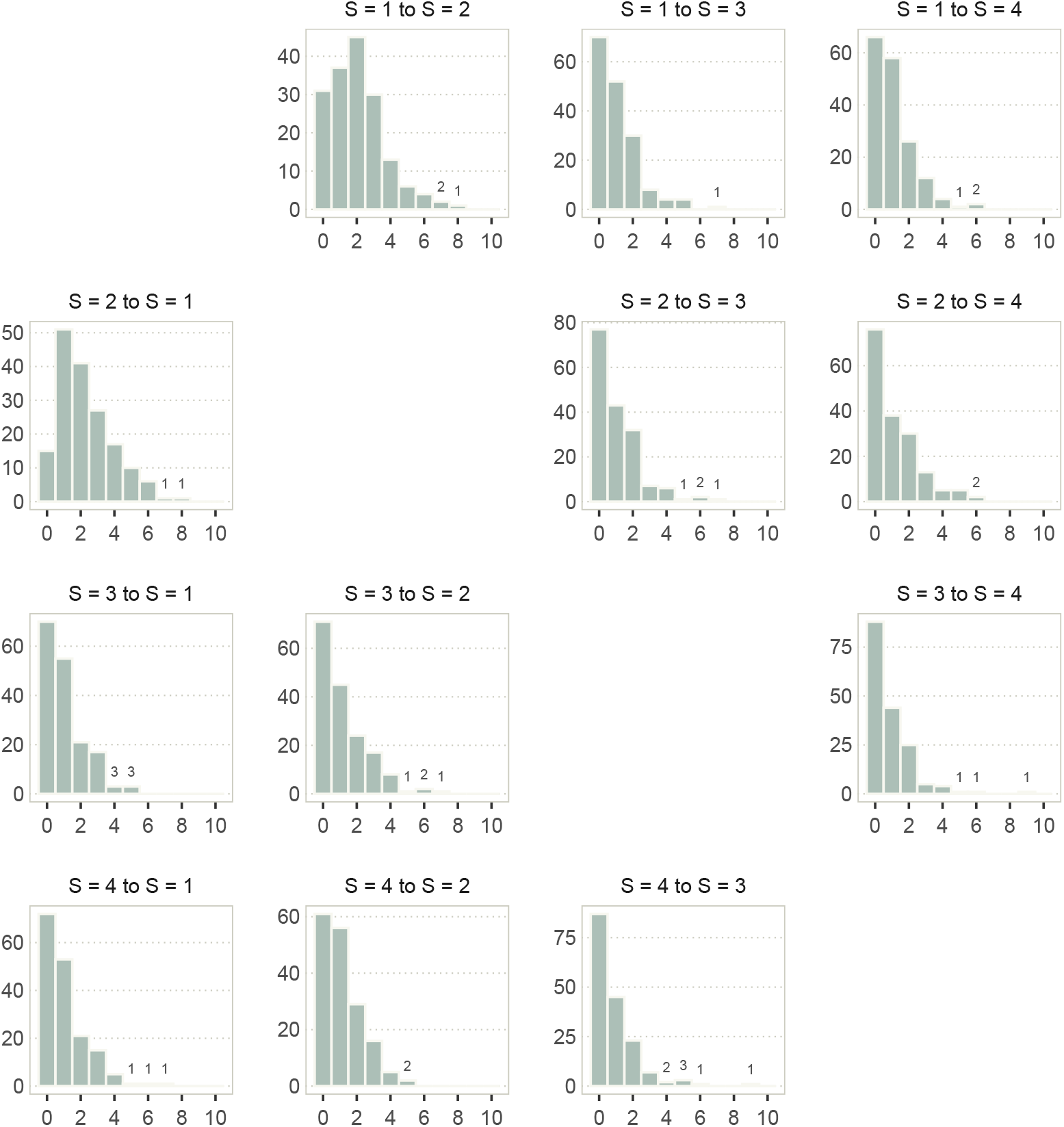
Number of observations transitions per dyad and per combination of current state k and future state l, for one iteration of our core generative model.

### S1.2. Statistical model

#### S1.2.1. Model definition

**Likelihood for holding times**

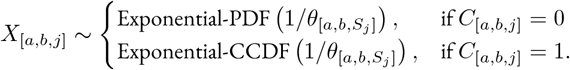

**Linear models**

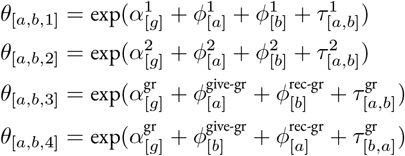

**Likelihood for state transitions**

S_[*a,b,j*+1]_ ∼ Categorical 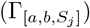, for *j* and *j* + 1 belonging to the same focal-follow protocol,

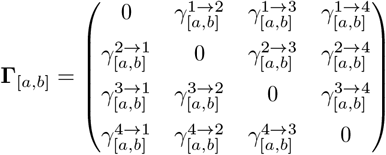

**Linear models and latent scores Ψ**

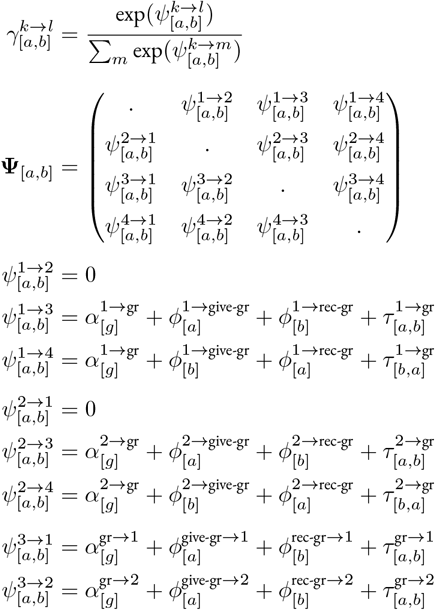

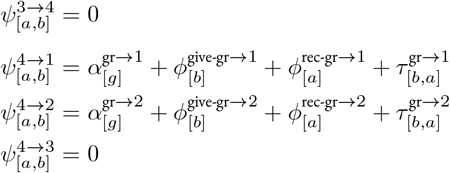

**Adaptive priors**

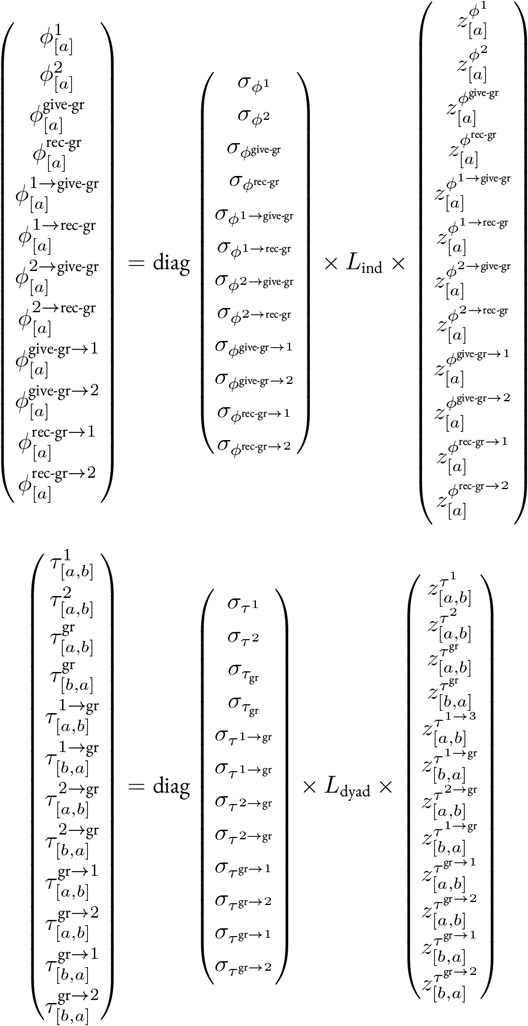

**Hyper priors**

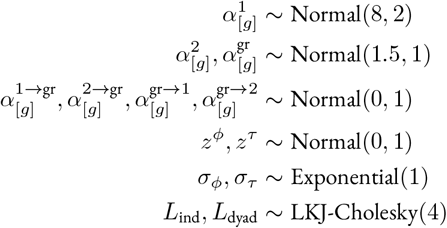

#### S1.2.2. Exponential PDF and CCDF

In this section, we briefly introduce the Exponential distribution and its different forms: the so-called PDF, CDF, and CCDF. In doing so, we hope to provide some guidance for readers who are unfamiliar with time-to-event modelling, so that they can make sense of the likelihood function described in section 2.2.

The Exponential Probability Density Function, or PDF, is given by the following expression:

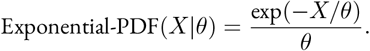

The function, explicitly given on the right-hand side of the equal sign, describes the curve of Figure S5**a**. This curve provides a measure called probability density for the following question:

*Q1: How likely is it that a waiting time takes on a certain value X?*

With the Exponential distribution, the most likely values are always close to *X* = 0 and decrease as *X* increases. How fast this curve decreases is controlled by *θ*, the distribution’s mean—or, alternatively, by its rate *λ*, defined as one over *θ*.

When we attempt to *infer θ* from observed waiting times *X* until an event happens, we flip the problem around, and assess how compatible observed waiting times are with different values of *θ*. We do so by defining the likelihood function of a statistical model using an Exponential PDF. This is what we did in section 2.2 to describe non-censored holding times—*i.e*., waiting times until the end of the states (the “events”) happen.

Suppose now that instead of asking how likely it is that the event happens at time *X* (*Q1*), we ask:

**Figure S5.**
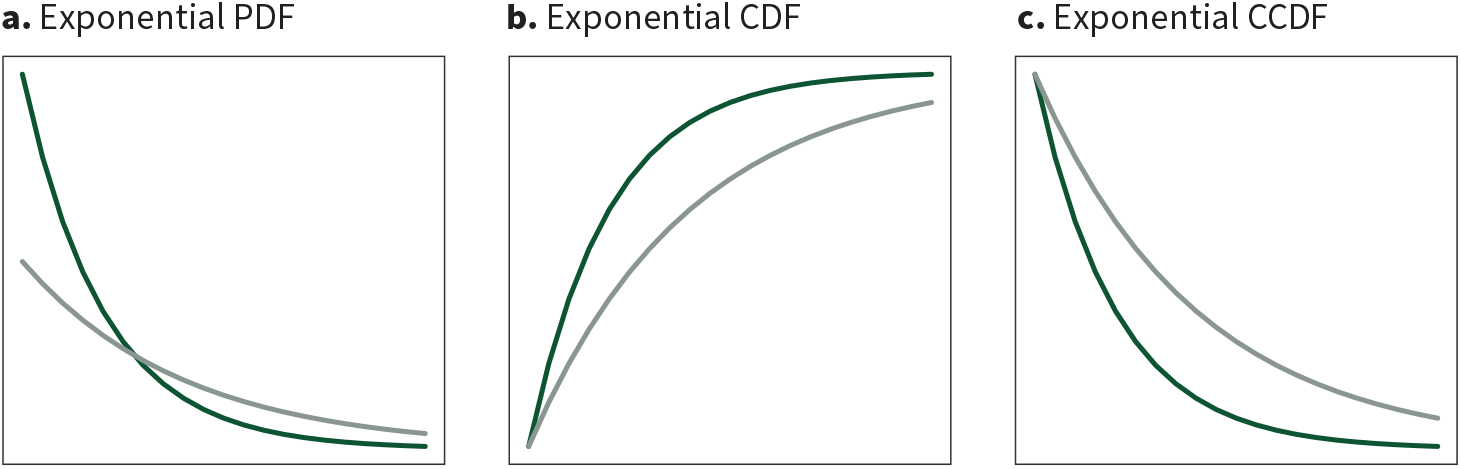
The different forms of the Exponential distribution. Probability density, on the y-axis, is shown as a function of *X*, on the x-axis. The two colours correspond to different values of *θ*—the distribution in light green has a mean twice as large as the one in dark green.

*Q2: How likely is it that after a certain amount of time X, the event has already happened?*

*Q1* and *Q2* may sound similar, but they are critically different. Close to *X* = 0, we should have a low probability that the event has already happened—after all, we have just started to wait. As time passes, the probability that the event has already happened should increase, then plateau. This pattern will arise even when the most likely waiting times are close to 0 (like in the case of the Exponential PDF). Our second question can be addressed by measuring the *area* under the PDF, from *X* = 0 to any other value *X* = *x*. The function to achieve this is called the Exponential Cumulative Density Function, or CDF.

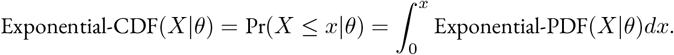

In English: the Exponential CDF for a certain value of *X* given the parameter *θ* is equal to the probability that the waiting time *X* is smaller or equal to a certain value *x*, and is described by the integral of the PDF taken between 0 and *x*. We plot the CDF in Figure S5**b**.

When modelling holding times in our network models, a number of sojourns are censored. For these observations, we know (i) that we have waited for a certain amount of time *X*, and (ii) that after this time, the event has not happened yet. When we wish to infer *θ*—that is, to know how compatible the censored waiting times are with *θ*—the question that we ask is thus closely related to that of the CDF. Here, we do not wish to know what is the probability that the event has already happened (*Q2*), but instead:

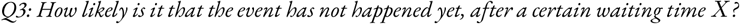

This quantity is simply given by 1 minus the CDF, a function otherwise known as the Complementary CDF, or CCDF:

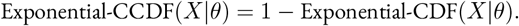

We show the Exponential CCDF in Figure S5**c**. For the reasons above, we modelled the non-censored sojourns (referring to *Q1*) using the Exponential PDF, and the censored ones (referring to *Q3*) with its CCDF. Note that these considerations apply to other possible choices of distribution (*e.g*., the Gamma distribution).

#### S1.2.3. Validation of the prior model

**Figure S6.**
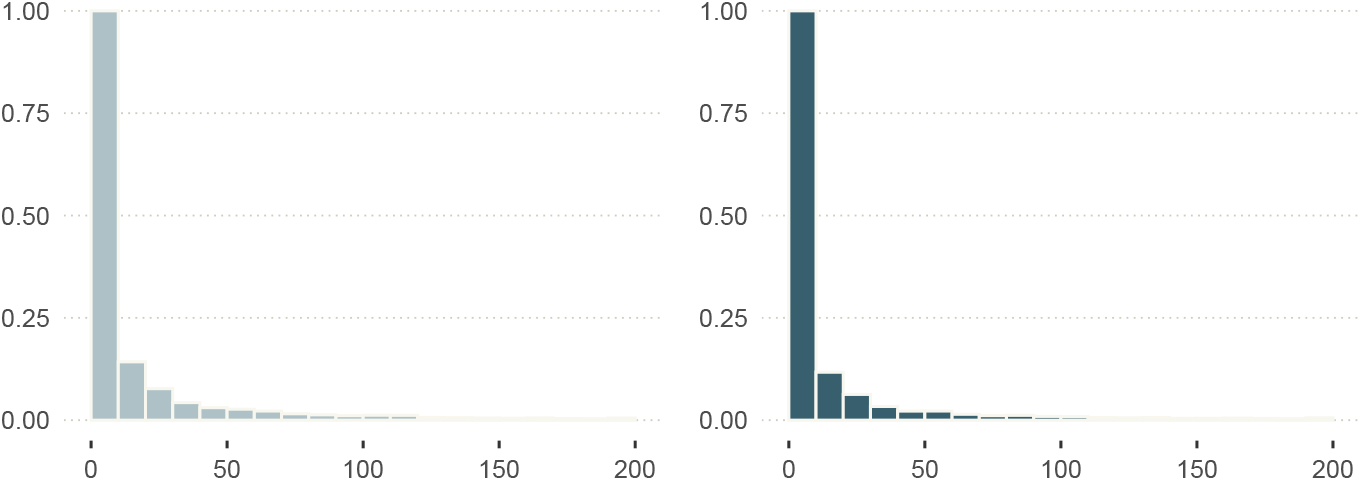
Prior probability distribution of θ_[a,b,1]_ (left), and prior predictive distribution of X_[a,b,j]_ | S[a,b,j] = 1 (right). The x-axis shows average holding times (left), and realised holding times (right) *in days*, for any pair of individuals (*a, b*). The y-axis represents probability density, approximated by 4000 MCMC samples.

**Figure S7.**
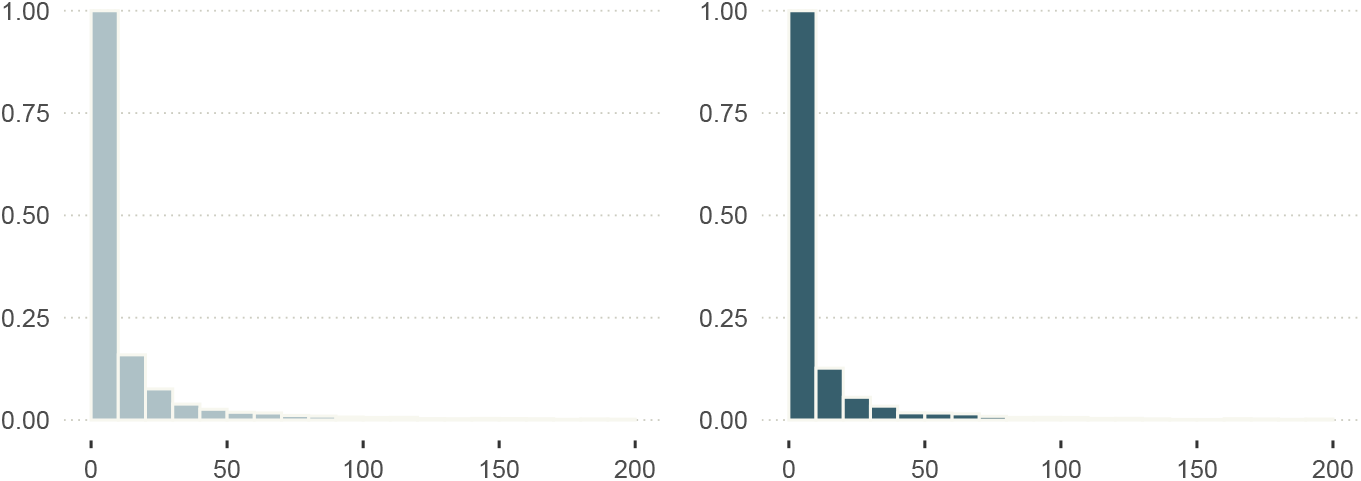
Prior probability distribution of θ_[a,b,k]_ with k ∈ {2, 3, 4} (left), and prior predictive distribution of X[a,b,j] | S_[a,b,j]_ ∈ {2, 3, 4} (left). The x-axis shows average holding times (left), and realised holding times (right) *in minutes*, for any pair of individuals (*a, b*). The y-axis represents probability density, approximated by 4000 MCMC samples.

**Figure S8.**
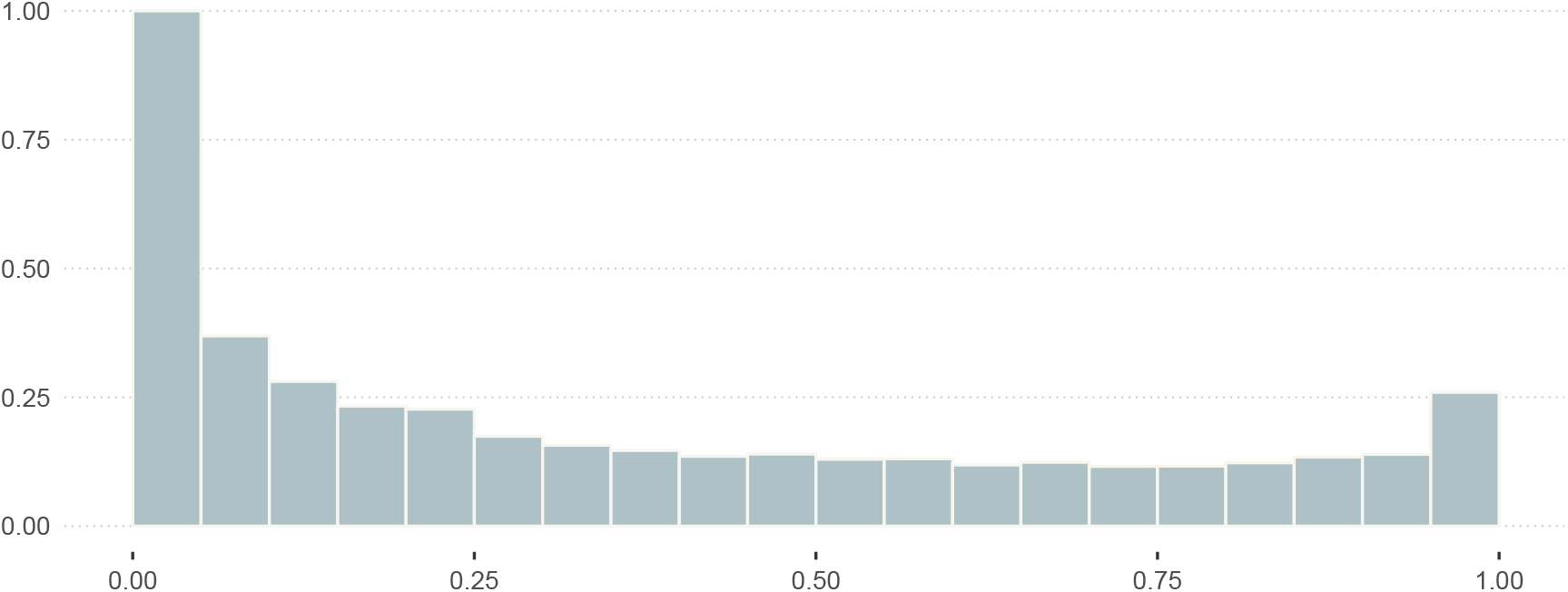
Prior probability distribution of Γ_[a,b,k,l]_, for any pair of individuals (a, b), and for combination of states k and l such that k ≠ l. The y-axis represents probability density, approximated by 4000 MCMC samples.

#### S1.2.4. Validation of the posterior distribution

**Figure S9.**
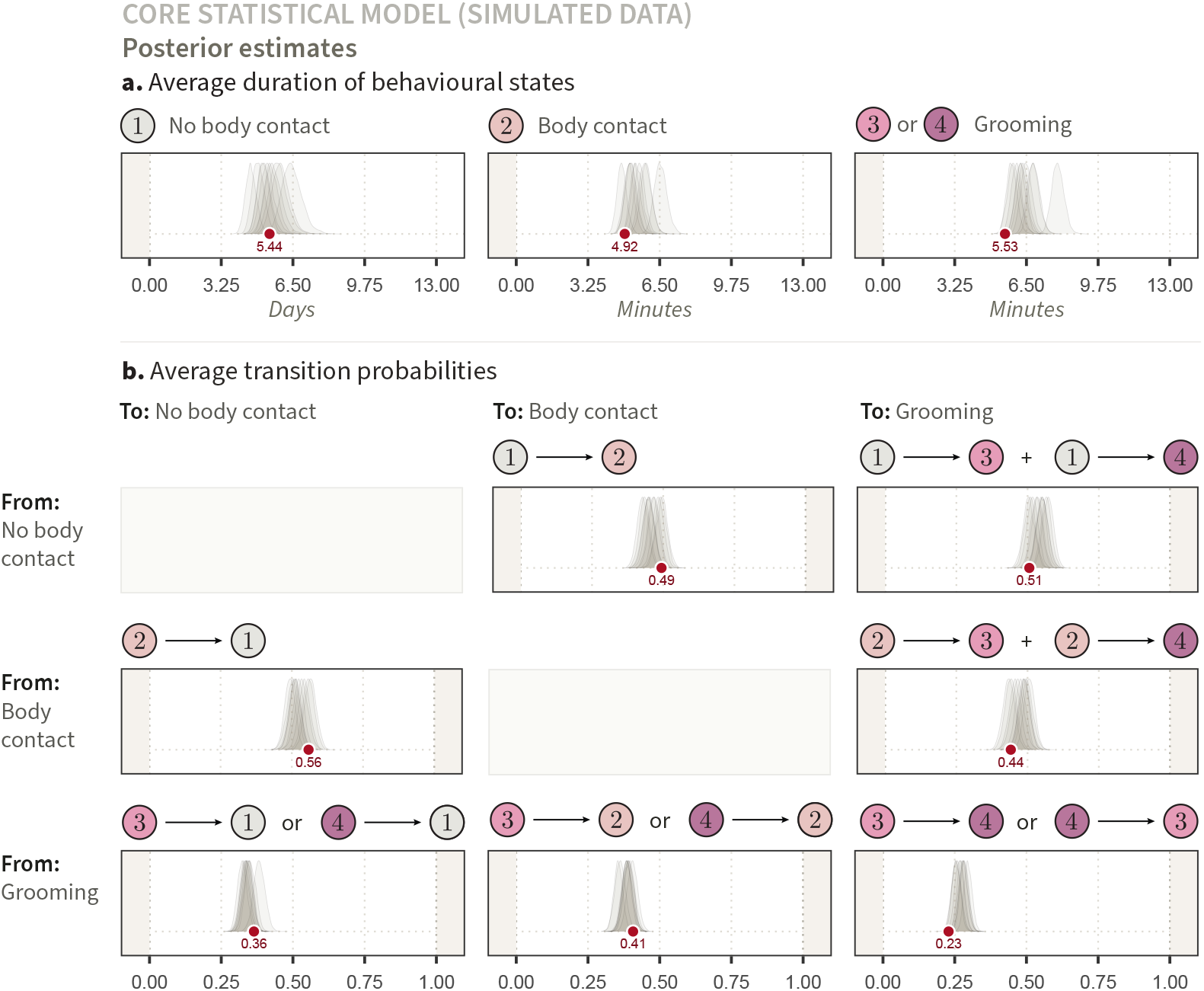
Posterior estimates of average holding times and transition probabilities (simulated data). The true parameter values, represented as red points, are derived from the generative model introduced in section 2.1. The marginal posterior probability distributions were obtained by updating the statistical model described in section 2.2 with ten data sets resulting from ten iterations of the generative model. **(a.)** Each marginal posterior distribution corresponds to an estimated average, over all dyads, of *θ*_[*a,b,k*]_. The numbers in coloured circles indicate the states *k* that were considered. On the right hand side, state “3 *or* 4” means that we appended *θ*_[*a,b*,3]_ and *θ*_[*a,b*,4]_ for all dyads (*a, b*), and took the average of the resulting vector to compute the average grooming bout duration. **(b.)** Each marginal posterior distribution corresponds to an average, over all dyads, of transition probabilities 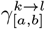. Again, the numbers in coloured circles indicate the states *k* and *l* that were considered. Where two pairs of states are connected by a “+” sign (*e.g*., on the top-right), it means that we summed the inferred average (*e.g*., average transition from state 1 to state 3, plus average transition from state 1 to state 4, for the transition from no body contact to grooming). Transitions connected by an “or” sign imply, like in panel **a**, that we appended the two vectors and took the average of the resulting vector—*e.g*., on the bottom-right, we computed the average probability of immediate reciprocation, whether from state 3 to state 4, or from state 4 to state 3.

### S1.3. Empirical data

**Figure S10.**
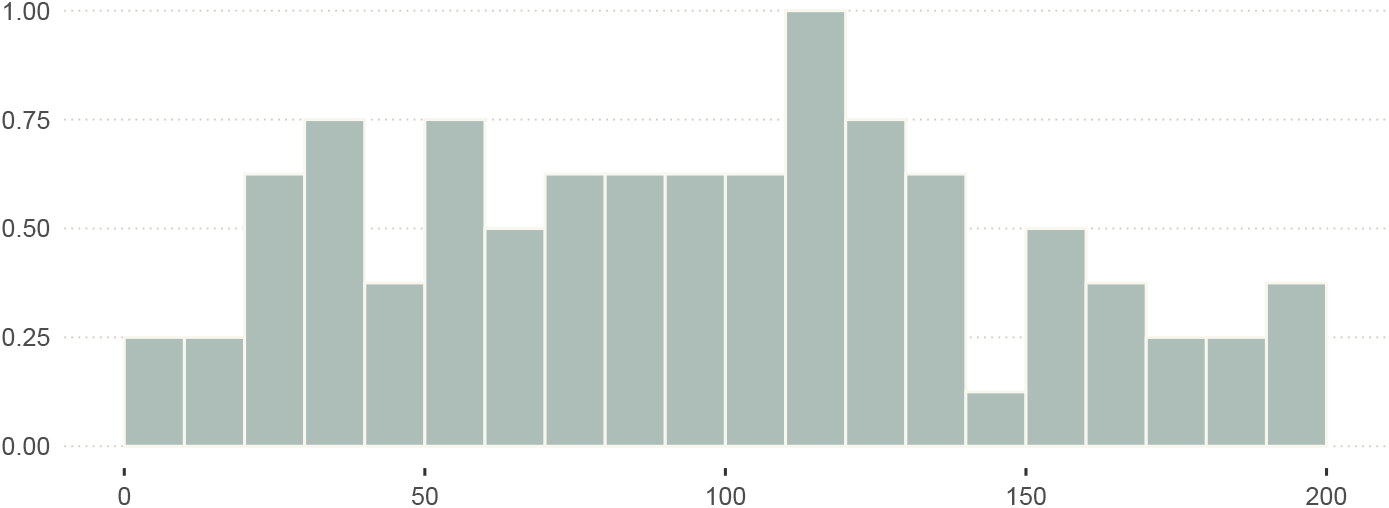
Empirical distribution of sampling effort (hours) across all individual animals.

**Figure S11.**
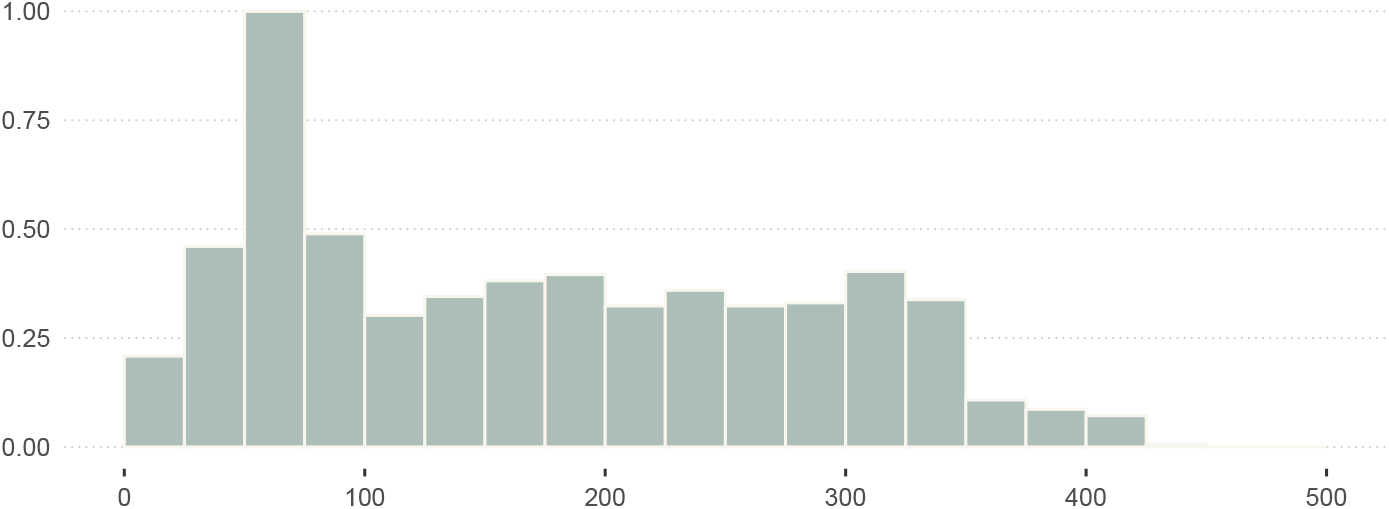
Empirical distribution of sampling effort (hours) across all dyads.

**Figure S12.**
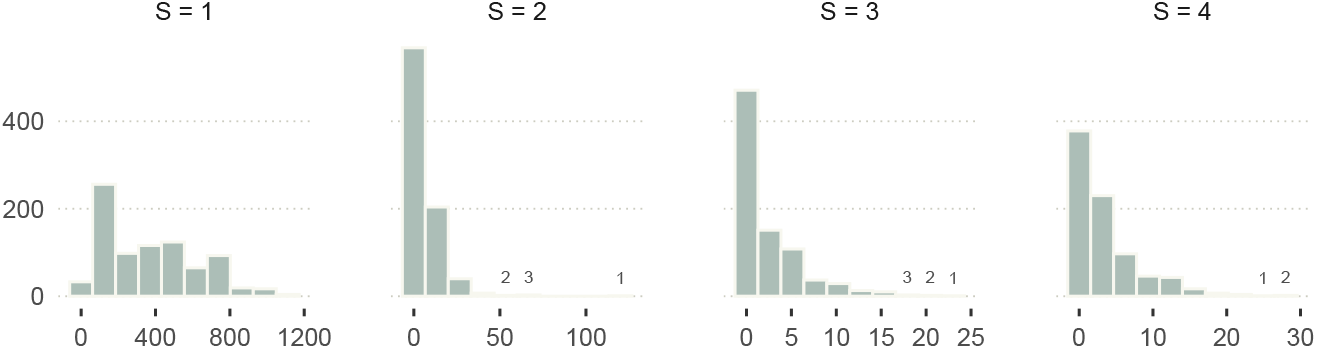
Number of observations j per dyad and per state k (empirical data set).

**Figure S13.**
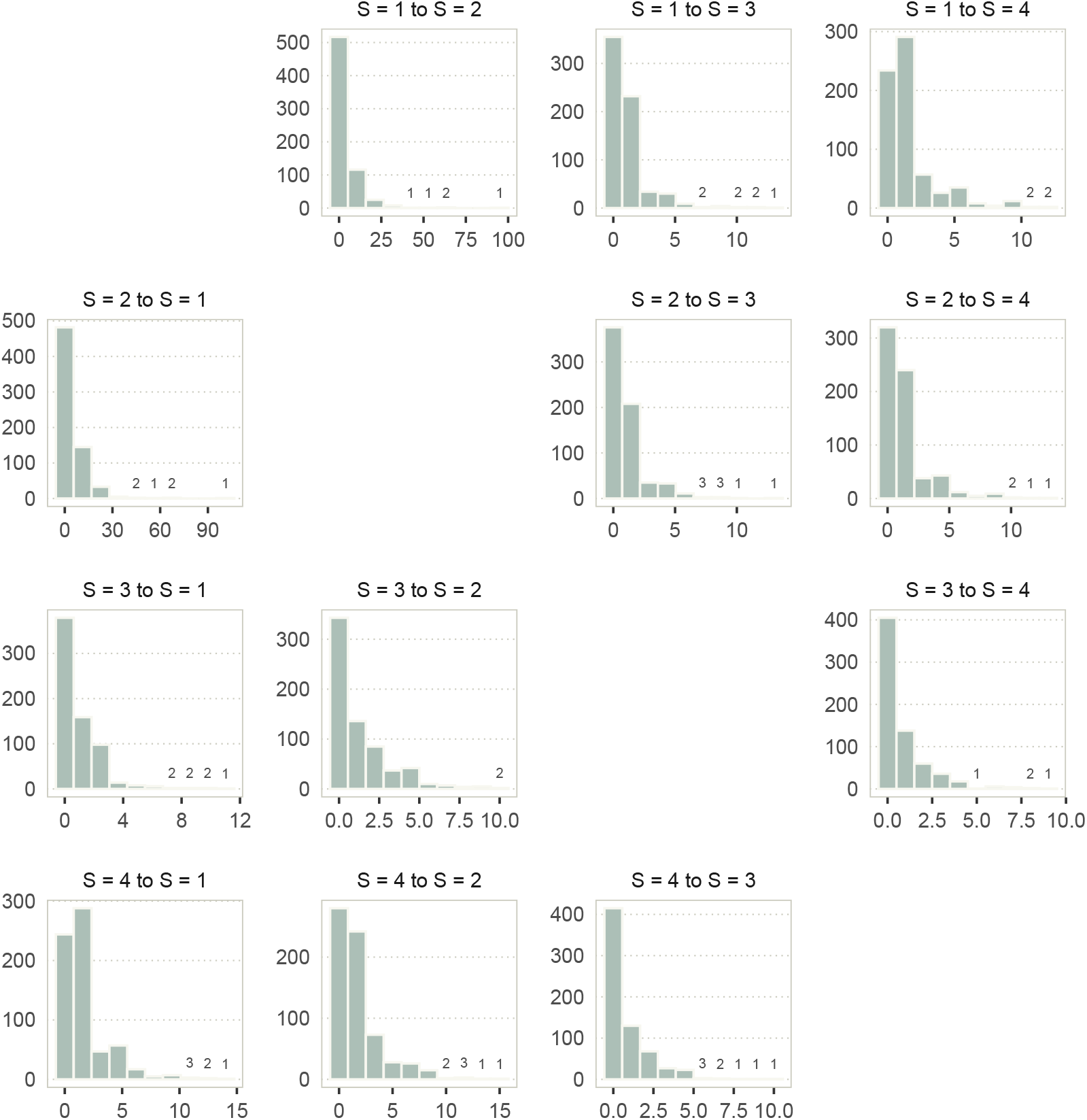
Number of observations transitions per dyad and per combination of current state k and future state l (empirical data set).

### S1.4. Empirical results

**Figure S14.**
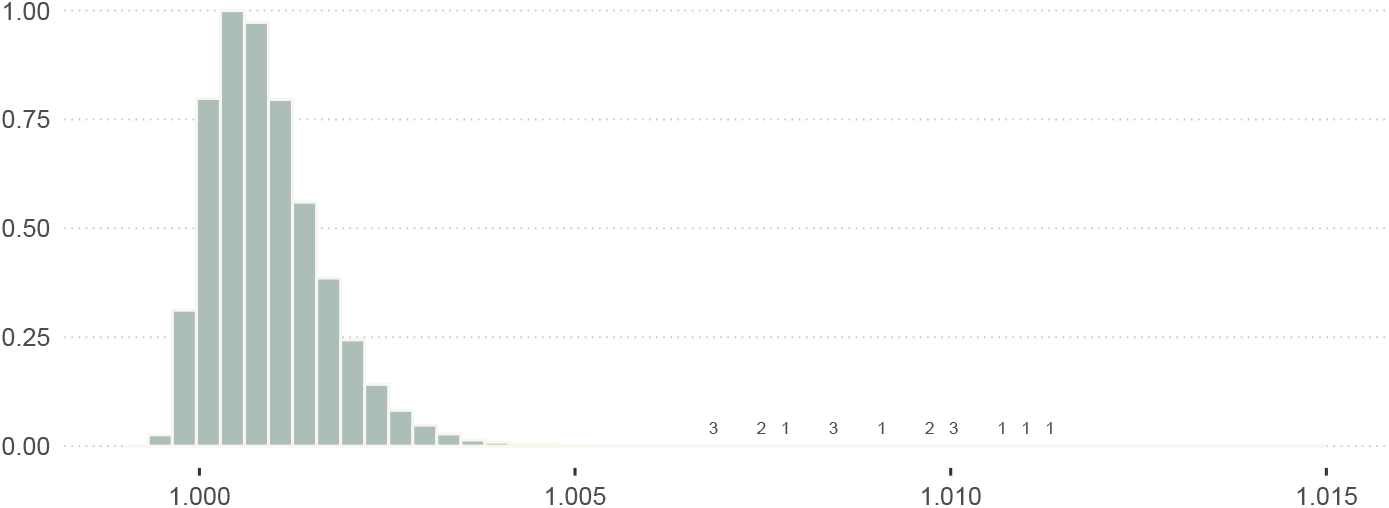
Distribution of 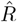 across all parameters of the core statistical model fitted to the empirical data (Vehtari et al., 2021).

**Figure S15.**
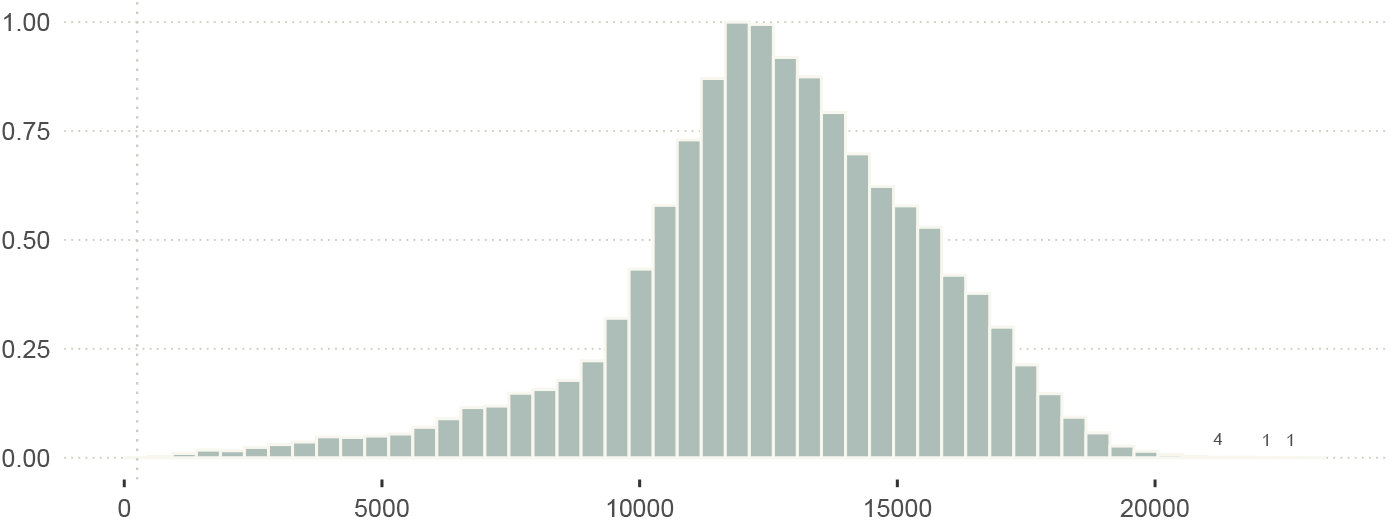
Distribution of the effective number of MCMC samples (bulk), across all parameters of the core statistical model fitted to the empirical data. The vertical line marks 250.

**Figure S16.**
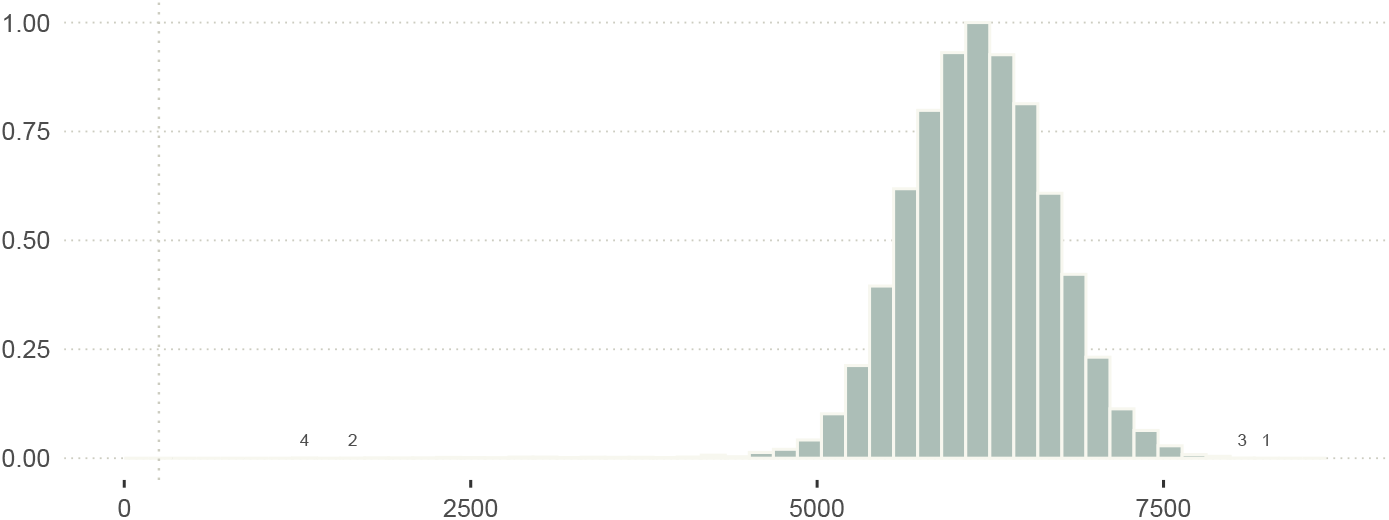
Distribution of the effective number of MCMC samples (tail), across all parameters of the core statistical model fitted to the empirical data. The vertical line marks 250.

#### S1.4.1. MCMC diagnostics

## S2. Extended models

### S2.1. Validation

**Figure S17.**
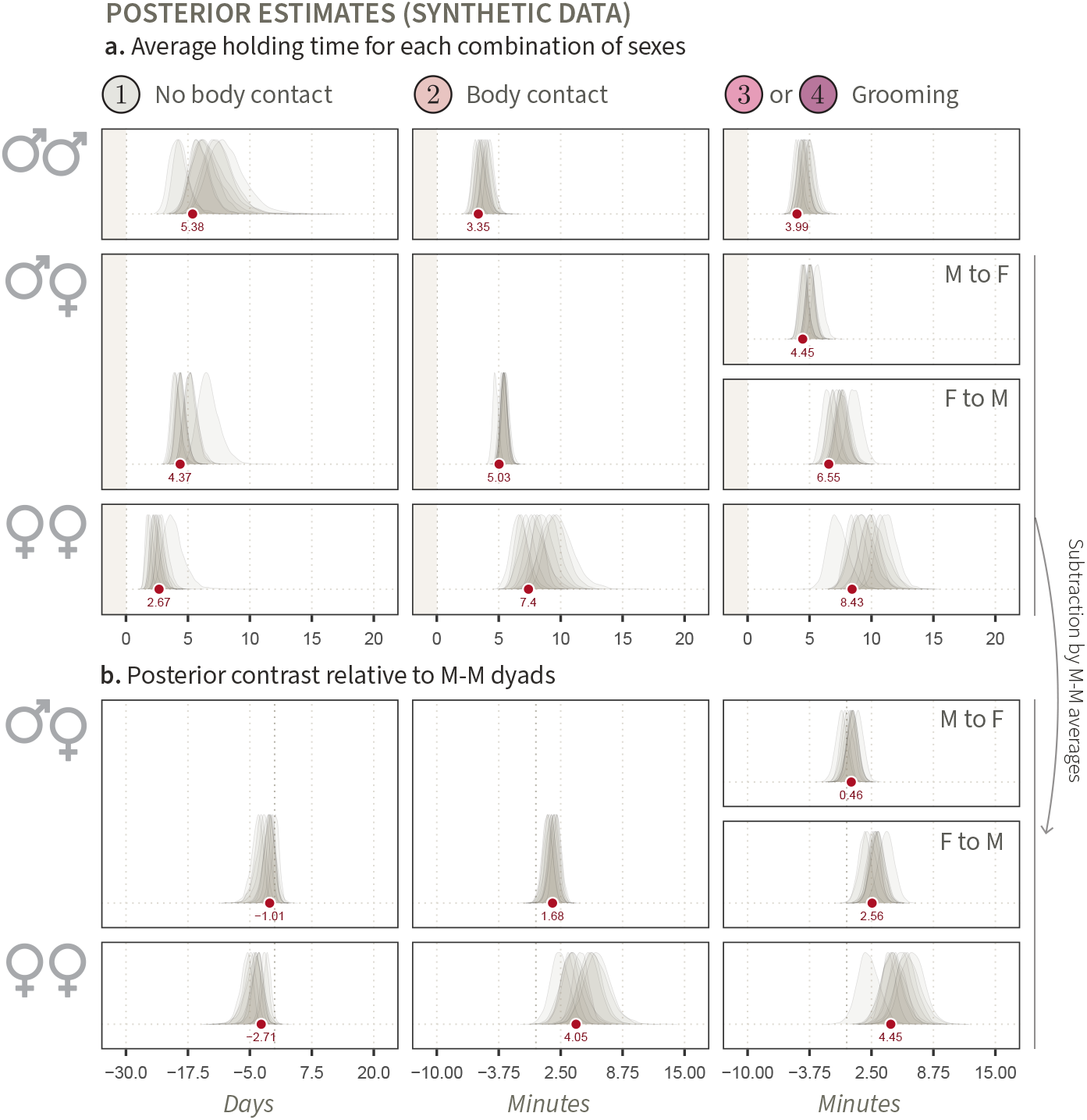
Posterior estimates of expected holding times and transition probabilities, averaged across all dyads (synthetic data) The figure is identical to Figure 11, except that we are here showing the estimates for synthetic data (10 iterations), which we compare to true values from the generative model (in red).

**Figure S18.**
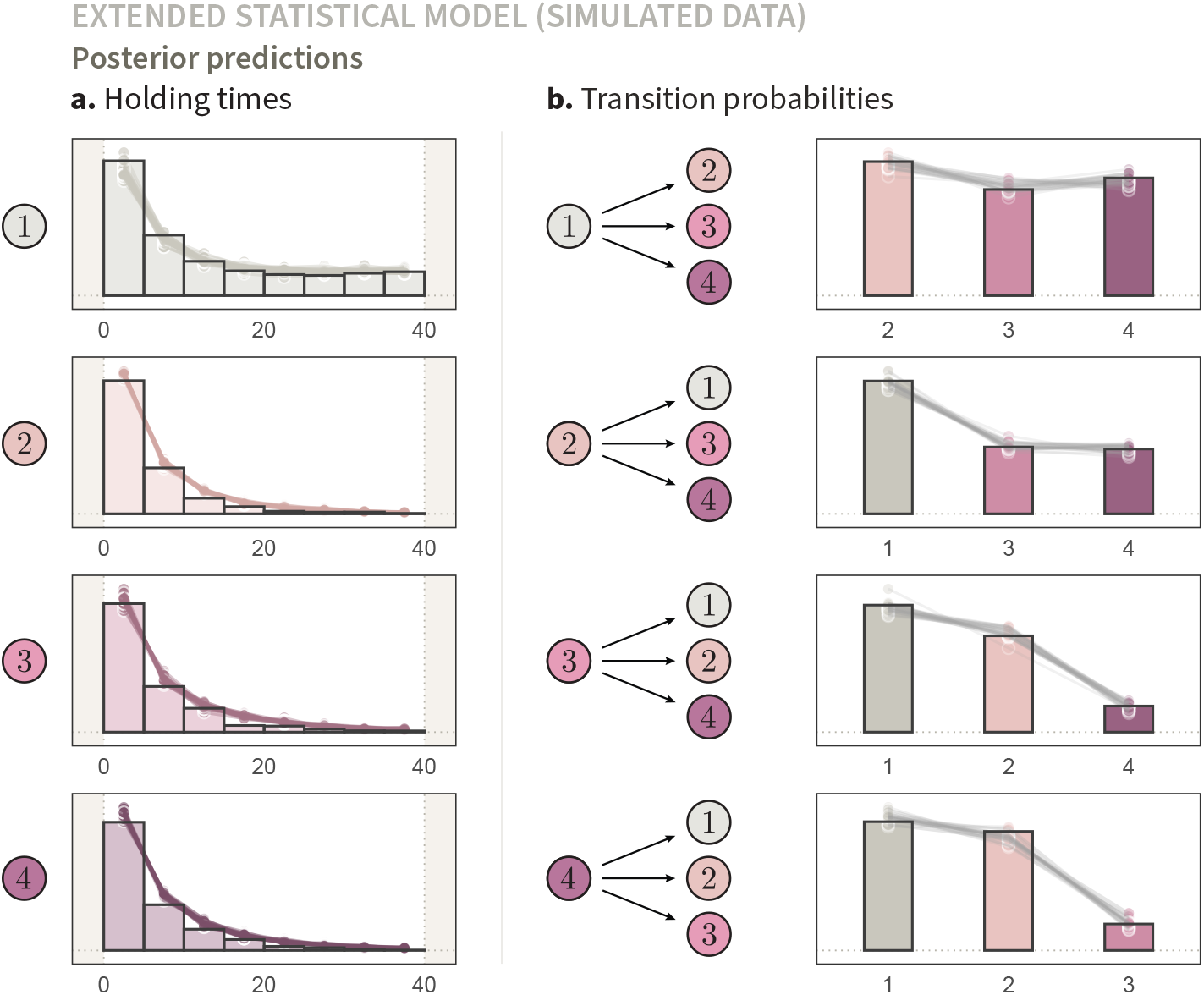
Posterior predictions of non-censored holding times and state transitions (synthetic data). This figure is similar to Figure 6, but the predictions shown here are generated from our extended model.

### S2.2. Empirical results

#### S2.2.1. MCMC diagnostics

**Figure S19.**
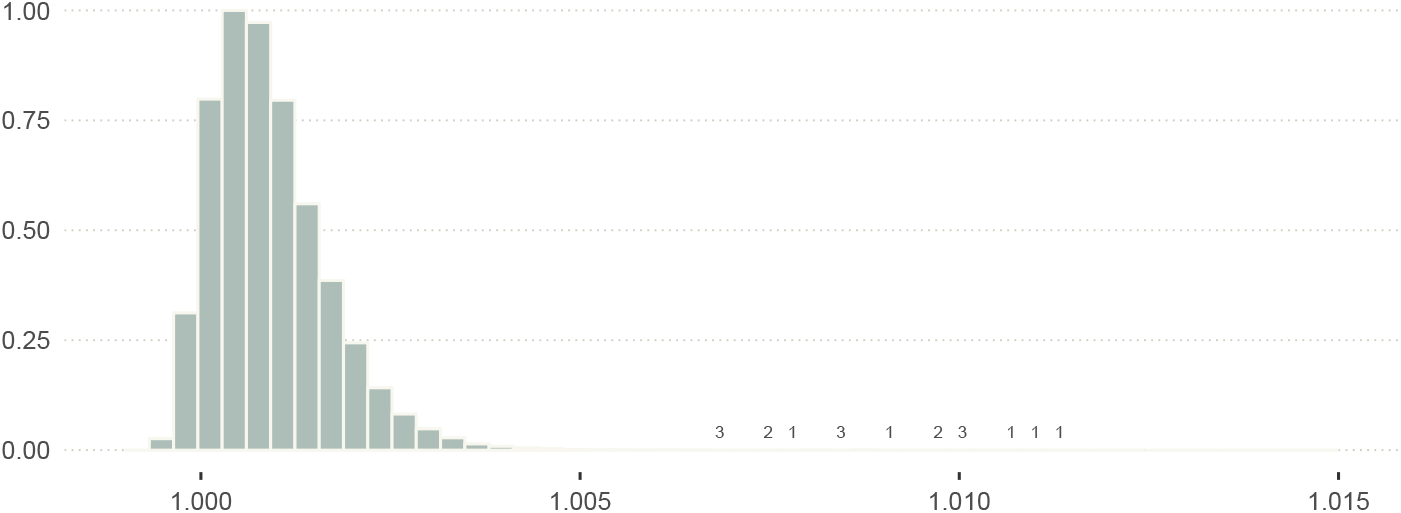
Distribution of 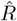 across all parameters of the extended statistical modelfittedto the empirical data (Vehtari et al., 2021).

**Figure S20.**
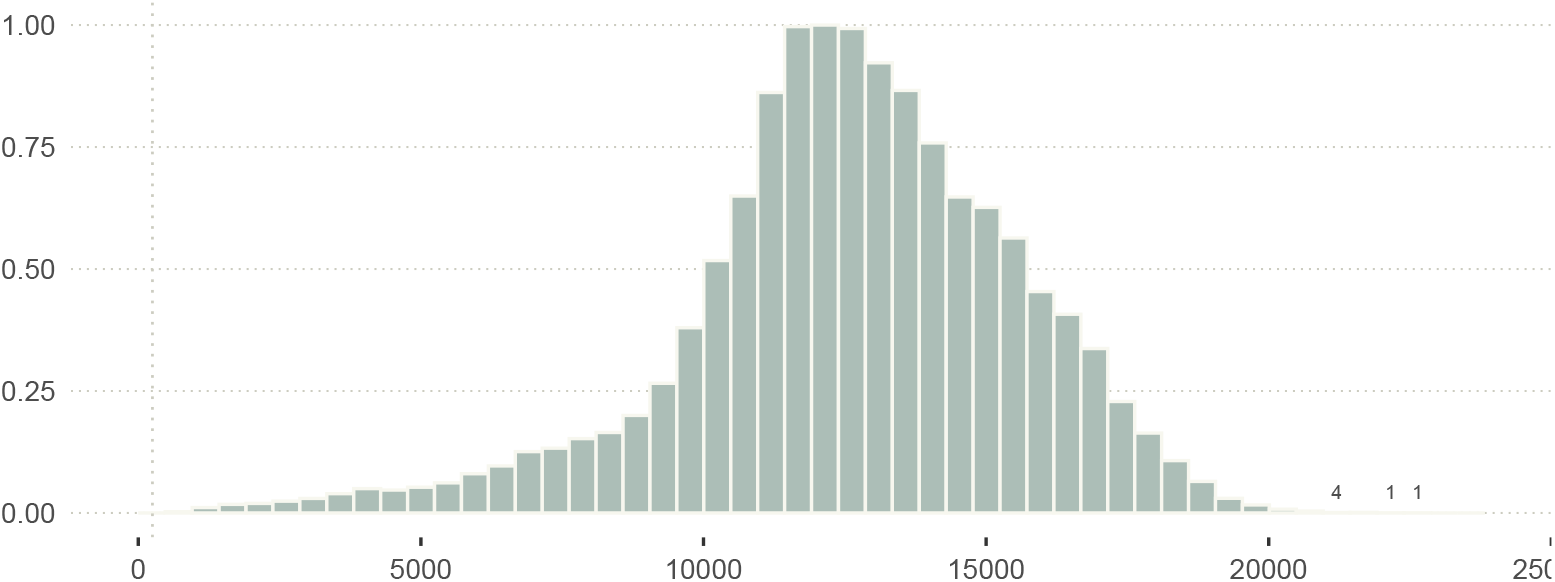
Distribution of the effective number of MCMC samples (bulk), across all parameters of the extended statistical model fitted to the empirical data. The vertical line marks 250.

**Figure S21.**
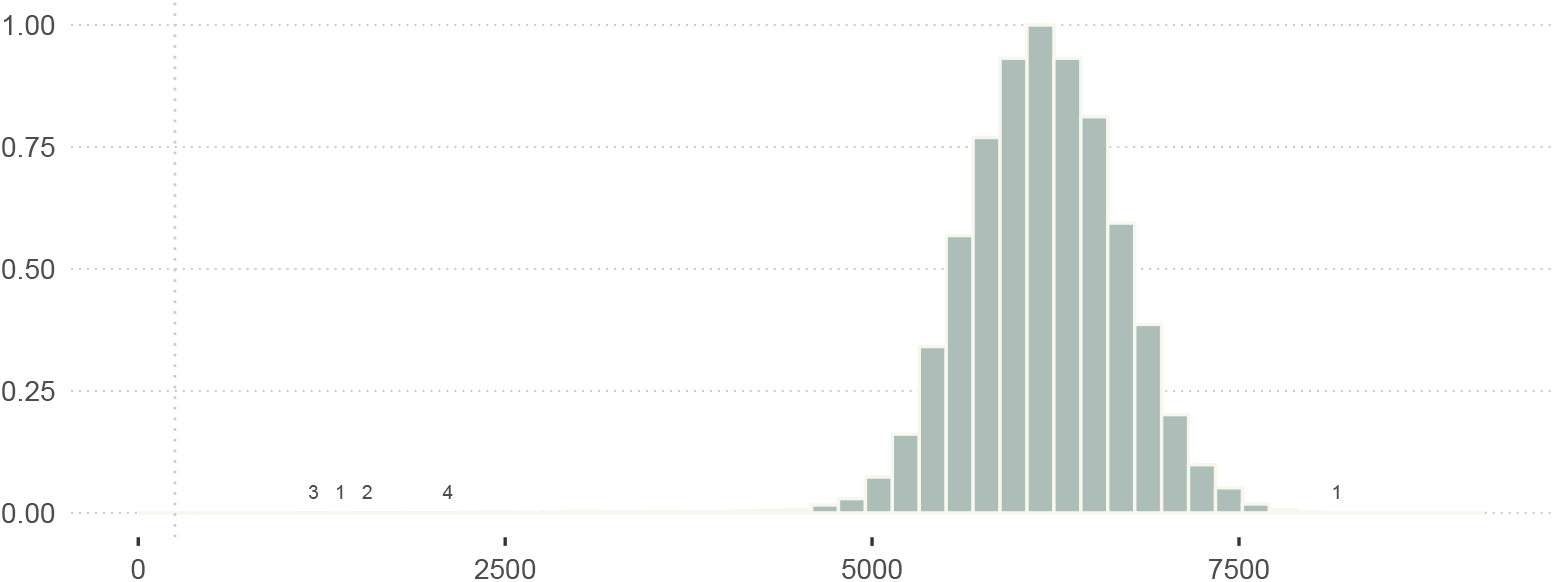
Distribution of the effective numberof MCMCsamples(tail), across all parameters of the extended statistical model fitted to the empirical data. The vertical line marks 250.

## Footnotes

1 In particular, the exponential distribution is *memoryless*. This property is defined as: 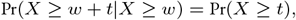 Where *X* is a random draw from the distribution, *w* is the time already spent waiting, and *t* is a duration greater or equal to 0 (Blitzstein & Hwang, 2019). This expression essentially states that the probability of waiting for a duration *t* does not depend on whether one has already waited for some time *w*. This property will prove very convenient, as we will see further below. Specifically, when designing a statistical model to recover the parameters *θ* from fragments of dyadic behavioural states, we will only need to consider whether the *ending times* of these states were observed, ignoring whether the starting times are known (see section 2.2).

2 Posterior estimates that are slightly biased across dyads (*e.g*., transition from *k* = 3 to *l* = 4 in Figure 5**a**) in small samples are not necessarily problematic. In fact, multilevel models—which are inherently biased—are often preferable to non-biased alternatives because they reduce overfitting. For a discussion on the bias-variance trade-off, see Greenland (2000).

